# Asymmetrical dose-responses shape the evolutionary trade-off between antifungal resistance and nutrient use

**DOI:** 10.1101/2021.11.29.469899

**Authors:** Philippe C Després, Angel F Cisneros, Emilie MM Alexander, Ria Sonigara, Cynthia Gagné-Thivierge, Alexandre K Dubé, Christian R Landry

**Affiliations:** Département de Biochimie, de Microbiologie et de Bio-informatique, Faculté des Sciences et de Génie, Université Laval, G1V 0A6, Canada; Institut de Biologie Intégrative et des Systèmes, Université Laval, G1V 0A6, Canada; PROTEO, Le regroupement québécois de recherche sur la fonction, l’ingénierie et les applications des protéines, Université Laval, G1V 0A6, Canada; Centre de Recherche sur les Données Massives, Université Laval, G1V 0A6, Canada; Département de Biologie, Faculté des Sciences et de Génie, Université Laval, G1V 0A6, Canada

## Abstract

Antimicrobial resistance is an emerging threat for public health. The success of resistance mutations depends on the trade-off between the benefits and costs they incur. This trade-off is largely unknown and uncharacterized for antifungals. Here, we systematically measure the effect of all amino acid substitutions in the yeast cytosine deaminase Fcy1, the target of the antifungal 5-FC (flucytosine). We identify over 900 missense mutations granting resistance to 5-FC, a large fraction of which appear to act through destabilisation of the protein. The relationship between 5-FC resistance and growth sustained by cytosine deamination is characterized by a sharp trade-off, such that small gains in resistance universally lead to large losses in canonical enzyme function. We show that this steep relationship can be explained by differences in the dose-response functions of 5-FC and cytosine. Finally, we observe the same trade-off shape for the ortholog of *FCY1* in *Cryptoccocus neoformans*, a human pathogen. Our results provide a powerful resource and platform for interpreting drug target variants in fungal pathogens as well as unprecedented insights into resistance-function trade-offs.

## Introduction

Pathogenic fungi are an rising threat to human health, agriculture and wildlife^1^. Therapeutic options remain limited and resistance to all antifungals classes has been observed in multiple species^2^. Very little is known about the functional landscape of antifungal targets: the genotype of a strain is rarely enough to infer its phenotype. Recent efforts to catalog mutations associated with resistance have yielded a handful of mutations per species for a few targets, mostly derived from small scale studies and not always experimentally validated^3^. More systematic efforts are being deployed to catalog resistance mutations in a comprehensive manner^4, 5^, but information remains sparse for most targets.

The benefit of resistance to a given drug is not the only factor that affects the long-term outcome of a mutation: the associated cost is also a key parameter^6^. For instance, a mutation in an essential protein that confers resistance but leads to a decrease in function would rapidly be lost upon treatment interruption. For example, resistance to the antifungal nystatin has been shown to have significant fitness costs in stressful environments^7^, which means they would actively be selected against when nystatin concentration drops. In *Candida albicans*, aneuploidies are associated with antifungal resistance but are deleterious in many environments^8^. Both a better inventory of the mutations granting resistance and the trade-offs associated with them are key to improve our ability to predict or interpret clinically relevant genotypes in complex environments. Ideally, we would need approaches that allow us to measure this type of trade-off in a systematic and comprehensive manner for resistance mutations. Beyond their impact on antimicrobial resistance, these relationships can play large roles in shaping which evolutionary trajectories are available in the face of harsh selective pressures, from adaptation to new environments in the wild to cancer evolution.

Most antifungals used in the clinic are drugs that bind their targets and disrupt their activity or impair their structural integrity^9, 10^. For instance, azoles bind to the active sites of their targets and inhibit their function. Resistance mutations therefore tend to occur in the substrate binding pocket^11^. An exception to this is 5-fluorocytosine (5-FC, also known as flucytosine), which is administered as a pro-drug that is deaminated into the cytotoxic nucleotide analog 5-fluorouracil (5-FU) inside fungal cells. The accumulation of 5-FU leads to defects in mRNA synthesis and protein translation as well as DNA damage through thymine starvation^12^, resulting in dose-dependent fungistatic or fungicidal activity. As one of the oldest antifungals, 5-FC is on the WHO list of essential medicines and is often used to treat cryptococcal meningitis in combination with amphotericin B^13^.

In yeast, the cytosine deaminase Fcy1 (Fca1p in *Candida albicans*) forms a homodimeric enzyme that converts 5-FC to 5-FU, and loss of function of this protein leads to 5-FC resistance^14^. Fcy1 is widely conserved throughout fungi, and has a distinct fold compared to the *Escherichia coli* cytosine deaminase^15^. As vertebrates lack a *FCY1* ortholog, 5-FC can be administered as a prodrug that will be selectively converted by fungal cells into 5-FU. *FCY1* deletion was shown to confer full 5-FC resistance to *S. cerevisiae*^14^, with little effect on fitness in multiple lab conditions^16, 17^. Gene loss of function, for instance through premature stop codons, thus appears to be an easily accessible mechanism of resistance. However, since Fcy1 is a dispensable enzyme in standard growth media, it is unclear if 5-FC resistance through loss of function leads to any trade-off since enzyme function is generally decoupled from growth. In contrast, other antifungal targets, such as *ERG11* or *FKS1*, are considered essential for growth and thus for virulence in some pathogens^18^. Interestingly, *FCY1* can be conditionally essential in yeast strains where the *de novo* uracil synthesis pathway has been disrupted. In this genetic background, cells must rely on the pyrimidine salvage pathway, where Fcy1 deaminates cytosine into uracil, complementing the auxotrophy. For example, in the lab strain BY4743, *FCY1* masks the loss of function of *URA3* in minimal media supplemented with cytosine^19^. By changing environmental conditions, the interplay between resistance and Fcy1 activity can thus serve as a model for a common scenario where an essential gene imposes strict constraints on available evolutionary trajectories because of cell metabolism requirements.

An important knowledge gap regarding the emergence of resistance is whether complete loss of function of Fcy1 is necessary or if partial loss of activity is sufficient, which would dramatically increase the mutational target size. Because the natural substrate, cytosine, and the toxic analog 5-FC have very similar structures, it is likely that a loss in the ability to process 5-FC will also lead to a loss in the ability to metabolize cytosine. Recent work examining the activity of enzyme variants on different substrates at high throughput have shown that the level of correlation between mutation effects can vary depending on substrate pairs and the enzyme itself^20, 21^. As such, it is difficult to predict which trade-off scenario applies in the case of Fcy1. If it is weak, there may be mutations that lead to resistance while maintaining the ability of the cell to utilize cytosine. These mutations could in principle lead to resistance without seriously affecting fitness in the absence of the drug. The relationship could be linear, in which case a 50% increase in resistance would lead to a 50% decrease in cytosine use. At the other end of the spectrum, there may be a strong trade-off, which means that all mutations leading to 5-FC resistance impede cytosine use. The emergence of loss of function alleles would entirely be at the expense of Fcy1 activity. For some types of coding sequence mutations, such as frameshifts or premature stop codons, loss of function can be easy to predict. However, significant challenges and uncertainty remain when trying to predict the effect of missense mutations distal from the active site. Possible consequences could include allosteric effects on enzymatic function, or simply destabilisation of the protein structure or Fcy1 dimer assembly. These perturbations would lower the effective protein concentration, potentially leading to loss of function phenotypes. Sensitivity to destabilisation would then be another key parameter in determining the mutational target size for resistance mutations.

Recent advances in genome editing and DNA sequencing have greatly increased the throughput at which protein variants can be assayed, allowing to draw a high-resolution map of fitness effects and potential trade-offs. Here, we use Deep Mutational Scanning (DMS) to explore the functional landscape of *FCY1* at its endogenous locus in the context of 5-FC resistance and cytosine deamination. We systematically assess protein variants to determine which mutations lead to drug resistance and explore which protein properties are linked to it. We examine how resistance mutations are distributed on the enzyme and we characterize the trade-off between resistance and cytosine use. We then show that this trade-off is conserved in the *FCY1* ortholog of *C. neoformans*.

## Results

### Deep mutational scanning of Fcy1

We used the megaprimer method^22^ to generate all possible single codon changes along the FCY1 coding sequence (see Methods). This approach results in the full set of premature stop codons, silent mutations and single position missense mutants (which result in protein variants). We integrated the mutations at the native *FCY1* locus by using a “bait and switch” genome editing strategy^23, 24^. Since our approach requires direct sequencing of the coding sequence and because *FCY1* is too large to sequence with a single read pair using high-throughput sequencing, we split our mutants into three pools each covering partly overlapping sections of the coding sequence (Figure 1A). We used two different media compositions: media with 5-FC to select for resistance, and media without uracil but enriched with cytosine (cytosine media) to select for cytosine deamination into uracil. In this cytosine media, the reaction is essential for growth because it complements the uracil auxotrophy of the strain. Using this approach, growth rate can be used as a reporter for Fcy1 activity for both enzymatic reactions in pooled competition assays (Figure 1B). Comparing the results in 5-FC and cytosine media allows us to measure the strength of the trade-off between the two phenotypes across the mutational landscape for all single mutations (Figure 1C). To define the conditions for the pool competitions, we identified which 5-FC or cytosine concentrations reduce growth rate to ∼50% of the maximum (also known as IC_50_, Figure S1). This corresponds to roughly 12 μM and 84 μ for 5-FC and cytosine, respectively.

**Figure 1:**
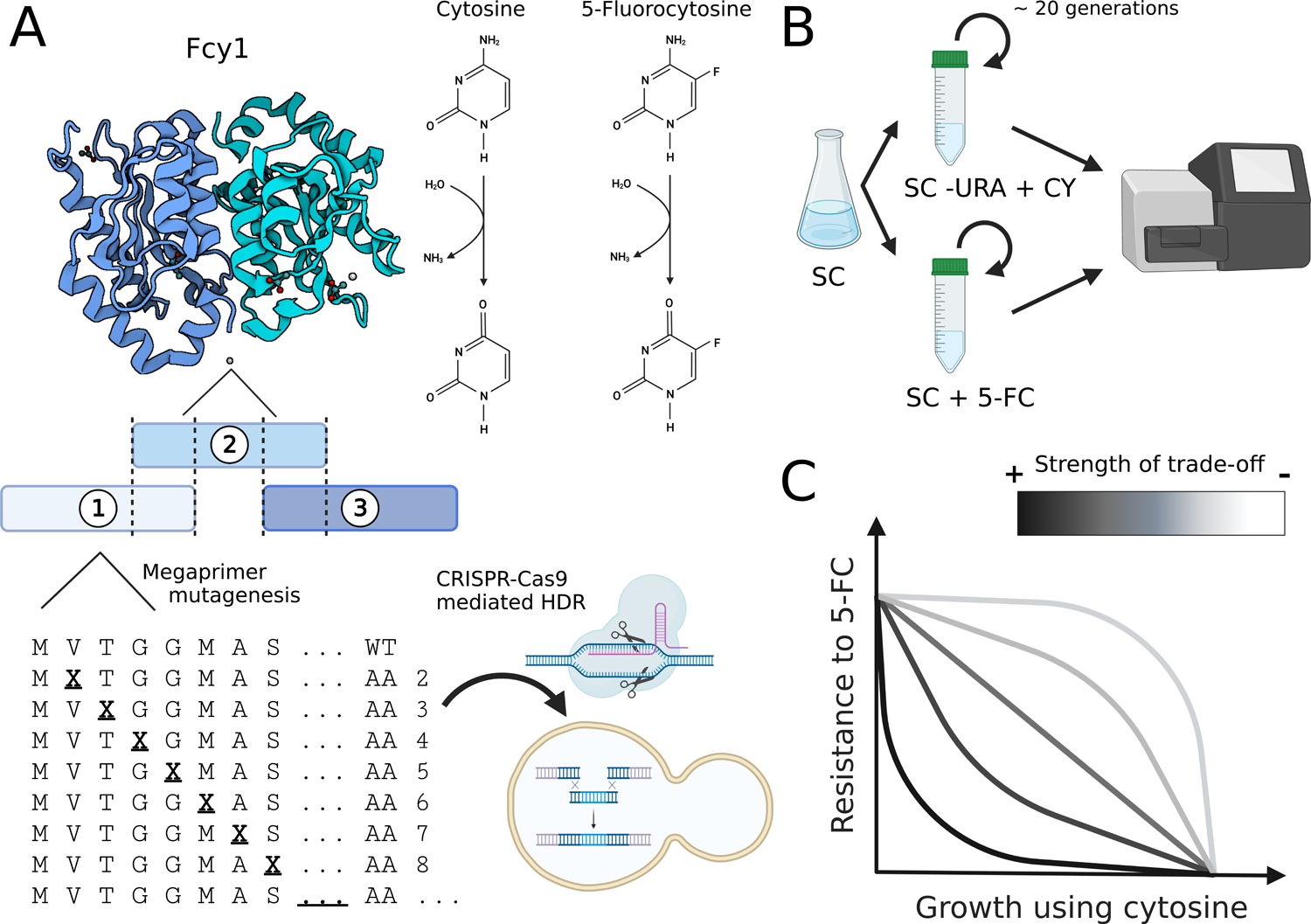
Fcy1 as a model for resistance-function trade-offs. **A)** Structure of the Fcy1 homodimer (pdb: 1p6o^15^) and two reactions it catalyzes with opposed effects on fitness. To explore *FCY1*’s functional landscape, each codon was mutated one by one to all other possibilities to generate a comprehensive pool of mutants. The mutants were split into three partly overlapping fragments, and studied in three pools: pool 1 covers from position 2-67, pool 2 from 49-110 and pool 3 from 93 to 158. The resulting libraries were then integrated at the native *S. cerevisiae* locus using CRISPR-Cas9 genome editing, therefore preserving its endogenous transcriptional regulation. **B)** Workflow for the pooled mutants growth assays. The FCY1 mutant libraries were expanded by a round of culture in non-selective media before being split and transferred into 5-FC and cytosine media where mutants competed for around ∼20 generations. Variant abundance before and after competition was measured by high-throughput sequencing. **C)** Trade-off of different intensities between resistance to 5-FC and growth using cytosine. In a weak trade-off scenario (light grey), resistance mutations that preserve the canonical enzyme function exist. Resistance can coexist with cytosine use. A strong trade-off (dark) instead implies that mutations that lead to resistance necessarily lead to loss the of cytosine deamination activity.

During preliminary experiments, we observed growth patterns on solid media suggesting that *fcy1*Δ strains could grow on cytosine media if wild-type strains were also present on the plates (Figure S2). We hypothesized this could be due to cross-feeding between wild-type and mutant cells, which could potentially impact selection strength during the pooled competition assay. Previous work has indeed shown that the yeast plasma membrane can leak uracil in the surrounding media^25^. The main contributor to uracil uptake in yeast is the transporter *FUR4*^26^, the deletion of which is lethal when combined with a non functional *URA3* allele in standard media^27^. As uracil auxotrophy is essential to one of our selection conditions, we could not disrupt *FUR4* to prevent cross-feeding. Instead, we quantified the impact of potential uracil leakage on selection intensity in liquid cultures. Using a cytometry based competition assay, we measured the effect of different cytosine concentrations and population compositions on the selection coefficient of a *FCY1* deletion. We found that high cytosine concentrations and a higher initial proportion of wild-type cells tended to lower selection strength against *fcy1*Δ (Figure S3). Reassuringly, the selection coefficient remained above 0.3 at the cytosine IC_50_ (84 µM), even in populations with as much as 90% of wild-type cells. This is well above the threshold required to detect fitness effects in a high-throughput sequencing based assay^28^.

We extracted genomic DNA and amplified the appropriate *FCY1* region for each pool and condition, and measured changes in allele frequencies between the start and the end of the competition. Variant libraries before selection had high coverage, with over 99% of single codon mutants with 10 or more read counts (Figure S4). Synonymous codons were used as internal replicates for individual amino acids: log_2_ fold-change for each protein mutant was calculated as the median of the codon level fold-changes. Spearman’s correlation coefficient for replicate pairs ranged between 0.814 and 0.945 (Figure S5). To adjust for the differences between the different mutant pools (each covering a third of the coding sequence), we harmonized the log_2_ fold-change to pool 2 using the variants in the overlapping regions of the pools. The log_2_ fold-changes of mutants in these two regions (codons 49 to 67 and 93 to 110 respectively) were well correlated between pools (Figure S6 and S7A-F). We then used the distribution of log_2_ fold changes from the synonymous and nonsense mutants to put both 5-FC and cytosine conditions on the same scale to obtain the DMS score (Figure S7G and H). This allowed us to obtain a full-length functional landscape of Fcy1 that covered 99.5% (2968/2983) of possible missense variants in both selection conditions (Figure 2A and B).

**Figure 2:**
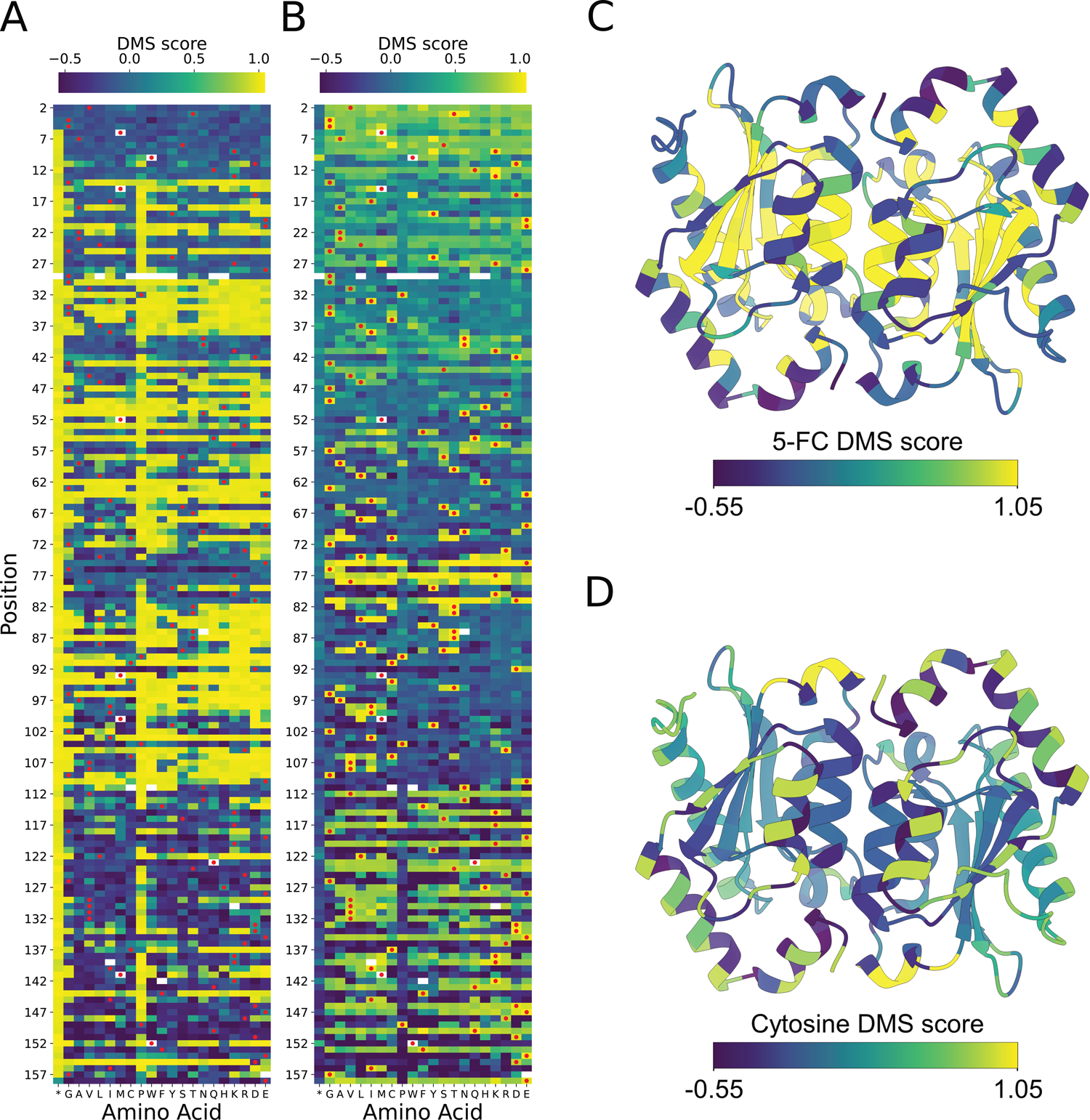
The functional landscape of Fcy1 identifies all resistance mutations. **A)** Fcy1 functional landscape in 5-FC media and **B)** in cytosine media. Red dots represent the wild-type amino acid at each position. The score represents the DMS mutant log_2_ fold-change, scaled by the median of scores of synonymous and nonsense mutations (see methods). The leftmost column on both heatmaps represents the scores for stop-codons (*). A positive DMS score indicates faster growth. **C)** Position-wise median 5-FC and **D)** cytosine DMS scores of missense mutants, mapped onto the Fcy1 homodimer (PDB: 1p6o^15^). The color scaling across the four panels is based on the 2th and 98th percentile of the 5-FC scores distribution.

The distributions of DMS scores are bimodal, with the two modes corresponding to WT sequences and premature stop codons (Figure S8A-C). Stop codon mutants can have different effects based on their position in the coding sequence, with generally weaker effects near the end of the coding sequence^29^. We observed that only the last amino acid of the C-terminus was partially dispensable. Surprisingly, stop codons that occured before the methionine in position 6 did not grant 5-FC resistance, suggesting that Fcy1 translation from an alternative start site, Met 6, remains efficient enough to confer sensitivity. We excluded stop codon mutants from positions 2-5 and 158 from downstream analyses.

Across the mutant library, 32% (940/2968) of missense mutants had 5-FC DMS scores that fell in the range observed for nonsense mutants (5-FC DMS score ≥ 0.921, 1^st^ percentile of nonsense scores). Of the resistant mutants, 284 can be recapitulated by a single nucleotide change in the *S. cerevisiae FCY1* coding sequence, representing a large set of easily accessible mutations leading to resistance. The mutational target size of resistance is therefore much larger than stop codon mutations. For mutants in pool 2, which cover most of the substrate binding and catalytic residues of Fcy1, we also performed the experiments in an additional condition to investigate whether some mutations could improve cytosine specificity. To test this, we used cytosine media with 24 μ 5-FC and 113 μM cytosine, in which wild-type cells grew at a similar rate as in the two other conditions. Mutant scores were highly correlated between the cytosine and 5-FC+cytosine condition (Figure S7D), with the only difference being better signal in the latter, probably due to more stringent selection. Based on this comparison, we did not identify any mutants in our library for which we could infer increased specificity for cytosine over 5-FC compared to the wild-type. Having catalogued which amino acid changes conferred resistance or maintained Fcy1 activity, we next sought to better understand the mechanisms behind mutational effects.

### 5-FC Resistance mutations are distributed across the protein and are amino acids rarely observed at orthologous sites

Mapping the DMS scores on the Fcy1 structure (Figure 2C and 2D), we found that the active site and the residues in close contact with the substrate were extremely sensitive to mutations. Our results corroborate what has been observed in small-scale studies on Fcy1 biochemistry. For example, Glu 64 is essential for catalytic activity^30^, and we did not observe any variant at this position that maintained 5-FC sensitivity or growth in cytosine. Focusing on functional sites known to be involved in the catalytic cycle, we found that the core components of Fcy1 were strongly intolerant to mutations with respect to cytosine use (Figure 3A). Surprisingly, most resistance mutations do not reside in the active site of the enzyme, suggesting that the enzyme is very sensitive to amino acid changes, either by allosterically affecting the enzymatic activity or perturbing the stability of the complex.

**Figure 3:**
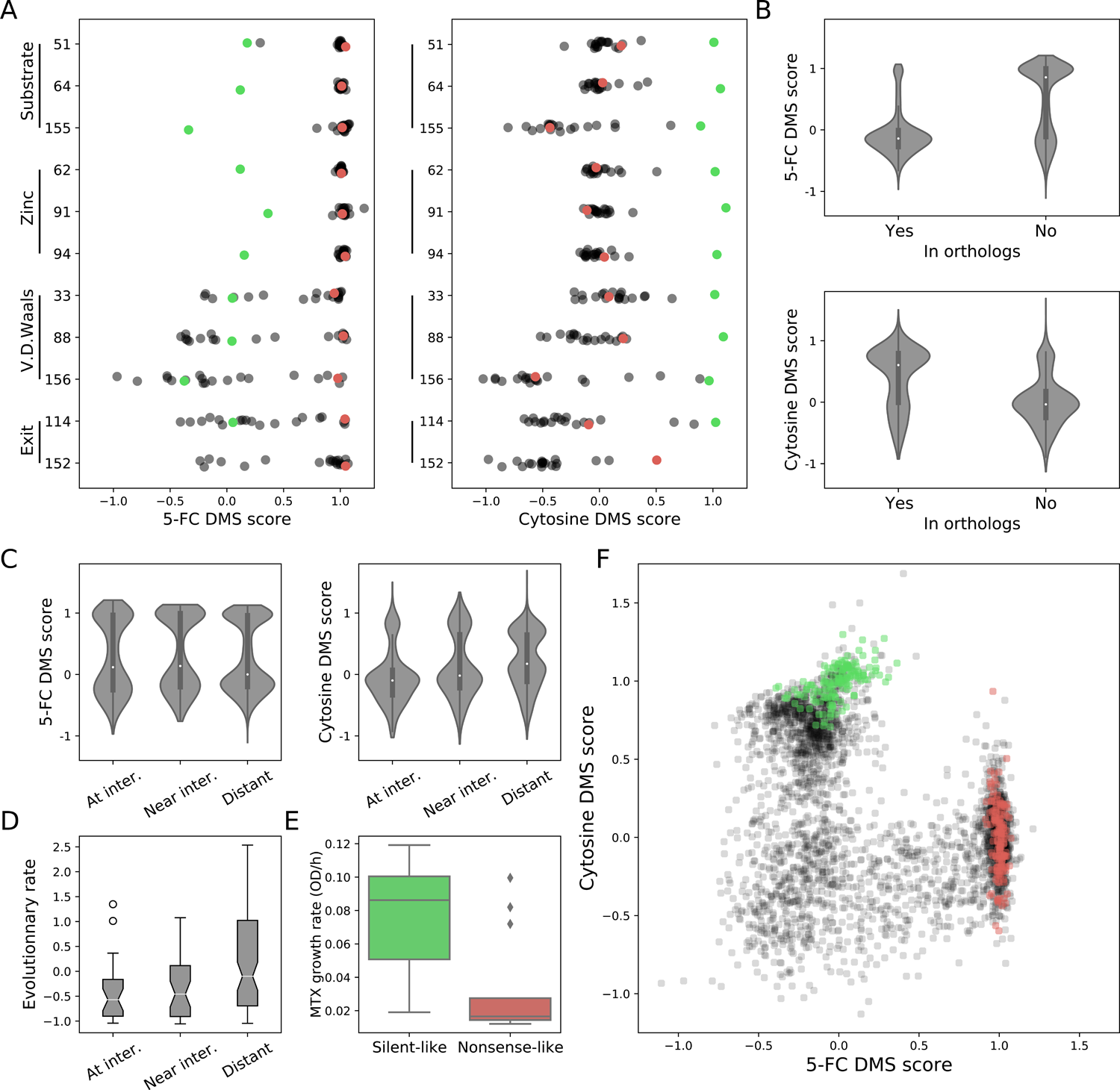
Fcy1 amino acid substitutions can lead to 5-FC resistance through multiple mechanisms. **A)** 5-FC and cytosine DMS scores of mutations at the substrate binding and catalytic residues (substrate), zinc cofactor binding residues (Zinc), residues involved in binding stabilization through Van der Waals interactions (V.d.Waals) and residues involved in guiding the exit of the product (exit). The score for the silent and nonsense variants for each position are shown in green and red respectively. **B)** 5-FC and cytosine DMS scores of missense variants as a function of their presence in orthologous sequences (n=1,078 observed, n=1,890 unobserved). The difference between medians was tested using Mood’s median test: for 5-FC scores, p=8.12×10^-1^^34^, and for cytosine, p=2.68×10^-69^. **C)** 5-FC and cytosine DMS score distributions for residues as a function of their distance to the dimer interface. n=531 variants at the interface, n=722 variants near the interface, and n=1,715 distant variants. **D)** Evolutionary rates of residues at, near or distant to the dimer interface. **E)** The stability of the Fcy1-Fcy1 complex measured by DHFR-PCA for varinats clustering with silent mutations is higher than for those clustering with nonsense mutations (p=0.006, Mood’s median test, n=10 silent-like mutants, n=18 nonsense-like mutants). In DHFR-PCA assay, growth in methotrexate (MTX) (y-axis) is coupled with the amount of complex formed. **F)** Two dimensional representation of 5-FC and cytosine DMS scores for all variants. Silent mutants are shown in green, nonsense in red.

To better understand the constraints linked with maintaining Fcy1 function, we assembled a set of 215 orthologs (Figure S9) and hypothesized that conserved sites would be more sensitive to mutations. On average, amino acids from *S. cerevisiae* Fcy1 were present in 60% of orthologous sequences at the homologous positions, but the distribution of occupancy was bimodal (Figure S10A). As much as 18.5% of residues were observed in more than 95% of sequences, indicating strong structural and functional constraints maintaining residues for millions of years of evolution. Sites not part of the active site or involved in the dimer interface accounted for 41% of highly conserved amino acids, highlighting the importance of distal sites for Fcy1 function.

We directly investigated the effect of missense mutants in our library for which an equivalent wild-type maino acid could be found in orthologs. Variants observed in orthologous sequences had much lower 5-FC scores and higher cytosine scores compared to those not observed (Figure 3B). Again the distributions are bimodal, which means for instance that some amino acids are compatible with the WT phenotype in *S. cerevisiae* but are not seen in orthologs. Since the number of orthologs is limited and does not cover the entire sequence space compatible with this enzyme’s function, this is not unexpected. There are few cases where amino acids present in orthologs lead to resistance to 5-FC when introduced in *S. cerevisiae* but those mainly concern amino acids present in the most distant species, which means that they could have been compensated for by other amino acid substitutions in the sequence^31^. Overall, only 56 amino acid changes observed in orthologs conferred resistance, with most of them being unique to a specific ortholog (median occupancy=0.47%). These shared functional constraints suggest resistance to 5-FC could arise by similar mechanisms in *FCY1* orthologs.

### Protein complex destabilisation is a major mechanism of resistance

Many of the highly conserved residues that lead to resistance when mutated are away from the active site. This is possibly due to the importance of structural integrity of the complex or instances of allosteric regulation of enzyme function. Using an *in silico* approach^32^, we predicted the effect on Fcy1 structure and dimer interface stability (ΔΔG) of all missense mutants (see methods). The predicted effects on stability of amino acid mutants were strongly correlated with their DMS scores, being positively correlated with 5-FC resistance (ρ=0.523, p=1.21×10^-206^, n=2,968) and negatively correlated with cytosine DMS scores (ρ=-0.422, p=1.1×10^-127^, n=2,968). Protein destabilisation is therefore a key determinant of 5-FC resistance.

The dimer interface also appears to play a role in maintaining Fcy1 function. While protein engineering efforts have investigated mutations predicted to stabilise the dimer^33^, whether dimerization is essential for activity remains unknown. The alpha-helix at the core of the interface is highly sensitive to mutations, suggesting that dimerization is important (Figure 2C and D). We found that mutants of residues in contact with the other subunit tended to have lower cytosine DMS scores compared to those near (within 6 Å of a residue from the other chain) or distant from the interface (Mood’s median test, p= 9.20 ×10^-6^ at interface vs near interface, p=9.15×10^-15^ near interface vs distant). Interestingly, we did not observe the same relationship for the 5-FC DMS scores, where only the difference in median score between mutants located near the interface and those distant was significant (Mood’s median test, p= 0.75 at interface vs near interface, p=0.018 near interface vs distant). Leveraging our set of orthologs, we calculated the evolutionary rate for each position in the alignment^34^ (Figure S10B), which revealed that both the core dimer interface and the interface rim evolved at a slower rate compared to other regions of the protein (Figure 3D). While homodimer formation appears to be essential to Fcy1 function, perturbing the interaction appears insufficient to grant resistance.

To validate the role of protein complex depletion on 5-FC resistance in the DMS data as well as our *in silico* prediction of destabilisation, we decided to investigate Fcy1 dimer formation *in vivo*. We used an assay that reports on the amount of protein complex formed in the cell. Dihydrofolate reductase protein complementation assay (DHFR-PCA) is an *in vivo* assay that allows quantitative measurement of a protein-protein interaction (PPI) by linking it to the reconstitution of a methotrexate-resistant DHFR^35^. Growth quantitatively varies as a function of both the abundance of the proteins interacting and PPI strength^35, 36^. This provides information on the abundance of the Fcy1 dimer in conditions in which *FCY1* is dispensable so as to not confound the PPI signal with the fitness of the mutants. Starting from a set of individually reconstructed mutants (see next section), we examined 54 Fcy1 variants to measure PPI abundance. We measured the interaction between a wild-type copy of Fcy1 and each mutant, which reflects the stability and ability of the WT-mutant complex to form.

Replicate control and methotrexate growth rates of the DHFR-PCA assays were well correlated between replicates (Figure S11A and B) and did not reveal any bias due to lower overall growth of some mutants (Figure S11C). As expected if destabilisation were causing resistance, variants that clustered with silent mutations in the DMS assays generally had detectable PPI. In contrast, those that clustered with stop codon mutants generally lost the interaction (Figure 3E, p=0.006, Mood’s median test). Importantly, PPI intensity was well correlated with FoldX predicted effects on protein structure stability (ρ = −0.475, p=0.0003, Figure S11D), as well as with variant growth in cytosine (ρ=0.375, p=0.009, Figure S11E) and negatively correlated with 5-FC resistance (ρ =-0.477, p=0.0005, Figure S11F). These results directly link altered protein complex abundance with mutant phenotypes, and validate the use of *in silico* tools to predict the effect of mutations on Fcy1 stability. On the other hand, there was no correlation between PPI intensity and *in silico* changes in PPI stability as predicted by FoldX (ρ=0.156 and p=0.259 for 54 mutants), limiting the relevance of such metrics for Fcy1. Based on these observations, structure or subunit destabilisation leading to lower protein complex abundance appears to be a frequent mechanism driving Fcy1 loss of function.

### A strong trade-off links 5-FC resistance and cytosine use

At first glance, 5-FC and cytosine DMS scores appear negatively correlated with each other. This is visible in Figure 2: sites that lead to resistance to 5-FC when mutated generally have reduced fitness in cytosine media. However, the scores are overall only weakly correlated (ρ=-0.23, p=5.5×10^-42^, n=3,272 variants). Directly examining the relationship between scores (Figure 3F) revealed that a large fraction of mutant were unfit in both conditions, as well clusters of variants with phenotypes similar to silent or nonsense mutations (green and red respectively). A potential explanation for this would be a very sharp trade-off resulting in a front minimum that leads to near vertical and horizontal curves outside of the inflexion point (i.e. the black line in Figure 1C). Due to the lack of resolution inherent to DMS assays, the precise nature of this relationship is difficult to determine from this type of data alone. To verify this hypothesis and validate the DMS scores, we selected 88 missense mutants across the two-dimensional phenotypic landscape inferred from DMS (Figure S12A) and generated the corresponding strain using genome editing by inserting the individual mutations at the *FCY1* locus. We included most mutants that showed large variation from the main clusters or that appeared to have high scores in both 5-FC and cytosine, as such outliers could have important implications for the emergence of antifungal resistance by escaping the trade-off. Based on the similarity of their DMS scores with nonsense or synonymous mutations, we assigned 46 mutants to clusters with another 42 that were considered outliers or had not been covered by the DMS assay.

For each mutant, we measured growth rates in 5-FC and cytosine DMS conditions (see methods). Growth rate replicates were highly correlated (Figure S12B and C). The 5-FC DMS scores were well correlated with validation growth rates (Figure S12G, ρ=0.72, 7.38×10^-14^), but the correlation was weaker for cytosine scores (Figure S12H, rho=0.30, 0.013), potentially due to noise introduced by uracil leaking in the large-scale experiment. This is supported by a stronger correlation between growth in cytosine and 5-FC+cytosine DMS scores for the mutants that were part of pool 2 (Figure S12I, ρ=0.568, p=0.0002, n=38). We observed a striking pattern in the relationship between a mutant’s growth rate in 5-FC and cytosine: all variants, including the outliers from the DMS screen, fell along the same front (Figure 4A). The relationship was well described by a power law equation with a negative k, characteristic of a strong trade-off, for which we determined parameters using a linear regression on the log_2_ transformed growth rates of individual mutants. In effect, this strong trade-off means that a Fcy1 mutation increasing growth rate in 5-FC media by ∼25% will incur a cost equivalent to ∼67% of growth rate of the wild-type in cytosine media. The steepness of the relationship generates a front minimum where a large fraction of mutants are unfit in both 5-FC and cytosine media, as seen in the DMS scores. We found that 45/46 of the mutants to which we had assigned a cluster in the DMS experiments maintained the same position in the validations. The better signal to noise ratio of the growth curve assay also allowed us to precisely measure the correlation between phenotypes, which was very strong (ρ=-0.851, p=1.83×10^-25^, n=87 variants). Finally, we note that the trade-off escape that was observed for some outliers of the DMS experiments could not be validated with these small scale validations.

**Figure 4:**
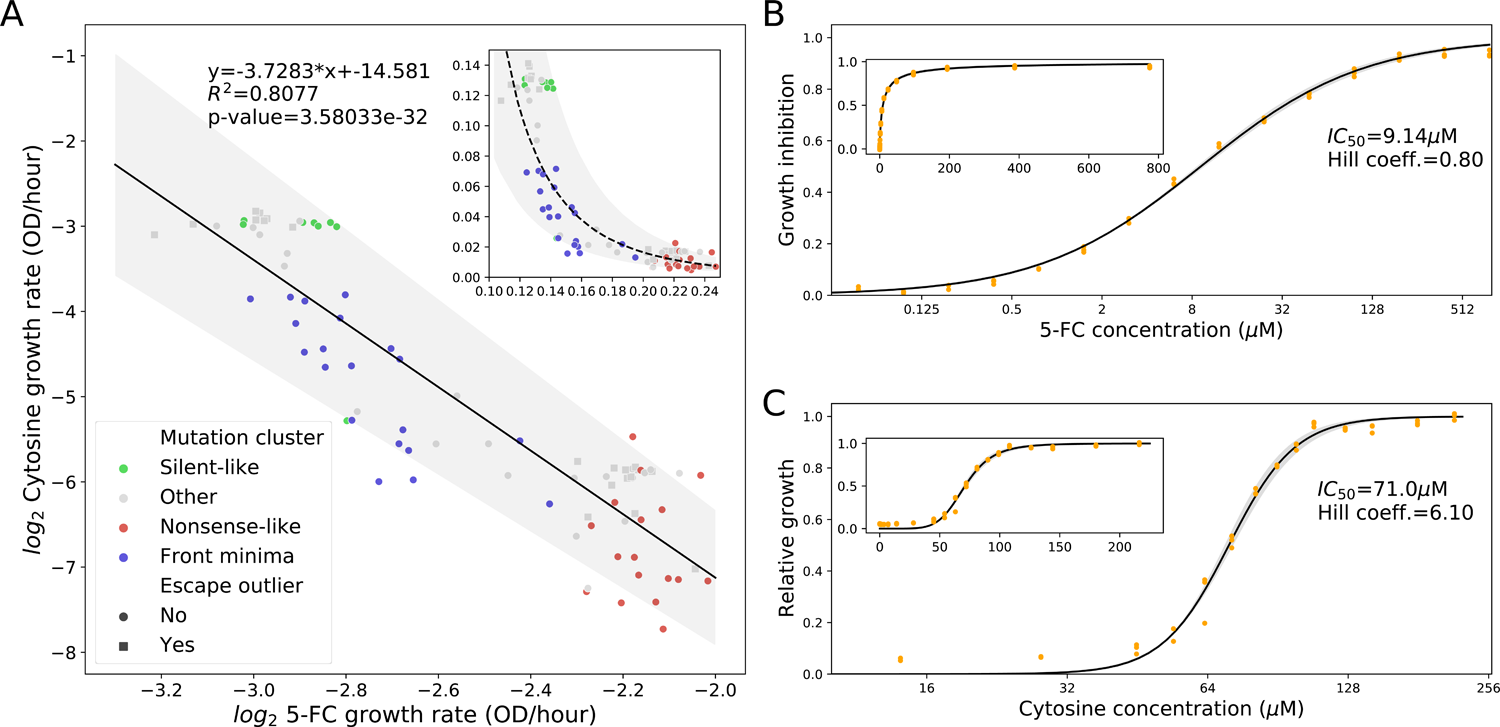
Fcy1 mutants are constrained to a small region of phenotypic space. **A)** Growth rates in 5-FC and cytosine of 87 validation mutants in the pooled competition conditions (SC + 12μM 5-FC or SC-Ura + 84 μM cytosine. The variants were assigned to different clusters based on DMS scores. Green variants clustered with silent mutations and exhibit a phenotype close to the wild-type. Red variants cluster with nonsense mutations, implying complete loss of function. Blue mutants had both low 5-FC DMS and cytosine DMS scores. Variants falling outside these clusters are shown in grey. The grey area represents the 95% confidence interval around the log-log linear regression slope fit. The inset shows the values on a linear scale. Variants that had 5-FC and DMS scores suggesting they might be mutants that escape the trade-off are shown as squares. **B)** Dose-response curve of wild-type *FCY1* to 5-FC and **C)** to cytosine. The Hill equation parameters were estimated using a non-linear least-square fit on relative growth rates, and the grey area represents the uncertainty around the dose-response curves based on the 95% confidence interval of the measured IC_50_ and hill coefficients. The inset shows the data on a linear scale.

Both the DMS results and our one-by-one growth curves point towards a sharp trade-off between resistance and cytosine use. We examined what the source of this strong trade-off might be. First, we examined wether the imbalance in the effect of mutations might be the result of large differences in the catalytic efficiency of Fcy1 when deaminating cytosine instead of 5-FC. However, based on reported values in BRENDA^37^, this is not the case: the distributions of catalytic efficiencies reported for the two substrates overlap, and the K_cat_/K_M_ of the median values shows only a small increase in efficiency for 5-FC deamination (111.9 mM^-1^s^-1^ vs 104.4 mM^-1^s^-1^, Figure S13). Furthermore, our dataset also contains some mutants whose presence in the front minima would be difficult to attribute to change in the catalytic efficiency of the enzyme. The stop codons mutants from positions 2-5, which did not confer 5-FC resistance, cluster with the front minimum mutants in the DMS data. These mutants are instead expected to affect phenotype through lower protein levels due to the alternative translation initiation site and not through changes in the *FCY1* coding sequence. These observations point toward a difference in the interplay between protein complex abundance or function and the response to 5-FC and cytosine levels as a way to explain the shape of the trade-off.

As the validations experiments showed, the phenotypes of Fcy1 in 5-FC and cytosine variants are strongly anticorrelated, but the effects are asymmetric. Additionaly, there does not appear to be any major difference between substrates in terms of metabolic flux for the wild-type enzyme. Accordingly, we hypothethized that the asymmetry we observe in the trade-off must be a cellular property linked to pathway flux and how it connects to growth rate, which should be reflected in the dose-response to each compound. While the dose-response assays shown in Figure S1 were sufficient to determine IC_50_ in preparation of our DMS experiments, they did not cover the full range of response to 5-FC and had only minimal coverage of the range of cytosine concentrations where most changes occurred. We therefore performed a second round of dose-response curves with a greater emphasis on precisely determining curve parameters (Figure 4B and 4C). The IC_50_ and Hill coefficient we had first determined were close to the values we found in our second round of experiments (5-FC: 12 μM vs 9.14 μM, cytosine: 84 μ M vs 71 μM). The most striking difference between 5-FC and cytosine response lies in the steepness of changes around the IC_50_. Growth inhibition by 5-FC varies along a gradient that spans a 2,048-fold difference in drug concentration: for cytosine, variation occurs within a range spanning only a 4-fold difference.

The dose-response curves explain the shape of the trade-off between 5-FC resistance and cytosine use we observe. Because of this difference in steepness around the IC_50_, the same level of loss of function leading to lower metabolic flux will result in different outcomes (Figure 5A). As the level of loss of function due to a mutation increases, the dose-curve response is expected to shift to reflect the loss of Fcy1 enzymatic activity (shown as the grey lines in Figure 5B and C), until there is no response to the drug or nutrient anymore (the red line). For example, a 2-fold decrease in effective 5-FC concentration due to lower Fcy1 activity starting from the IC_50_ would result in 36% (instead of 50%, Figure 5B) growth inhibition based on the dose-response curve parameters. The same change in effective cytosine concentration due to loss of activity instead leads to loss of the ability to grow (from 50% to 1.4% relative growth, Figure 5C). This leads to strong asymmetry that explains the front minimum we observed in the DMS experiment. This difference in dose-response should result in strong unidirectional effects, so that mutants having a full resistance phenotype would not recover the ability to use cytosine at higher concentrations. Conversely, some mutants in the front minima should be able to grow if provided with higher cytosine concentrations to offset their lower Fcy1 activity.

**Figure 5:**
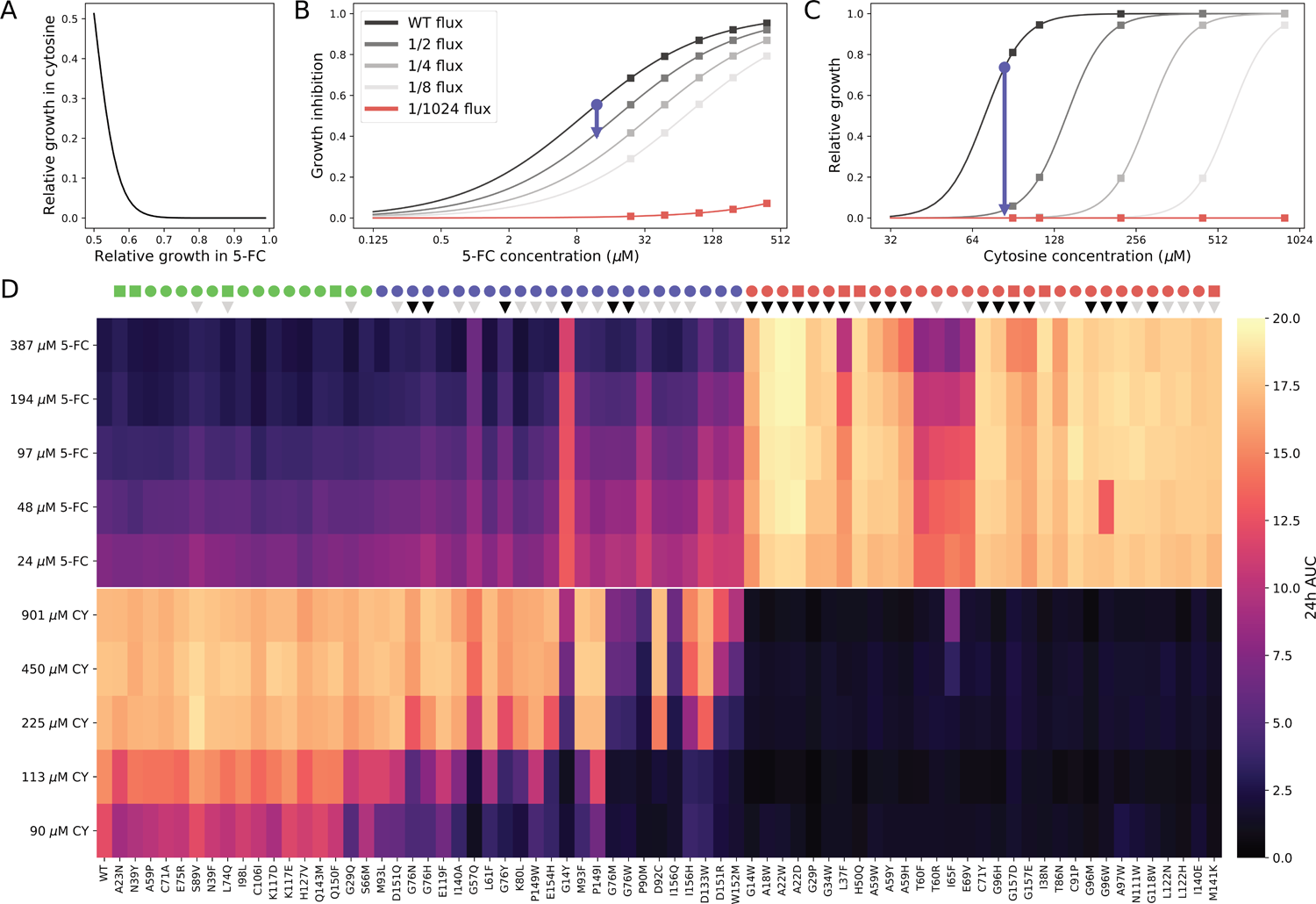
Dose-response parameters predict the properties of Fcy1 mutants. **A)** Relationship between growth in 5-FC and cytosine when concentrations drop beneath the IC_50_ based on the dose-response curves. Growth in cytosine media is abolished so quickly by lower flux that the same change has little effect on 5-FC resistance. **B)** Effect of reduced Fcy1 flux on growth inhibition by 5-FC and **C)** growth in cytosine media based on the dose-response curve parameters. The grey lines show the predicted dose-response curves if Fcy1 metabolic flux is reduced. The blue line represents the phenotype of a Fcy1 variant which reduces pathway flux by 50% in validation conditions (12 μL 5-FC or 84 μM cytosine). **D)** Response of validation mutants to higher concentrations of 5-FC and cytosine. The mutants were ordered based on their position along the trade-off front, starting from the top left in Figure 4A (no 5-FC resistance, full growth in cytosine). The colored dots above represent the cluster to which the mutants were assigned based on the validation data. Variants with mutations predicted to affect stability significantly are annotated with triangles: mutations annotated with grey triangles have a ΔΔ of 1-5 Kcal/mol and those with black triangles a ΔΔG >5 Kcal/mol.

To test our hypothesis, we performed growth curves and measured the area under the curve (AUC) at 24h for 72 validation mutants in five additional 5-FC and cytosine concentrations higher than previous conditions (Figure 5D, full growth curves shown in Figures S14 and S15 and AUC distributions in S16). Large increases in cytosine concentrations (450 μM and 901 μM) could restore growth to levels comparable to the wild-type for multiple mutants that showed no growth at concentrations closer to the IC_50_. Conversely, a few mutants showed some level of sensitivity to higher levels of 5-FC (i.e L37F, T60F, T60R), suggesting residual levels of Fcy1 activity. Many front minima mutants (i.e. G76H, K80L, D92C) had sharp on/off dynamics in cytosine, consistent with the shifted dose-response curve our model predicts. We observed differences in growth between 450 μM and 901 μM cytosine for some mutants, indicating that the activity of the cytosine permease *FCY2* was not limiting in our assay. Importantly, the relationship between responses to 5-FC and cytosine was correctly predicted. No fully resistant mutant recovered the ability to use cytosine, and resistance gains were always linked to greater losses in cytosine use. The phenotypes of Fcy1 mutants therefore support our trade-off model based on metabolic flux and shaped by different response patterns to 5-FC and cytosine.

Finally, we also found that the predicted effect of a mutation on structure stability was a strong predictor of whether it behaved as a complete loss of function (no response to either 5-FC or cytosine), as would be expected in a situation where protein complex abundance is critical. Mutations with a predicted ΔΔG from 1 to 5 Kcal/mol were significantly more likely to retain sensitivity or growth at higher compound concentrations compared to those with ΔΔG > 5 Kcal/mol (19/27 vs 6/18, odds ratio = 7.125, p= 0.002, two-sided Fisher’s exact test). Thus, we find that changes in protein stability, which translates into lower complex abundance, are an important factor in determining the level of loss of function of Fcy1 and predicting resistance.

### The resistance-function trade-off is conserved in a *FCY1* ortholog

Having characterized the relationship between 5-FC resistance and cytosine use, we wondered if our observations were specific to the *S. cerevisiae FCY1* (*scFCY1*) or was conserved in its orthologs. In clinical settings, 5-FC (flucytosine) is most often used against *Cryptococcus* species. We therefore tested whether mutants of the *C. neoformans* ortholog of *FCY1* (*cnFCY1*) were constrained by a similar trade-off. The two proteins share 41% residue identity and 58% residue similarity, implying a similar fold and catalytic site architecture. Since the structure of cnFcy1 has not been solved, we used AlphaFold^38, 39^ to predict it. We found it was almost identical to scFcyp (RMSD=0.631 Å over 141 residues, Figure 6A), with matching sequence alignment positions generally sharing the same position in the structure.

**Figure 6:**
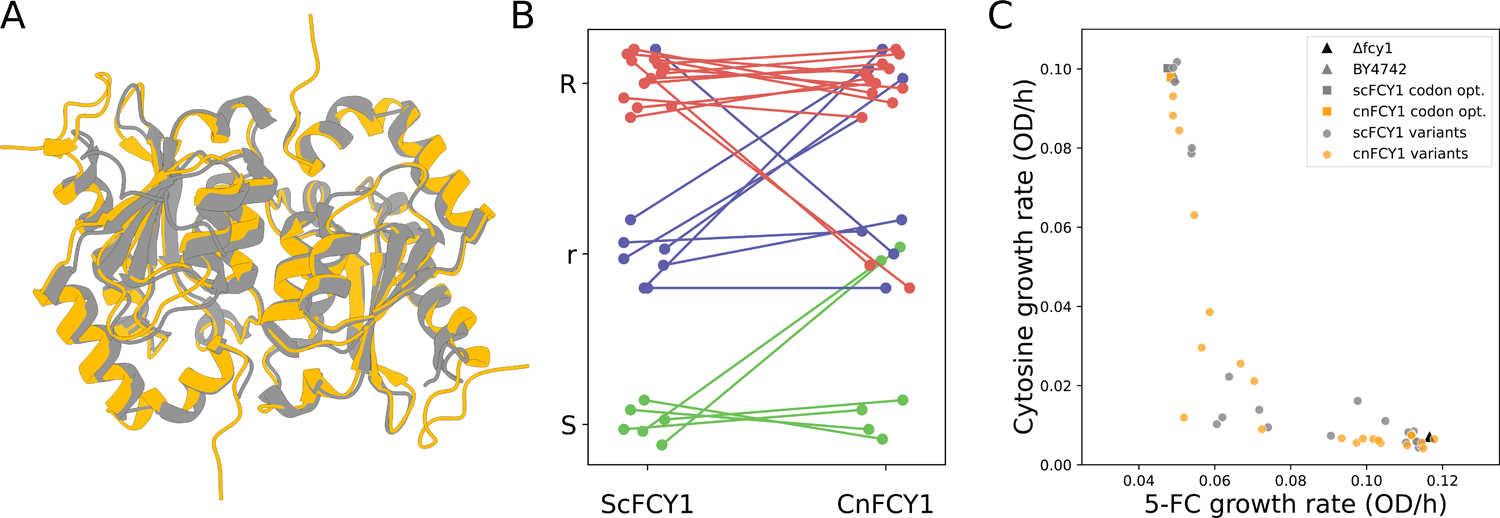
The resistance-function trade-off is conserved in the yeast *FCY1* ortholog of the pathogen *C. neoformans*. **A)** Structural alignment of cnFcy1 (in orange) as predicted by AlphaFold and scFcy1 (in grey, pdb: 1p6o). **B)** 5-FC resistance spot dilution assay phenotypes (see Figure S18 for images) for matched variant pairs, where S: sensitive, r: low growth and R: full resistance. Variant pairs are colored by the position of the *scFCY1* variant in the trade-off front. **C)** Resistance-function trade-offs of *scFCY1* and *cnFCY1* variants in the same conditions as the pooled DMS assays (SC + 12μM 5-FC or SC-Ura + 84 μM cytosine). The parental strain (BY4742, positive control) and the deletion strain (*fcy1*Δ, negative control) are shown as triangles. sc*FCY1* variants are shown in grey and those of *cnFcy1* in orange, with the wild-type codon optimized sequence shown as a square.

To test *cnFCY1* variants, we codon optimized the protein of the ortholog for expression in yeast and used it to replace the endogenous *FCY1* coding sequence. We then generated 27 strains with *cnFCY1* alleles matched to *scFCY1* validation mutants from across the phenotypic space (Figure S17A), prioritizing variants at conserved positions. We then performed spot dilution assays to compare the 5-FC resistance phenotype of each variant pair at high 5-FC concentration (194 μM). Yeast expressing *cnFCY1* showed the same level of sensitivity to 5-FC as either unoptimized or codon optimized *scFCY1* (Figure S17B). Among variants, we observed three phenotypes: wild-type like sensitivity, slow growth, and full resistance (Figure S17C). We found the majority of phenotypes were conserved within orthologous variant pairs (18/27, Figure S18). Interestingly, we did not observe any cases where a fully resistant mutant was fully sensitive or vice-versa in the ortholog. All changes in phenotypes were incremental, suggesting the effects of some variants differ just enough between the two proteins to move them slightly on the trade-off front, which is very steep.

Because of uracil leakage, growth using cytosine can not be assayed using spot assays like 5-FC resistance. We therefore selected a subset of 22 variant pairs for which we performed growth curves in screen conditions to measure both phenotypes. Strinkingly, we found that variants from both orthologs appear to evolve on the same trade-off front (Figure 6C). Growth rates in 5-FC and cytosine media are distributed similarly for both orthologs (D=0.23, p=0.63 for 5-FC and D=0.18, p=0.87 for cytosine, two sample Kolmogorov-Smirnov test). As in the spot dilution assays, phenotypes were strongly correlated within variant pairs (Figure S17D and S17E, ρ=0.600 and p=0.003 for 5-FC, ρ=0.461 and p= 0.031 for cytosine, n=22 pairs). Variants with diverging phenotypes did not deviate from the trade-off, as growth rate changes in the two conditions remained negatively correlated (Figure S17F). These cases of slight changes in phenotype for the same mutations in the ortholog are potential examples of epistasis whereby the effects of mutations are strongly contingent on other residues^40^. A staring phenotypic landscape shared by variants in the two orthologs suggests that the resistance-function we characterized here might be a generalizable property of cytosine deaminases from the *FCY1* orthogroup.

## Discussion

Systematically characterizing the impact of mutations on drug resistance and normal cellular function is a key aspect of improving our ability to design new molecules, manage existing ones and potentially exploit trade-offs associated with resistance mutations. These goals include interpreting mutations that have never been reported before and understanding the mutational target sizes of different drug targets. Using genome editing and DMS, we systematically characterized the impact on 5-FC resistance and canonical enzyme function of all single amino acid mutations in *S. cerevisiae* Fcy1 and this at its endogenous locus. We catalogued 940 missense and 151 nonsense mutations that conferred 5-FC resistance, nearly 33% of all single codon mutations. Resistance mutations were found all across the enzyme and not only near the active site. No single amino acid mutation was found to escape the resistance-function trade-off. Our computational work and experimental assays suggest this is largely due to effects on protein stability. The fact that destabilizing mutations in Fcy1 can be sufficient to lead to resistance points to the importance of maintaining the endogenous transcription levels for these assays as overexpression could mask relevant effects.

An important factor influencing the fate of resistance mutations is the cost associated with them. Understanding the mechanisms that determine the relationship between resistance and growth in the absence of drugs is important because it could eventually open the door to using conditions that maximize the cost of mutations. Here we show an example of an extremely steep resistance-function relationship and provide a mechanistic explanation. Our results demonstrate that the shape of a trade-off is inherently linked to how steep the function linking growth to flux through the pathway is compared to the function linking flux and growth inhibition by the toxic analog. During treatment, the target serum concentration of 5-FC is between 25 and 100μg/ml (194 to 775μM)^41^. In our dose-response curve, this corresponds to the upper plateau. Because of its low steepness, a mutation would have to reduce Fcy1 flux by over 400-fold to make the effective 5-FC concentration inhibit growth by less than 10%. At this level of loss of function, the remaining cytosine deaminase activity would be irrelevant to cell metabolism. The yeast membrane’s propensity for uracil leakage^25^ would also be expected to reduce the impact of Fcy1 loss of function, as an initial mutant would probably be surrounded by wild-type cells that could provide it with enough uracil to compensate for partial loss of the salvage pathway, alleviating the strength of the trade-off. The cost of resistance mutations could therefore be frequency dependent. How the shape of the trade-off we uncover affects evolution *in vivo* therefore needs to be investigated.

The characterization of trade-offs may help design better treatment strategies against pathogens. In the case of 5-FC, combining it with another compound that targets the *de novo* pyrimidine synthesis pathway could provide a way to reduce or abolish its ability to buffer loss of functions in the salvage pathways. For example, olorofim is a newly developed antifungal currently undergoing phase IIb clinical trials that inhibits fungal dihydroorotate dehydrogenase, which is part of the *de novo* pathway^42, 43^. While it has no activity against *Candida* and *Cryptococcus* species, olorofim has strong activity against molds such as *Aspergillus* species, many of which are also sensitive to 5-FC *in vivo*^44, 45^. New compounds such as these could provide opportunities for combination regimens that take advantage of the potency of 5-FC while slowing the emergence of resistance during treatment. One key result of our experiment is that we did not observe any mutants that escaped the strong resistance-function trade-off. This suggests that gaining resistance inevitably leads to loss in the ability to use cytosine.

The concept of trade-off is ubiquitous in biology^46^. We used a relatively simple model (one protein, two traits) to explore general questions linked with the evolutionary importance of trade-offs when conflicts arise between two traits. Our data provide an example of how whole-organism traits like dose-response dynamics can lead to strong trade-offs at the molecular level and create scenarios where they are inevitable. Arguably, one limitation is that we investigated single-step mutations and multiple-step mutations may be able to break this trade-off. However, in a situation where mutation rate is relatively weak and population sizes are limited, single-step mutations may be more representative of what could happen during treatment. Mutations outside of the *FCY1* locus could also provide alternative evolutionary paths allowing both resistance and nutrient use. Recent work has revealed new pathways and mechanisms in *C. neoformans* associated with 5-FC resistance outside the pyrimidine salvage pathway^47, 48^. Investigating the role of multiple genes at the same time in this context will provide insights on how increasing complexity and epistasis can influence the properties of trade-offs.

Antifungal resistance is a growing threat to both human health and food supply, but there is still limited data on the molecular mechanisms underlying it. By systematically exploring the effects of all possible single amino acid mutants, we hope to facilitate the interpretation of new genotypes in the clinical or industrial setting. Numerous efforts have been made in this sense in the field of antibiotic resistance in bacteria, where antibiotic resistance genes such as the β-lactamase TEM-1 have been thoroughly dissected and evolutionary pathways to resistance have been characterized^49^. In human genetics, DMS studies have helped shed light on the phenotypic effects of thousands of variants of unknown significance in disease or cancer genes^50^. In effect, the 1,091 mutants we identified will more than quadruple the number of amino acid substitutions recorded in the MARDy database, which catalogs all known antifungal resistance mutations^3^.

Although we focused on 5-FC resistance in a model species, the high correlation between the phenotypes of *scFCY1* and *cnFCY1* suggests our results could be used to predict resistance in pathogens. This could prove invaluable for interpreting population genetics of clinical isolates or mutants isolated in drug resistance studies. For example, genetic variation within pathogenic lineages sometimes leads to 5-FC resistance even in the absence of treatment, as observed for *Candida dubliniensis* isolates from the C3 clade, which bear a S29L substitution in its Fcy1 ortholog, Fca1. In our DMS experiment, the equivalent mutation in yeast Fcy1 (G35L) confers resistance to 5-FC^51^. Based on a review of known resistance associated mutation in fungal pathogens, our results correctly predict the resistance phenotype of 8/9 variants across 5 species (Supplementary table 1). Despite how widespread *FCY1* orthologs are, not much is known about their contribution to cell survival in natural or host environments, except that it can provide increased metabolic efficiency in some conditions^52^. The conservation of the enzyme across fungi suggests it plays a key role, which contrasts with the fact that it is dispensable in laboratory conditions. Further studies that examine other orthologs will help determine if the trade-off we observed for *scFCY1* and *cnFCY1* is a general property of the *FCY1* orthogroup. By generating a resistance mutation catalogue and elucidating the mechanisms between resistance and function, the data we obtained for the *S. cerevisiae* protein will help interpret genotypes from across fungi. Using this approach on other antifungal targets could help proactively identify resistance mutations before they are observed in the wild.

## Methods

### Strains, plasmids and culture media

Strains constructed as part of this study are listed in supplementary Table 2, plasmids used or constructed in supplementary Table 3, and oligonucleotides in supplementary Table 4. Media recipes can be found in supplementary Table 5. All yeast growth experiments were performed at 30°C unless stated otherwise and all bacteria were grown at 37°C. Liquid cultures were incubated with 250 RPM agitation. All Sanger sequencing was performed at the CHUL sequencing platform (Université Laval, Québec). All high throughput sequencing was performed using the MiSeq Reagent Kit v3 on an Illumina MiSeq for 600 cycles (IBIS sequencing platform, Université Laval).

All strains were constructed in the BY4741 and BY4742 genetic backgrounds^53^ unless stated otherwise. We used a standard lithium acetate transformation protocol^54^ to construct gene deletions strains. The *FCY1* deletion strains BY4741 *fcy1*Δ*::NatNT2* (strain s_001) and BY4742 *fcy1*Δ*::HphMX4* (strain s_002) were constructed by amplifying the *NatNT2* or *HphMX4* cassettes from pFA-NatNT2^55^ or pAG32^56^ respectively using oligonucleotides del_for and del_rev and transforming the product in competent cells of the parental strain. Gene deletions were verified by PCR using oligonucleotides FCY1_A, Nat_B or Hph_B.

### Cytometry competition assay

To measure competitive fitness of wild-type vs *fcy1*Δ cells, we constructed a fluorescent *fcy1*Δ We reporter strain where the endogenous copy of *PDC1* is tagged with *mCherry* (s_003). plified the mCherry tag from a modified version of plasmid pBS35^57^ where the am selection marker is *NatMX4* using oligonucleotides PDC1_tag_for and PDC1_tag_rev, and used a standard lithium acetate protocol to transform the cassette in BY4742 *fcy1*Δ*::HphMX4* and plated the cells on YPD+Nat. Positive transformants were validated by PCR (using oligonucleotides PDC1_C and mCherry_B) and by fluorescence cytometry using a Guava® easyCyte 8HT (excitation: 532 nm, detection 609 nm). To perform the competition assay, four independent pre-cultures of BY4742 and BY4742 *fcy1*Δ*PDC1-mCherry* were grown overnight in SC media. The following morning, the cells were diluted to 0.1 OD in SC media and allowed to reach exponential phase. Cells from each culture were centrifuged and resuspended in sterile water to a concentration of 1 OD. The culture pairs were then mixed in 1:9, 1:1, 9:1 (*FCY1:fcy1*Δ*PDC1-mCherry)* and 20μL of the mix was used to inoculate 180μL 1.11X SC-ura media with either 225, 113, 84, or 56μM cytosine so that the media reached 1X with the addition of the cells (final concentration: 0.05 OD) in a Greiner flat bottom transparent 96-well plate. The abundance of each genotype was then measured by counting the number fluorescent vs non-fluorescent yeast cells in each well, and the initial cell density was measured by counting the number of cells/μL. After ∼16h incubation at 30°C, the final OD for each well was measured, and the cells were diluted to 0.05 OD in sterile water. The abundance of each genotype and the cell density was measured again for all wells. The selection coefficient was calculated using the following approach for each condition and replicate:

First, we measure the number of doublings for wild-type and *fcy1*Δ*PDC1-mcherry* by comparing the absolute abundance of each genotype at the start and the end of the competition:

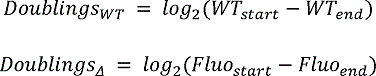

With the endpoint counts adjusted to the final optical density at 16h. We then use the number of doublings of the wild-type and *fcy1*Δ*PDC1-mcherry* strains to calculate the relative fitness based on the number of doubling of the wild-type:

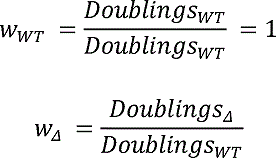

From there, since the selection coefficient is defined as

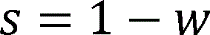

We have

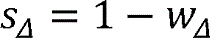

Which gives us the selection coefficient for the *fcy1*Δ*PDC1-mcherry* strain. As there was little to no overnight growth in the 56 μ cytosine samples, we excluded them from the selection coefficient calculation.

### Creations and validation of the *FCY1* DMS library

To be able to differentiate it from the wild-type sequence in eventual diploid assays, we decided to start from a codon optimized *FCY1* sequence to build the DMS library. The yeast codon optimized sequence was synthesized and cloned into a high copy vector (plasmid pTWIST-FCY1_opt, Twist Biosciences, San Francisco, USA). We then used the mega-primer method^22^ to generate codon by codon mutant libraries by using oligonucleotides with three degenerate nucleotides (NNN). In the first PCR step, the degenerate forward oligonucleotides are paired with a reverse primer to generate a short amplicon where one codon has been mutated:

| Reagent | Volume (μL) |
| --- | --- |
| Kapa HF buffer 5X | 3 |
| dNTPs 10 mM | 0.45 |
| Reverse Primer 10 μM | 0.45 |
| Codon-specific NNN oligo 10 μM | 0.45 |
| Kapa polymerase | 0.1 |
| Plasmid DNA 30 ng/μL | 0.23 |
| PCR grade water | 10.33 |
| Total | 15 |

With the following PCR cycle:

|  |  |  |
| --- | --- | --- |
| 95 °C | 5 mins |  |
| 98 °C | 20 s | 35x |
| 60 °C | 15 s |  |
| 72 °C | 30 s |  |
| 72 °C | 50 s |  |

For codons 2 to 57, the reverse oligonucleotide was mega_1, mega_2 for codons 58-133 and mega_3 for codons 114-158. The resulting mega-primer product was verified on agarose gel and used as a primer for the directed mutagenesis PCR reaction with the following parameters:

| Reagent | Volume (μL) |
| --- | --- |
| Kapa HF buffer 5X | 3 |
| dNTPs 10 mM | 0.45 |
| Mega-primer | 0.75 |
| Kapa polymerase | 0.1 |
| Plasmid DNA 30 ng/μL | 0.23 |
| PCR grade water | 10.48 |
| Total | 15 |

With the following PCR cycle:

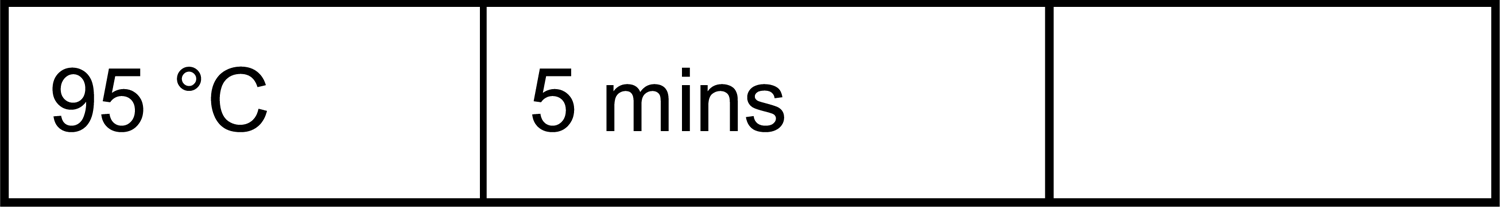
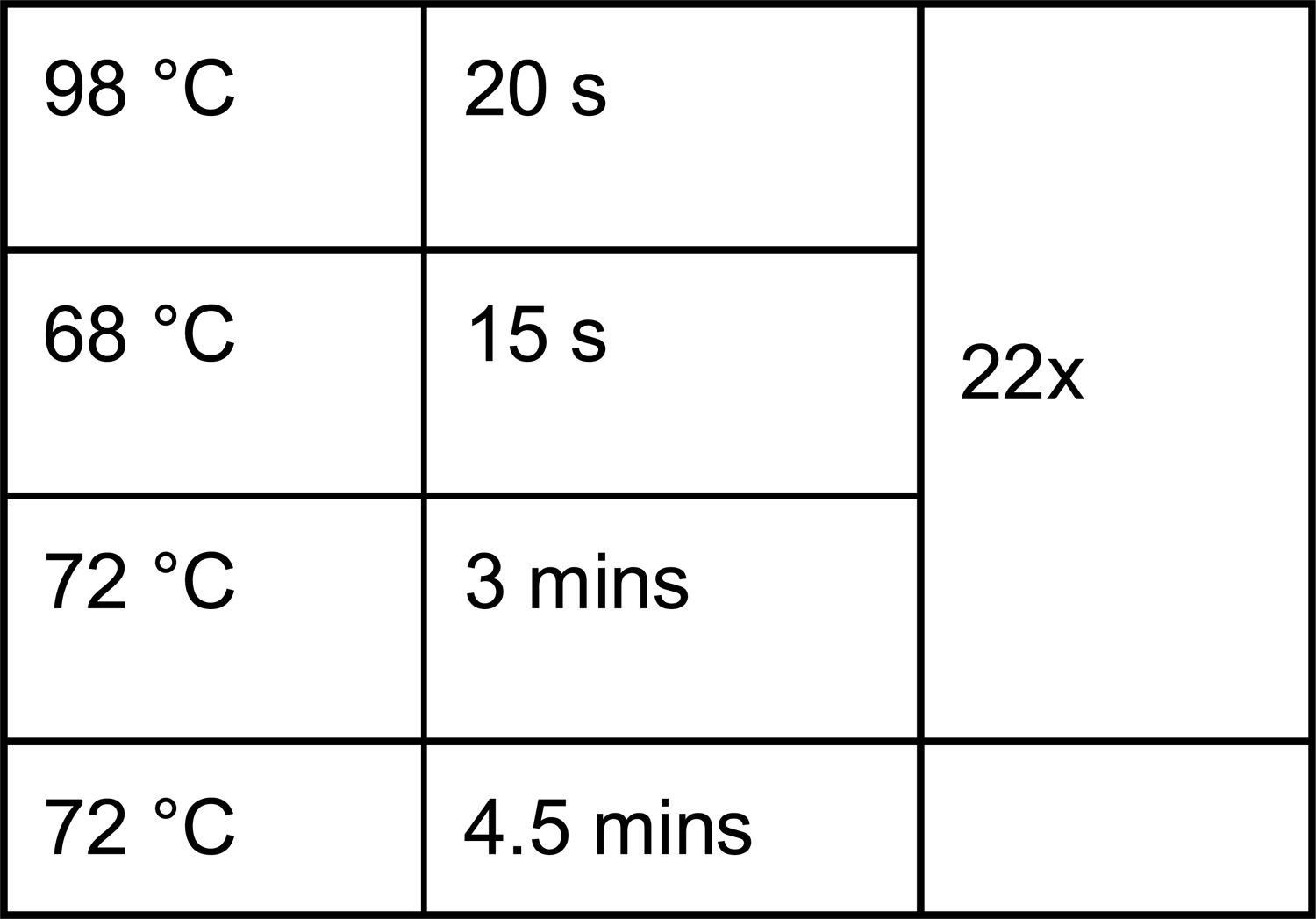

|  |  |
| --- | --- |
| 95 °C | 5 mins |

|  |  |  |
| --- | --- | --- |
| 98 °C | 20 s | 22x |
| 68 °C | 15 s |  |
| 72 °C | 3 mins |  |
| 72 °C | 4.5 mins |  |

After amplification, 0.3 μL 20,000 U/ml Dpnl (New England Biolabs) was added to the reaction mix which was then incubated at 37°C for 1 hour to digest the template plasmid. We l of the digested mutagenesis reaction into E. coli MC1061 *([araD139]_B/r_ Δ(araA-leu)7697 ΔlacX74 galK16 galE15(GalS) λ-e14-mcrA0 relA1 rpsL150(strR) spoT1 mcrB1 hsdR2)*^58^ using a standard chemical transformation method. This was sufficient to obtain hundreds to thousands of clones per codon, more than enough to cover all 64 possibilities.

The mutagenesis was performed in three batches corresponding to the three different reverse primers in the first PCR steps. For each codon, transformants were recovered by soaking the transformation plate with 5 mL 2YT media and scraping with a glass rake to recover cells in a 15 mL falcon tube. The pools were homogenized by vortexing and then archived by mixing 800 μl bacteria pool and 200 μL 80% glycerol for storage at −80°C. Finally, plasmid DNA was extracted from each pool with a standard miniprep kit (FroggaBio, Concord, Canada) using 1.5 ml of bacterial suspension as input and eluting in 50 μL kit elution buffer (10 mM Tris-HCl pH 8.5). After all extractions were completed, plasmid DNA concentrations were normalized to 1 ng/μL in two Greiner V-shape 96 well plate by first diluting to 10 ng/μL in 20 μL PCR grade water and then adding another 180 μL of PCR grade water. This plasmid DNA dilution was then used as the template for quality control of the DMS library.

To validate the mutant plasmid pools, we used Row-Column (RC) indexing^59^ to generate barcoded position by position libraries for high-throughput sequencing in three PCR steps. We split the *FCY1* coding sequence into three overlapping regions for Illumina paired-end sequencing. First, we amplified the appropriate *FCY1* region with oligonucleotides containing flanking universal priming sites (PS) at each end using with specific oligonucleotides (F1: pool_1_for, R1: pool_1_rev, F2: pool_2_for, R2: pool_2_rev, F3: pool_3_for, R3: pool_3_rev):

| Reagent | Volume (µL) |
| --- | --- |
| Kapa HF buffer 5X | 3 |
| dNTPs 10 mM | 0.45 |
| PS-Forward oligo 10 µM | 0.45 |
| PS-Reverse oligo 10 μM | 0.45 |
| Kapa polymerase | 0.1 |
| DMS plasmid library DNA 1 ng/μL | 4.5 |
| PCR grade water | 6.05 |
| Total | 15 |

With the following PCR cycle:

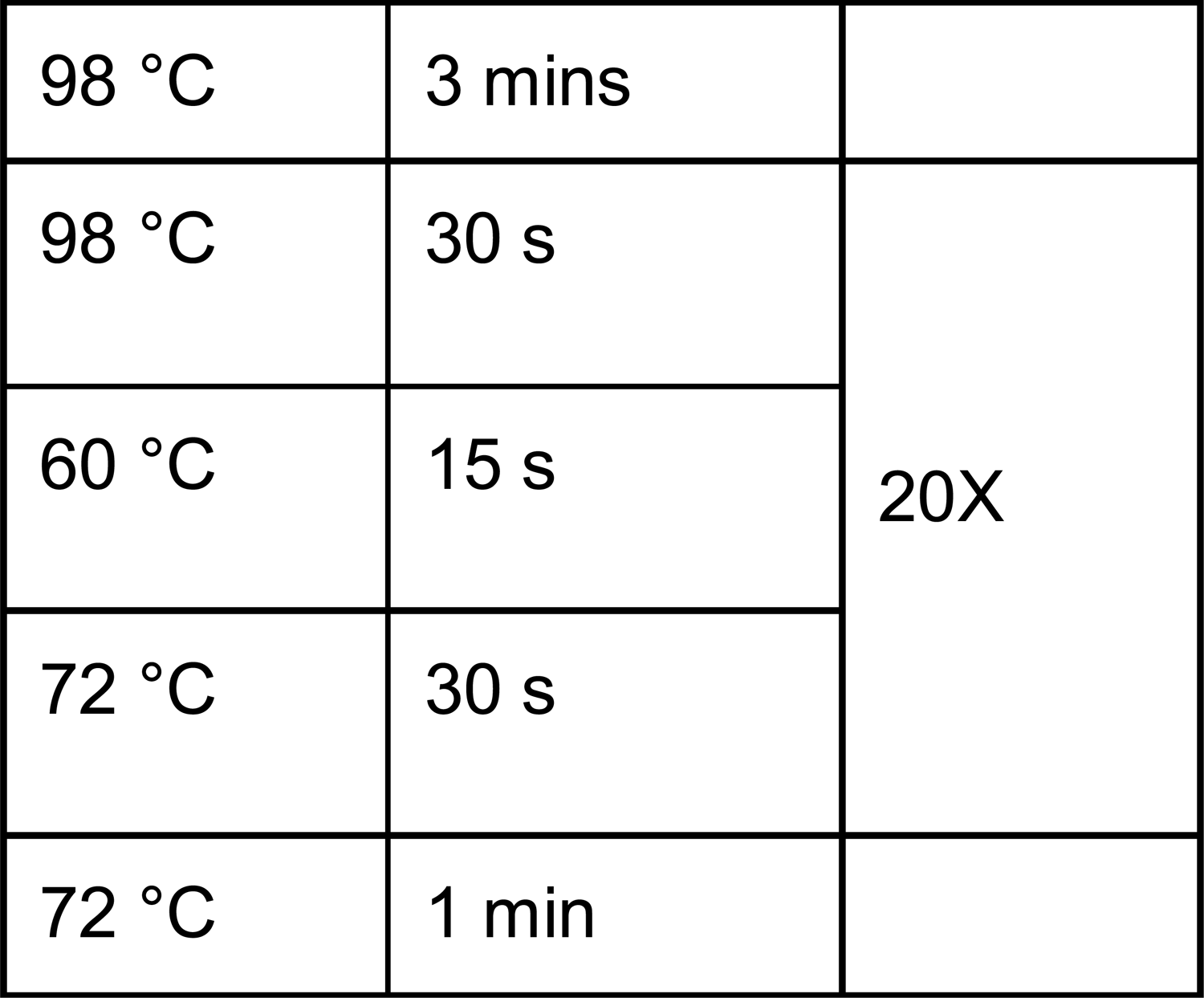

After gel verification, the amplicons were diluted 1/2,500 by two consecutive 2μL PCR: 98μL PCR grade water dilutions. This was then used as a template for a second PCR round where plate position specific barcodes were added at both ends of the amplicon (forward: oligonucleotides row_1 to row_8, reverse: oligonucleotides col_1 to col_12).

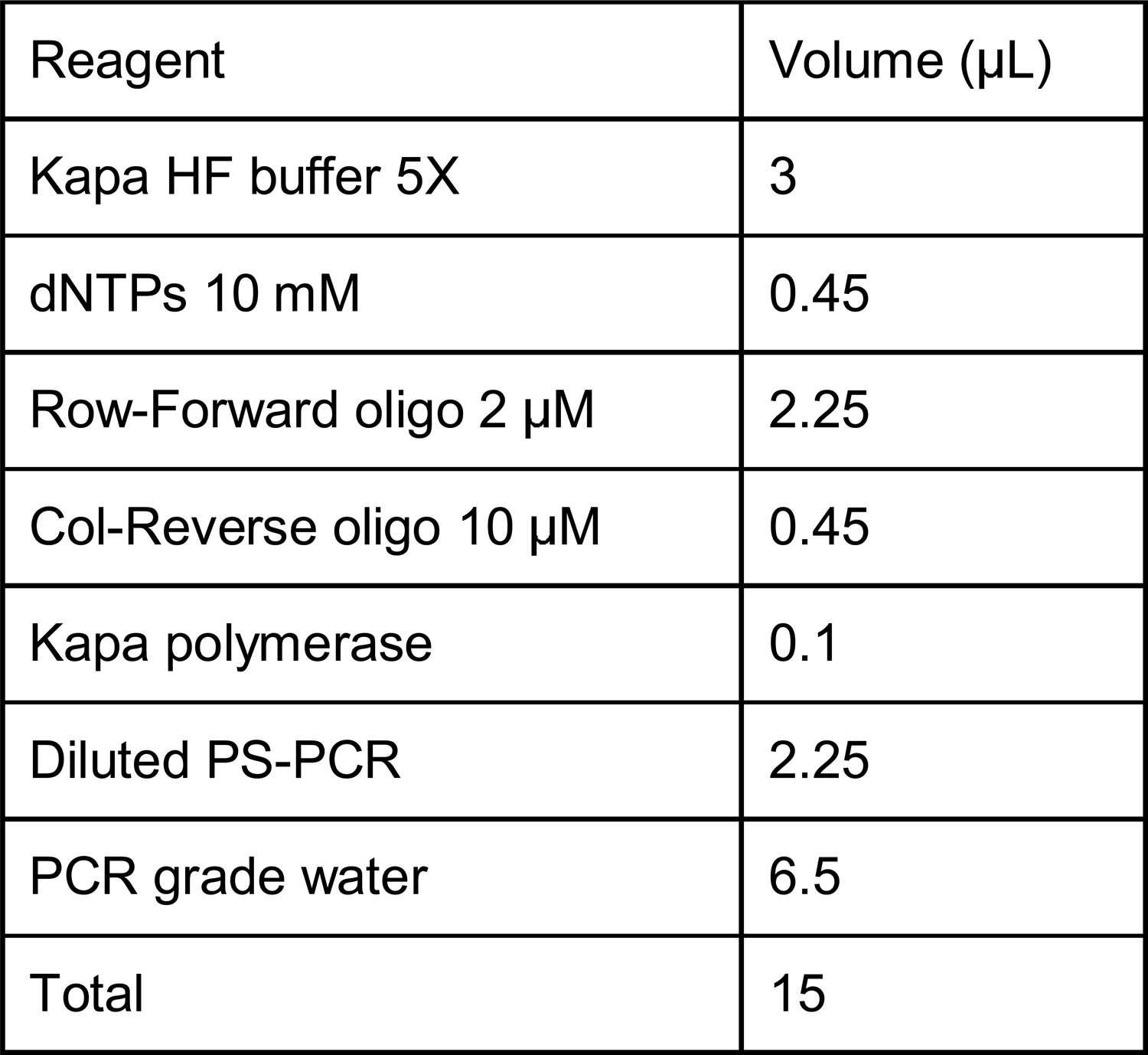

| Reagent | Volume (μL) |
| --- | --- |
| Kapa HF buffer 5X | 3 |
| dNTPs 10 mM | 0.45 |
| Row-Forward oligo 2 μM | 2.25 |
| Col-Reverse oligo 10 μM | 0.45 |
| Kapa polymerase | 0.1 |
| Diluted PS-PCR | 2.25 |
| PCR grade water | 6.5 |
| Total | 15 |

With the following PCR cycle:

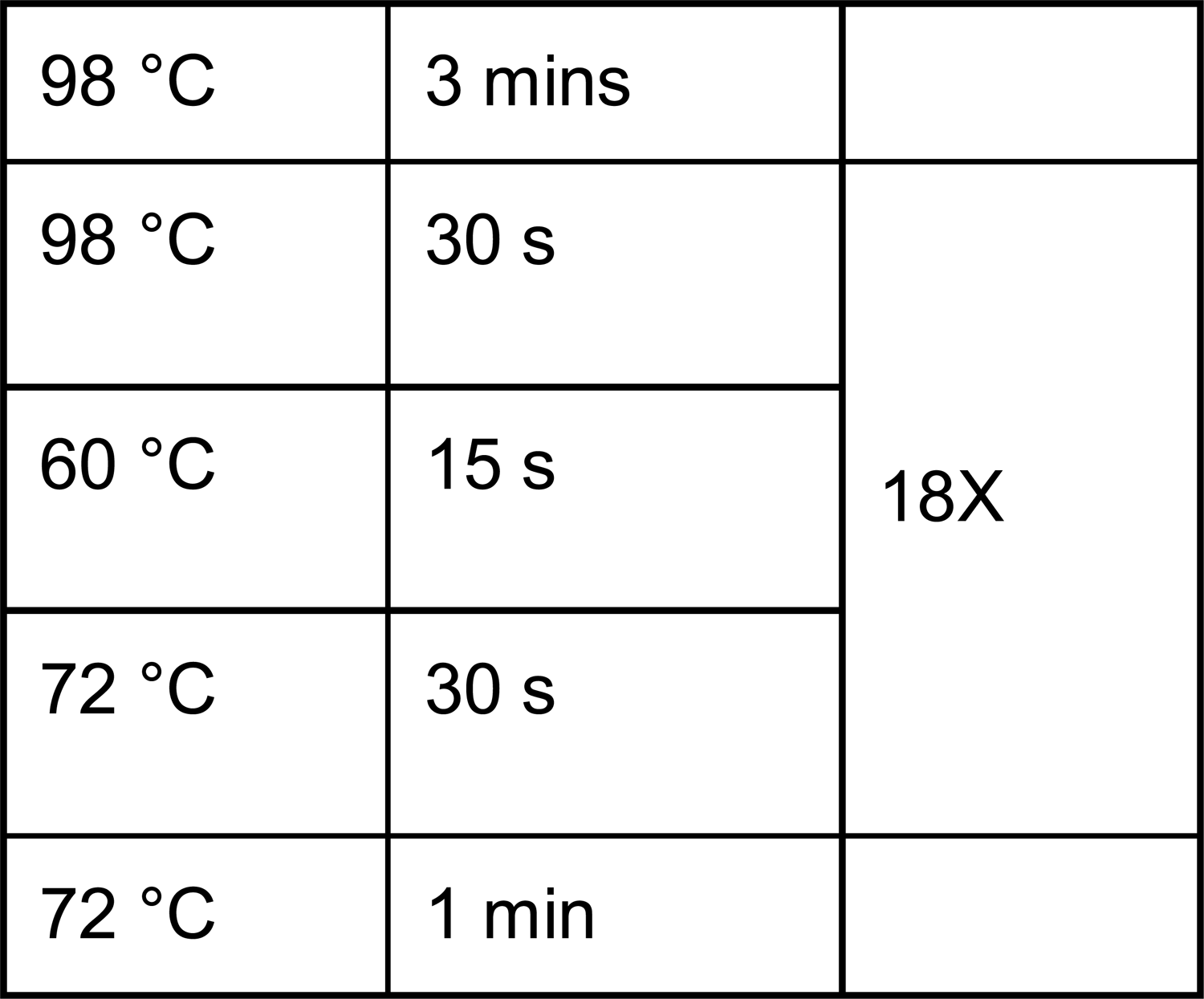

All barcoded amplicons of the same pool were then mixed in roughly equivalent quantities after gel band intensity quantification. The resulting amplicon mixes were purified using magnetic beads and used as templates for a final PCR round performed in quadruplicate to add plate barcodes at each end as well as the Illumina p5 and p7 sequences using a different combination of oligonucleotides for each pool (pool 1: plate_for_1/plate_rev_3, pool 2: plate_for_2/plate_rev_3, pool 3: plate_for_3/plate_rev_3):

| Reagent | Volume (μL) |
| --- | --- |
| Kapa HF buffer 5X | 3 |
| dNTPs 10 mM | 0.45 |
| plate_for oligo 10 μM | 0.45 |
| plate_rev oligo 10 μM | 0.45 |
| Kapa polymerase | 0.1 |
| Purified RC-PCR pool 0.01 ng/μL | 0.45 |
| PCR grade water | 10.1 |
| Total | 15 |

With the following PCR cycle:

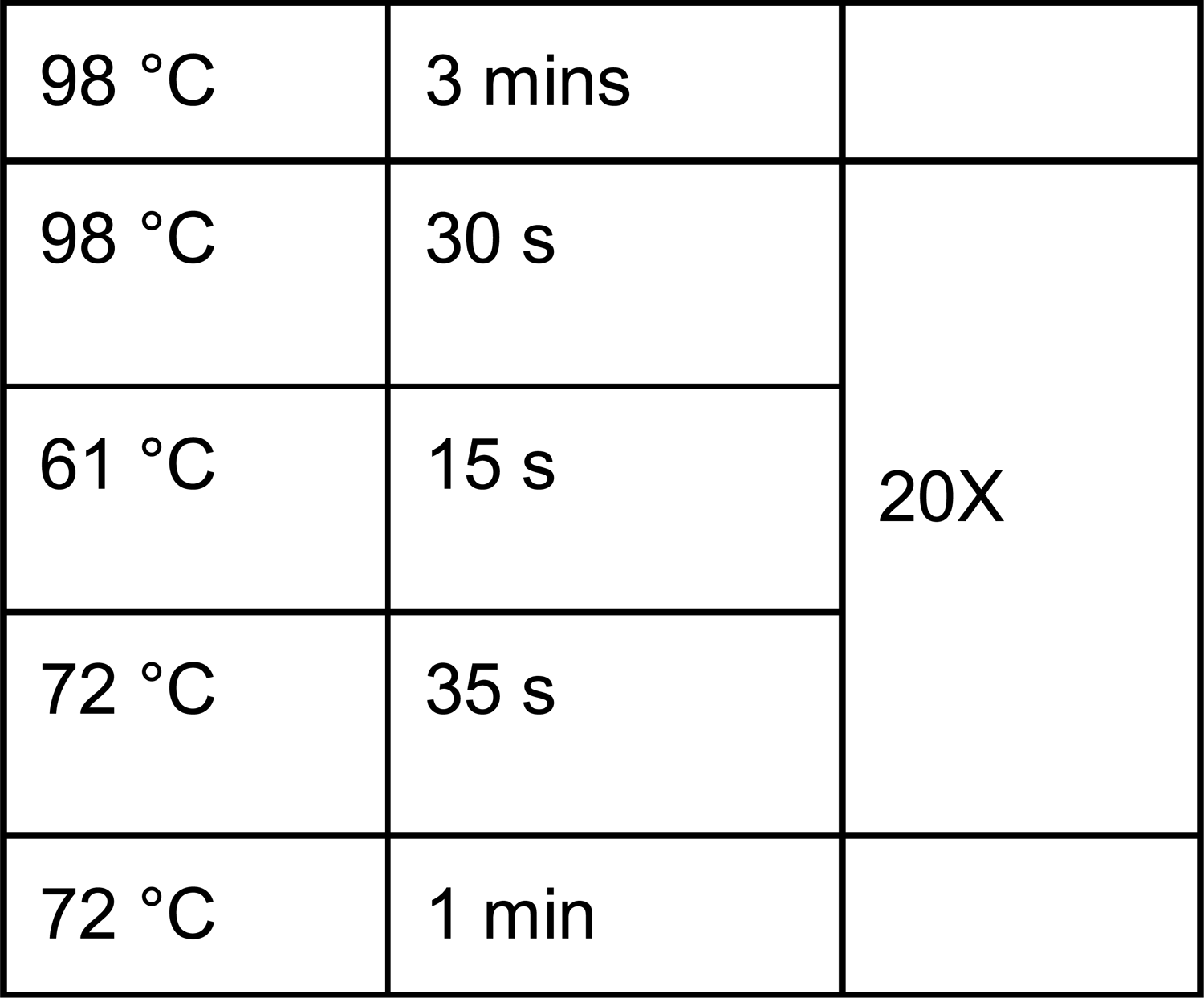

The replicate reactions were pooled and then purified on magnetic beads and gel verified to obtain the final libraries.

For a few codons (19, 80 and 91), the mutagenesis primers failed systematically and we therefore used equivalent degenerate oligonucleotides (alt_dms_19, alt_dms_80 and alt_dms_91) on the reverse strand with a matching forward oligonucleotide (mega_4) to generate the mutant plasmid pools. We also performed a second round of mutagenesis for some positions that showed poor coverage in the QC analysis (codons 83, 84, 107 and 108). These 7 pools were validated by Sanger sequencing.

### Next-Generation Sequencing data analysis

After sequencing, samples were demultiplexed using custom python scripts (https://github.com/Landrylab/Despres_et_al_2021) according to plate and RC barcodes after quality control with FastQC^60^ and trimming with Trimmomatic^61^. Data analysis scripts made use of the following packages: matplotlib^62^, numpy^63^, pandas^64^, seaborn^65^ and scipy^66^. After demultiplexing, the forward and reverse reads were merged using the PANDA-Seq software^67^ and collapsed into 100% identity clusters with vsearch^68^. Reads were then aligned to the appropriate *FCY1* fragment sequences using the Needle software from EMBOSS^69^. All raw sequencing files were deposited on the NCBI SRA, accession number PRJNA782569.

### CRISPR-Cas9 Knock-in of *FCY1* variants

We used a modified version of the protocol presented in Ryan et al. 2016^70^ to generate all the competent cells required for the DMS yeast transformations in a single batch. Colonies from a BY4742 *fcy1*Δ*::HphMX4* plate were used to inoculate a 20 mL YPD + Hyg overnight pre-culture. The following day, the cells were diluted to 0.2 OD in 300 mL YPD and grown to ∼1 OD. The culture was split in 50 ml falcon tubes and the cells were pelleted by centrifugation at 448 *g* for 2 minutes. The supernatant was removed and the pellet was washed with 0.5 mL LATE solution (100 mM Lithium acetate, 1mM Tris-Cl, 0.1 mM EDTA, filter sterilized) before being transferred to a 1.5 ml microcentrifuge tube. The cells were then pelleted again by centrifugation at 8,600 *g* for 1 minute, and the supernatant was removed. Finally, the cells were resuspended in 300 μl LATE solution and 300 μl 50% glycerol, mixed by a quick vortexing, and stored at −80°C until use.

To integrate the DMS pools for each codon in yeast, we co-transformed a pCAS^70^ variant bearing a gRNA targeting the *Hph* resistance gene (pCAS-Hph, p_005) as well as donor DNA amplified from the codon-specific mutant plasmid pool recovered from bacteria. The modified pCAS vector was generated using the same approach as in Dionne et al. 2021^24^, using oligonucleotides pCAS_hph_for and pCAS_hph_rev. The donor DNA for each codon was amplified using the following reaction:

| Reagent | Volume (µL) |
| --- | --- |
| Kapa HF buffer 5X | 3 |
| dNTPs 10 mM | 0.45 |
| Oligo knock_in_for 10 μM | 0.45 |
| Oligo knock_in_rev 10 μM | 0.45 |
| Kapa polymerase | 0.1 |
| DMS plasmid library 2 ng/μL | 3.5 |
| PCR grade water | 7.05 |
| Total | 15 |

With the following PCR cycle:

|  |  |  |
| --- | --- | --- |
| 95 °C | 5 mins |  |
| 98 °C | 20 s | 29X |
| 60 °C | 15 s |  |
| 72 °C | 45 s |  |
| 72 °C | 1.5 mins |  |

We performed one transformation/codon in a 96 well PCR plate using a similar approach to what is described in Dionne et al 2021^24^. Each well contained 20μL competent cells, 2 μL carrier DNA (salmon testes ssDNA 10 mg/mL, boiled for 5 min and then cooled on ice) and 3.75 μL 200 ng/μL pCAS-Hph, prepared as a master mix and dispensed before adding 10 μL donor DNA. A 200 μL volume of freshly prepared PLATE solution (40% PEG 3350, 100 mM Lithium acetate, 1mM Tris-Cl, 0.1 mM EDTA, filter sterilized) was then added to each well. The plates were sealed with sterile aluminum foil, vortex mixed, and incubated at 30°C for 30 minutes before a 15 minutes heat shock at 42°C. The plates were then centrifuged at 448 *g* for 6 minutes to pellet the cells, the supernatant was removed, 100 μL YPD was added, and the plates were sealed with aluminum foil again for a 3h30 outgrowth at 30°C. The cells were then plated on YPD+G418 with glass beads and incubated at 30°C for ∼64 hours.

To recover cells, the plates were soaked with 5 ml YPD, scraped with a glass rake and the cell suspension was transferred to a 15 mL falcon tube. The cell suspensions were homogenized by vortexing, and the OD was measured for each pool. Based on the cell concentration, a volume equivalent to 5 OD units of each pool was mixed to generate the three DMS mutant pools: pool 1 contained codons 2 to 67, pool 2 codons 49 to 110, and pool 3 codons 93 to 158. The OD of each pool was measured to measure their final cell concentration, and pool aliquots were archived by mixing 380 μL of cell pool with 180 μL of 80% glycerol. Pool aliquots were stored at −80°C until use. A 500 μL was spun down and used as input for genomic DNA extraction with the phenol/chloroform method^71^. Next-generation sequencing libraries were generated and sequenced on illumina for a final round of quality control to ensure proper coverage in each pool.

The same approach was used to build the strain BY4742 *FCY1_codon_opt* (s_004), but using the wild-type plasmid and oligonucleotides KI_short_for and KI_short_rev to generate the donor DNA, and recovering single colonies instead of scraping the entire transformation plate. Transformants were used to inoculate 1 mL YPD cultures, grown overnight, and then streaked on YPD. One colony from each streak was resuspended in 200 μL water and spotted on YPD as well as YPD+G418 and YPD+Hyg to check for pCAS loss and successful loss of the *HphMX4* cassette. The strain was then validated by PCR and Sanger sequencing.

### DMS library screen and library preparation

An archived aliquot of each pool was thawed ∼30 minutes on ice. From the corresponding tubes, a volume corresponding to 2 OD units was used to inoculate 20 ml SC media in duplicate. The cells were grown for 24h and then split in the 5-FC and cytosine conditions: SC + 12 μM 5-FC or SC-ura + 84 μM Cytosine. Each 12h, 2 OD units of cells were centrifuged for 5 minutes at 224 *g*, the supernatant was removed, and the pellet was resuspended in 1 ml sterile water. The cell suspension was then used to inoculate 19 mL of 1.05X media, so that the addition of the cells brought the media to 1X. The competition was stopped once pools reached ∼20 generations based on OD measurements: this took two passages for the 5-FC condition, and 4 passages for the cytosine condition. Genomic DNA was extracted from 2 OD pellets (approximately 2×10^7^ cells) of the final passage with the phenol/chloroform method. The same approach was used for pool2 and the 5-FC+cytosine condition, but with SC-ura + 24 μM 5-FC + 113 μM cytosine.

To prepare NGS libraries for deep sequencing, we used a similar method as was used for the plasmid pool QC, but barcoding individual timepoints instead of codons, and with each reaction being performed in duplicate:

| Reagent | Volume (μL) |
| --- | --- |
| Kapa HF buffer 5X | 3 |
| dNTPs 10 mM | 0.45 |
| Pool forward oligo 10 μM | 0.45 |
| Pool forward oligo 10 μM | 0.45 |
| Kapa polymerase | 0.1 |
| DMS screen gDNA 30 ng/μL | 5 |
| PCR grade water | 5.55 |
| Total | 15 |

With the following PCR cycle:

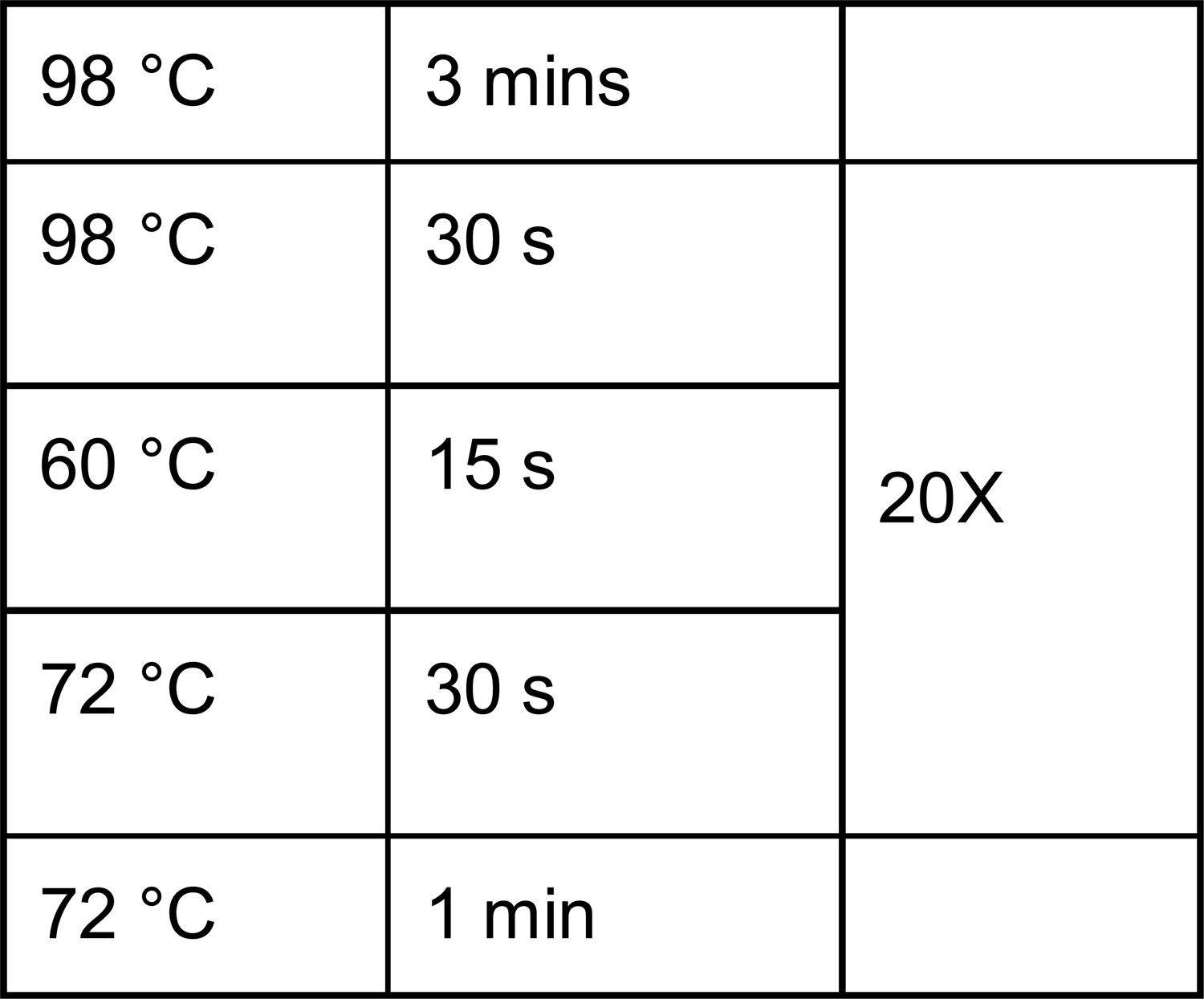

After gel migration, the amplicons were diluted 1/2,500 by two consecutive 2 μL PCR: 98μL PCR grade water dilutions. This was then used as a template for a second PCR round where plate position specific barcodes were added at both ends of the amplicon (primers row_1-3 with primers col_1-6), with each reaction performed in duplicate.

| Reagent | Volume (µL) |
| --- | --- |
| Kapa HF buffer 5X | 3 |
| dNTPs 10 mM | 0.45 |
| Row-Forward oligo 2 µM | 2.25 |
| Col-Reverse oligo 10 µM | 0.45 |
| Kapa polymerase | 0.1 |
| Diluted PS-PCR | 2.25 |
| PCR grade water | 6.5 |
| Total | 15 |

With the following PCR cycle:

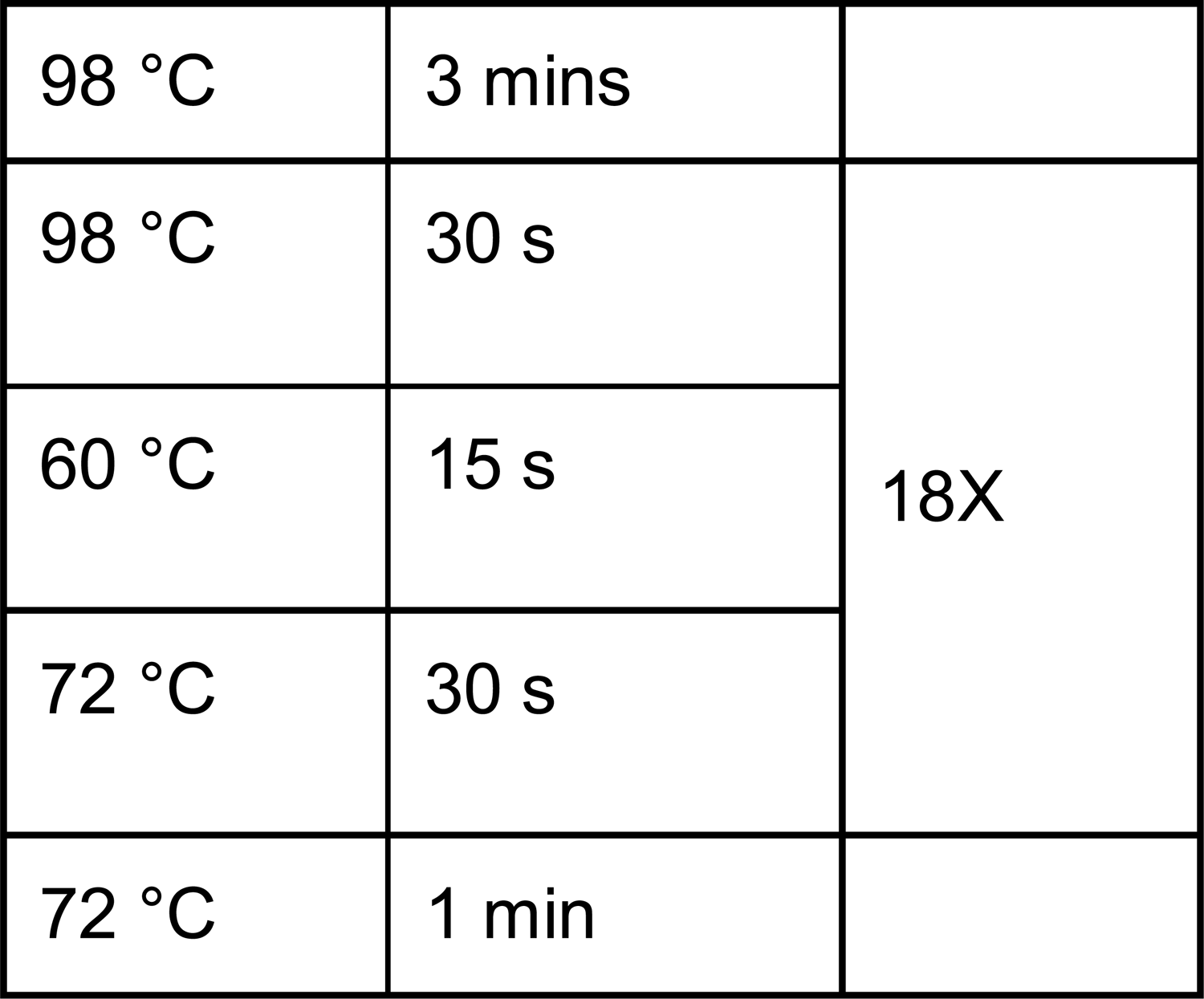

Replicate reactions were pooled together, purified using magnetic beads, and quantified using a NanoDrop spectrophotometer. Barcoded amplicons of the same pools were then mixed in equivalent quantities. The resulting pools diluted and used as templates for a final PCR round performed in quadruplicate to add plate barcodes at each end as well as the Illumina p5 and p7 sequences:

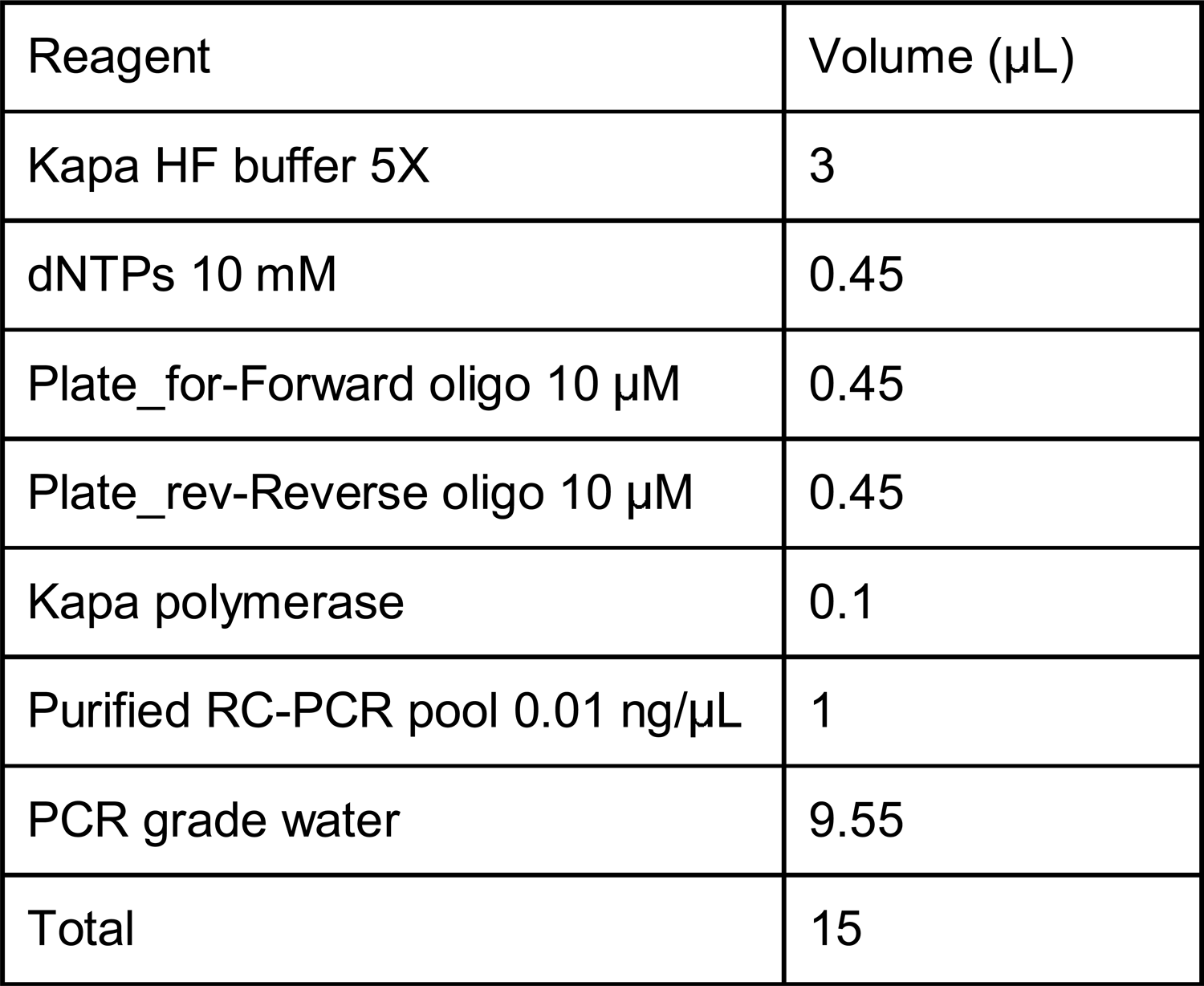

With the following PCR cycle:

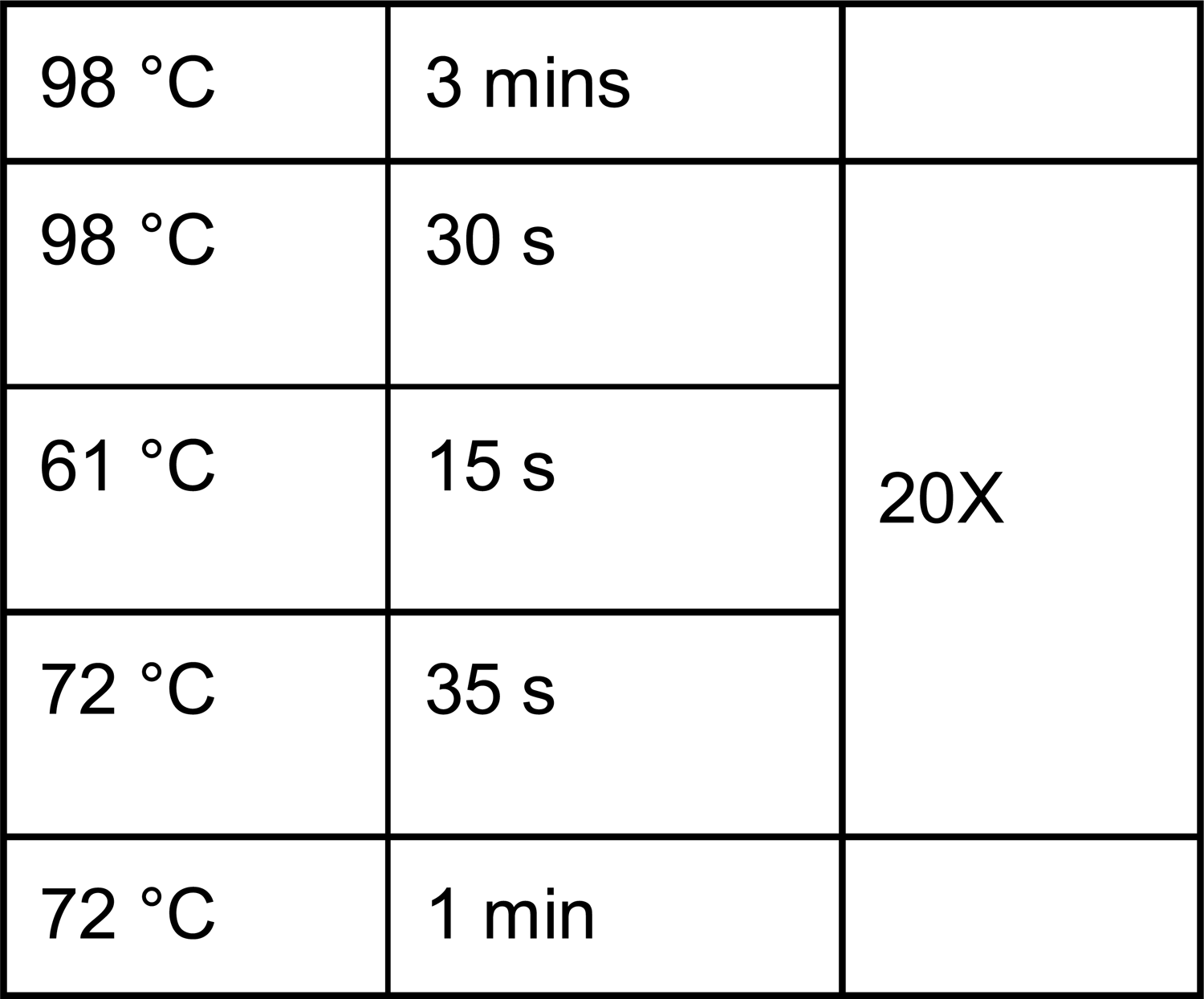

The replicate reactions were pooled, purified on magnetic beads and gel verified to obtain the final libraries.

### DMS score

Data analysis of the DMS screen was performed using custom Python scripts. Briefly, samples were demultiplexed into individual fastq files containing merged reads using the same strategy as was used for the plasmid pool QC analysis. Identical reads were then collapsed using vsearch^68^ and aligned on the appropriate *FCY1* region using Needle^69^ to detect mutations. At this stage, we filtered alignments to only keep query sequences that covered 80% or more of the reference and that had less than 25 differences. Reads with more than three mutations in the target DMS sequence or with mutations in multiple codons were discarded. Finally, we applied a stringent filter to avoid including double mutants: only reads where all differences in the *FCY1* coding sequence occurred within the same codon were considered when measuring variant abundance. The total number of read counts obtained per library is shown in Supplementary Figure 4A, and raw read counts for each codons in each library are provided as Supplementary table 6.

When computing log2 fold-change between timepoints, we followed the standard practice of adding one to all variant counts to make fold-change calculations possible for variants that dropped out during the competition. Variant read counts were then divided by total read count (including the WT sequence) to obtain relative abundances. Relative abundances were then compared between the end of the SC media passage and the end of the selection conditions (5-FC or cytosine) to obtain log_2_ fold-changes for all mutant codons. We discarded all codon level variants that were covered by less than 10 reads in one of the T0 replicates (prior to selection), which was a rare occurrence (Supplementary Figure 4B). Most codon level variants were well covered in the initial timepoints (median codon variant coverage > 60 reads for all pools, Supplementary Figure 4C), which lead to have a high amino acid level coverage (Supplementary Figure 4D). To convert to amino acid log2 fold-change, we considered each group of synonymous codons as replicates and calculated the median fold-change of each group. This allowed us to obtain three (one per pool) amino-acid level matrices per condition. Finally, we averaged log_2_ fold-change between replicates (as shown in Figure S4). As some *FCY1* positions in pools 1 and 3 overlapped with those in pool 2 (positions 49 to 67 and 93 to 110 respectively), ∼400 mutants/pool had their log_2_ fold-change measured in duplicate. As the fold-changes of same mutants were well correlated between pools and spanned the full range of possible values, we fit a linear regression that allowed us to adjust all scores to pool 2. For mutants in the overlapping regions, we used the average of the two log2-fold changes (one from pool 2, and the adjusted value from pool 1 or 3).

Once we had harmonized log_2_-fold changes along our three pools, we wanted the values for both experiments (5-FC and cytosine) to be on the same scale to facilitate comparison. To do this, we scaled values using the median log2-fold change of synonymous mutations and nonsense mutants using the following equations:

For 5-FC:

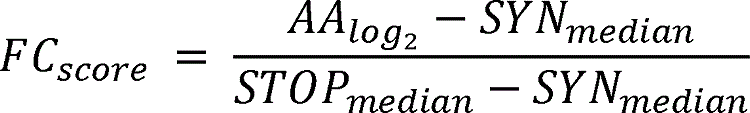

For cytosine

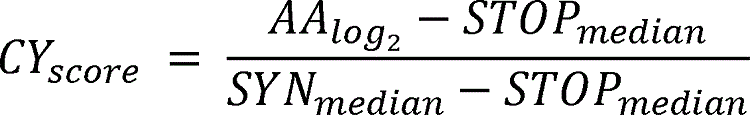

This scaling relates the log_2_ fold-change observed for a DMS mutant with the difference observed between the values of synonymous (silent) and nonsense (stop) mutants. For example, a score of 0.5 would represent a log_2_ fold-change value that is perfectly in the middle between the medians of synonymous and nonsense mutations. We chose to set the score according to the phenotype measured, so that in the 5-FC condition nonsense mutations would have a median FC_score_=1 and synonymous mutation would have a median median CY_score_=0. Conversely, in the cytosine condition, synonymous mutations would have a median CY_score_=1 and nonsense mutations would have a median median FC_score_=0.

### Amino acid and mutation properties analysis

We used the structure from Ireton et al. 2003^15^ (PDB: 1P6O) to compute residue relative solvent accessibility (RSA), distance to the active site and interface, and as baseline for the FoldX^32^ ΔΔG predictions which were performed using the same approach as in Marchant et al. 2019^72^. Briefly, the biological assembly was generated and energy minimization was performed using the FoldX Repair function 10 times to ensure convergence^73^. Then, the FoldX BuildModel and AnalyseComplex functions were used to estimate mutational effects on ΔG. As mentioned in the text, FoldX predictions were generally comparable between dimer bearing one or two mutated copies. RSA and temperature measurements of both chains were averaged to obtain the final value. The distance to the active site was considered to be the shortest distance between a specific amino acid and any active site residue. To determine which residues were part of the interface, we used a definition based on distance to amino acids in the other subunit, where residues whose two closest non-hydrogen atoms are separated by a distance smaller than the sum of their van der Waals radii plus 0.5 Å are considered to be at the interface (coded as 1). Residues whose alpha carbons are within 6 Å of the other chain were considered to be at the interface rim (near the interface, coded as 0.75)^72^.

We retrieved a set of 218 high confidence *FCY1* orthologs by combining sequences from Emsembl^74^, MetaPhOrs^75^, and the Yeast genome order browser^76^. The sequences were aligned using MUSCLE^77^ with default parameters. After visual inspection, three sequences were removed as they were discordant with major aligned blocks, yielding a final set of 215 sequences (Supplementary Data 1). The orthologs were aligned with Muscle again, using the stable.py script to keep the same sequence order in the output. Surprisingly, we found a higher than expected proportion of orthologs in bacterial and archaeal genomes. Previous work had hypothesized that cytosine deaminases from the *FCY1* family were exclusive to fungal species, with bacteria usually having orthologs of *E. coli codA*^15^. We instead found over 30 orthologs in bacteria and seven in archaea, suggesting a more complex evolutionary history for this orthogroup. This alignment was then used to calculate normalized evolutionary rates using Rate4site^34^ using default parameters. To determine variant occupancy, the positions of the alignment corresponding to the *S. cerevisiae* amino acids were parsed to identify the frequency of each amino acid. The fully annotated dataframe containing log2 fold-changes, 5-FC and cytosine score, as well as all annotations for mutants is provided as Supplementary table 7.

### Construction of the *FCY1* mutants validation set

The *FCY1* variant strains tested as part of the validation studies were constructed using the same strategy used to generate the DMS library, starting from BY4742 *fcy1::HphMX4*. We initially selected 80 variants for validation studies, but not all strain constructions attempts were successful, explaining why data is not presented for all of them. To generate mutated donor DNA, we used a fusion PCR strategy where oligonucleotides containing the mutations of interest are used to generate two overlapping amplicons both bearing a mutation starting from genomic DNA extracted from strain BY4742 *FCY1_codon_opt*. The first reaction, which amplified from the five-prime flanking homology arm before the start of the coding sequence up to the mutated amplicon overlap, used oligonucleotides fusion_for and mutant_rev_1 to 80. The second reaction, which amplified from the mutated overlap to the three-prime homology arm after the coding sequence, used oligonucleotides mutant_for_1 to 80 and fusion_rev. Both resulting amplicons are then diluted 1/2,500 and used as template for a second reaction using oligonucleotides fusion_for and fusion_rev to reconstitute the full length *FCY1* sequence.

We then performed one by one CRISPR knock-ins using the same reaction setup as for the DMS library, but using the fusion PCR products as donor DNA. Transformants (four/mutant) were used to inoculate 1.2 mL YPD cultures, grown overnight, and then streaked on YPD. One colony from each streak was resuspended in 200 μL water and spotted on YPD as well as YPD+G418 and YPD+Hyg to check for pCAS loss and successful loss of the *HphMX4* cassette. *FCY1* integration was verified by PCR using oligonucleotides FCY1_alt_A and fusion_rev on quick DNA extractions^78^. PCR positive strains were then validated by Sanger sequencing using fusion_for as the sequencing oligonucleotide. We succeeded in constructing 73/80 mutants. Variant growth rates were measured by performing growth curves in 384 well plates with a 80 μL sterile water border. Cells from overnight pre-cultures in SC media were spun down and resuspended in sterile water at a concentration of 0.4 OD. From these dilutions, 20 μl was used to inoculate either 60 μL 1.33X SC media + 12 μM 5-FC or 60 μl 1.33X SC-ura media + 84 μM cytosine, for a final media concentration of 1X and 0.1 OD cell density. Each strain was tested in duplicate for both conditions: measurements were taken every 15 minutes for 48 hours without agitation on a Tecan Spark (Tecan Life Sciences, Männedorf) with an active Tecool temperature control module. We excluded one mutant due to a technical problem with the plate cover, leading to the final count of 72 mutants. To further examine variants that appear to escape the trade-off, we constructed an additional 15 strains with high 5-FC and cytosine DMS scores, as well as some mutants that could not be measured in the DMS assays. The growth rates of these mutants in SC + 12μM 5-FC or SC-Ura + 84 μM cytosine was measured in triplicate (48 hour, 15 minutes intervals) using growth curves in 96 well plates in a Tecan Infinite M Nano (Tecan Life Sciences, Männedorf). A set of 15 mutants present in the first round of validations was used to scale the growth rates of this experiment to those of the initial validations using a linear regression (Figure S12 D-F). The data and annotations related to validation mutants is presented in supplementary table 8.

### Measurement of mutant-WT dimer interaction

Although the Fcy1*-*Fcy1 homodimer interaction had already been reported to be detectable using this DHFR-PCA^79^, we reconstructed a full set of DHFR strains (s_005 to s_008) and validated that we could observe it. The C-terminal DHFR-tagged strains were constructed by amplifying the DHFR tags from pAG25-DHFR[1,2] and pAG32-DHFR[3] using oligonucleotides from the DHFR collection (DHFR_tag_F and DHFR_tag_R)^35^. The tagged strains were validated by PCR and Sanger sequenced.To generate a N-terminal DHFR[1,2] tagged allele of Fcy1, we used fusion PCR to amplify the DHFR[1,2]-linker fragment from pGD009^80^ (oligonucleotides Nter_DHFR_for and Nter_DHFR_rev) and then fuse it to the codon optimized sequence (oligonucleotides linker_fcy1_for and knock_in_rev) which was amplified from strain *FCY1_codon_opt* (s_004) genomic DNA. The fused amplicon was then used as donor DNA for a CRISPR transformation of *fcy1::HphMX4* with pCAS-Hph. The positive clone was validated by PCR and Sanger sequencing, yielding strain s_009 (*DHFR[1-2]-FCY1_codon_opt*). Mutant alleles tagged in N-terminal with DHFR[1,2] were generated using the same strategy, but using genomic DNA from strain s_009 as template instead. We were able to successfully construct 75/80 N-terminal DHFR[1,2] tagged strains.

To measure changes in PPIs with wild-type Fcy1, tagged variants were crossed with BY4741 *FCY1-DHFR[3]* by mixing 5 μl of a SC-complete DHFR[1,2]-*FCY1* mutant pre-culture with 5 ul of a YPD+Hyg *FCY1-DHFR[3]* pre-culture in 2 ml YPD and incubating ∼8 hours at 30 C. After incubation, 5 ul/mating was spotted on SC-lys-met + Hyg (pH buffered) media and incubated overnight at 30°C to select for diploids. Parental strains were also spotted in parallel to control for background growth. Cells from each spot were resuspended in sterile water and spotted a second time on SC-lys-met+Hyg to perform a second round of selection. From these spots, we inoculated SC-lys-met pre-cultures that were grown overnight. The next morning, the cells were spun down and resuspended in sterile water at a concentration of 0.4 OD. From these dilutions, 8 μl was used to inoculate either 72 μl 1.1X MTX media or 72 μl 1.1X DMSO media (see supplementary Table 5), for a final media concentration of 1X and a 0.1 OD cell density. Each strain was tested in duplicate for both conditions: measurements were taken every 15 minutes for 48 hours without agitation in a Tecan Spark plate as described above.

### Dose-response curves and growth rates measurements in multiple 5-FC and cytosine concentration

The dose-response growth curves were generated by measuring the maximum growth rate of BY4742 in 20 different concentrations of 5-FC or cytosine in SC and SC-ura media respectively. The range of concentrations tested was specific to the compound to ensure proper coverage of the dynamic range. The growth rates were measured in triplicate, with each replicate starting from a different colony. The values used to fit the Hill equation are presented as supplementary Table 9. The shifted dose response curves shown in figure 5B and C were obtained by adding a term to take into account Fcy1 loss of function in the Hill equation, so that the 5-FC or cytosine concentration is lowered based on diminished metabolic flux:

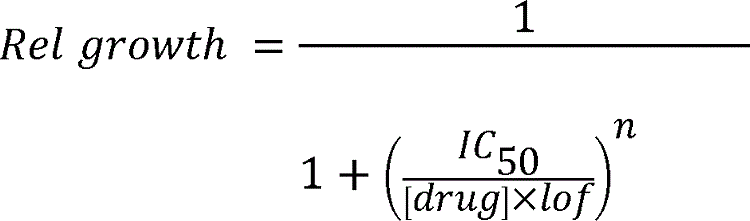

Where *IC_50_* and *n* are the hill equation parameters determined from the dose-response curve, and lof is a term representing the level of Fcy1 loss of function (e.g. *lof* = 0.5 is a loss of half the wild-type metabolic flux).

We measured the growth rates of Fcy1 mutants in 96 wells plates that contained the 73 Sanger validated mutants, of which 13 were present in duplicate. The plate also included 4 replicates of *FCY1_codon_opt* and 4 replicates of BY4742, as well as two blank wells with culture media and no cells. The growth curves were started by diluting overnight pre-cultures and inoculating 180 μL 1.11X SC or SC -ura media containing different amounts of 5-FC or cytosine. The optical densities were manually read for each plate using a Tecan Spark with an active Tecool temperature control module at regular intervals for 24h, at which point the cultures were fully resuspended and a final datapoint was acquired. The AUC measured for the wild-type strains (*FCY1_codon_opt* and BY4742) clustered together in all assay conditions (Figure S16A), and AUCs for replicate Fcy1 mutant cultures were well correlated across all conditions (ρ=0.8-0.99, n=13 mutants) except in SC media without 5-FC, which is expected to be non-selective for Fcy1 function (Figure S16B). For mutants present in duplicate, the values for each replicate were averaged, and the WT AUC was obtained by taking the median of the 8 WT measurements (the median of 4 replicates of *FCY1_codon_opt* and 4 replicates of BY4742). Data from variant T86M was excluded from Figure 5 because no cytosine growth rate from the validations was available. Raw curves for all mutants are shown in Figure S14 (5-FC) and S15 (cytosine).The AUC values used to generate Figure 4D and Figure S16 are available as supplementary Table 10.

### Generation of orthologous mutants in the *Cryptococcus neoformans* ortholog

The protein sequence of the *Cryptococcus neoformans* JEC21 ortholog of FCY1 (*CnFCY1*) was retrieved from NCBI and codon optimised for expression in yeast using the same tool as for the S. cerevisiae protein. To obtain the structure of cnFcy1, we used Alpha Fold 2^38^ via the ColabFold interactive notebook^39^ to predict its structure. We used the cnFcy1 protein sequence as input with default parameters and following the guidelines for dimer structure prediction. All output files for the prediction are provided as Supplementary Data 2. The structures were aligned using the align function of ChimeraX^81^, resulting in a pruned structural alignment with a length of 141 residues with RMSD = 0.631 Å (unpruned alignment length = 151 aa, RMSD = 3.129 Å). The coding sequence was synthesised and cloned into a high copy vector (plasmid pTWIST-FCY1_opt_JEC21, Twist Biosciences, San Francisco, USA) with homology arms for recombination at the *FCY1* genomic locus. We then used the same CRISPR-Cas9 mediated knock-in strategy to insert the wild-type allele at the *FCY1* locus and generate a *S. cerevisiae* strain bearing *CnFCY1*. We then selected 24 variants from validations that occurred at sites conserved between the two orthologs and designed oligonucleotides to generate the corresponding mutations in *CnFCY1* by fusion PCR. Because mutations at conserved sites tended to have more deleterious effects, we also included 5 mutants at non-conserved sites that had wt-like phenotypes to cover the whole range of cytosine media growth rates.

We generated the mutants using the same protocol as previously described for *S. cerevisiae* variants, but using genomic DNA from the CnFCY1 as a template. We were able to generate and validate 27/29 strain by Sanger sequencing. The spot dilution assays were performed by growing strains overnight in SC media and then adjusting them to 1 OD in sterile water. For each strain, we spotted 5 μL of each dilution on SC complete and SC complete + 194 μM 5-FC. Phenotypes were scored after 48h of growth. Matched growth curves were performed in duplicates on a randomly chosen subset of variant pairs. Cells from the spot assay pre-cultures were diluted to 0.1 OD in the same media used for the validations. OD measurements were taken every 15 minutes for 48h in 96 well plates using a a Tecan Infinite M Nano (Tecan Life Sciences, Männedorf). The maximum growth rate in both conditions was then calculated using the same method as for previous validations. Growth rate values of scFCY1 and cnFCY1 variants used to generate Figure 5C and Figure S17A and D-F are available as Supplementary Table 11.

## Acknowledgements

The authors thank members of the Landry lab for helpful discussions, in particular Daniel Evans-Yamamoto, Mathieu Hénault, Florian Mattenberger and Romain Durand. This work was supported by the Canadian Institutes of Health Research Foundation grant 387697 to C.R.L. and a Vanier graduate scholarship to P.C.D, as well as by the National Science and Engineering Research Council through the EvoFunPath CREATE grant (555337-2021) and by FRQNT through team grant (2022-PR-298169) and a PBEEE scholarship to A.F.C. CRL holds the Canada Research Chair in in Cellular Systems and Synthetic Biology. Figure 1 was generated using Biorender. Molecular graphics and analyses performed with UCSF ChimeraX, developed by the Resource for Biocomputing, Visualization, and Informatics at the University of California, San Francisco, with support from National Institutes of Health R01-GM129325 and the Office of Cyber Infrastructure and Computational Biology, National Institute of Allergy and Infectious Diseases.

