## Supplementary Figures 1-18 for "Asymmetrical dose-responses shape the evolutionary trade-off between antifungal resistance and nutrient use"

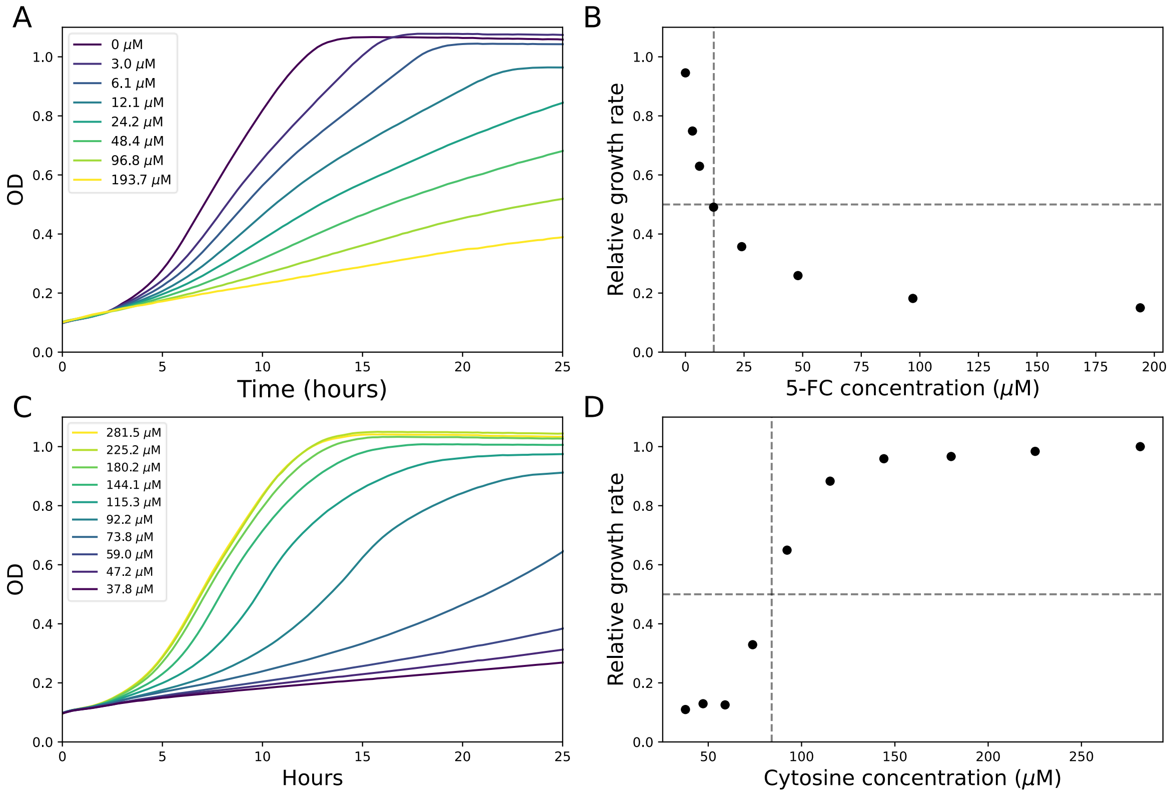


**Figure S1: Growth titration of cytosine and 5-FC to identify concentrations at which growth rate is comparable. A)** Growth of yeast strain BY4742 (*MATα his3Δ1 leu2Δ0 lys2Δ0 ura3Δ0*) at different concentrations of 5-FC in synthetic complete media. **B)** Growth of BY4742 as a function of 5-FC concentration relative to growth in the absence of 5-FC. A reduction in growth of 50% is observed at 12 μM 5-FC. **C)** Growth of BY4742 at different concentrations of cytosine in synthetic complete media depleted in uracil. **D)** Relative growth of BY4742 as a function of cytosine concentration compared to growth at the highest tested concentration. Based on this data, 84 μM μg/ml cytosine was interpolated as the concentration that would reduce growth by 50%.

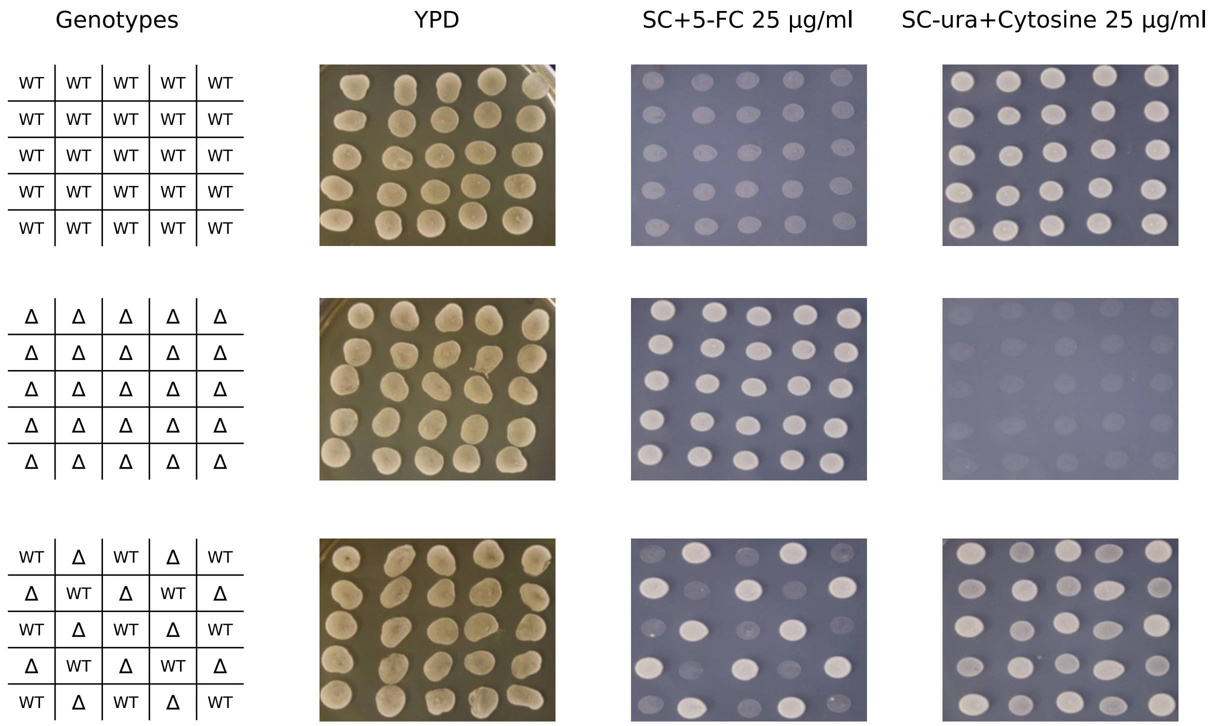


**Figure S2: Wild-type cells can rescue the phenotype of *fcy1Δ* cells when co-cultured on solid media.** Cells in the exponential phase were spotted (5 ul at 0.1 OD) on non-selective media (YPD), media with selection against Fcy1 function (SC + 5-FC) and media with selection for Fcy1 function (SC-ura + cytosine). Pictures were taken after 48h incubation at 30°C. See Supplementary Data 3 for source images.

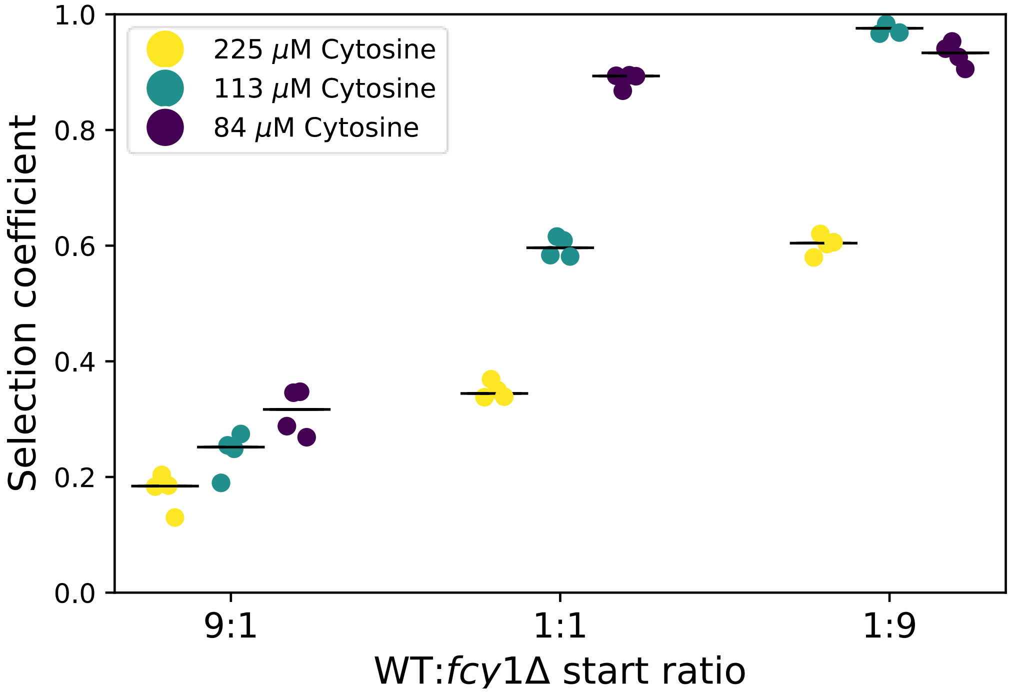


**Figure S3: Population composition and cytosine concentration change the selection coefficient of the *FCY1* deletion.** The x axis shows different initial population compositions at the start of the competition of WT cells and *fcy1Δ* cells. The selection coefficient represents the difference in relative fitness between the wild-type, which is considered to have a fitness of 1, and *fcy1Δ* relative fitness (population doublings of the mutant/population doublings of the wild-type). The black bars represent the median of the four replicates for each concentration by population combination. For the same ratio of wild-type to *fcy1Δ*, lowering the cytosine concentration increases the selection coefficient associated with the deletion.

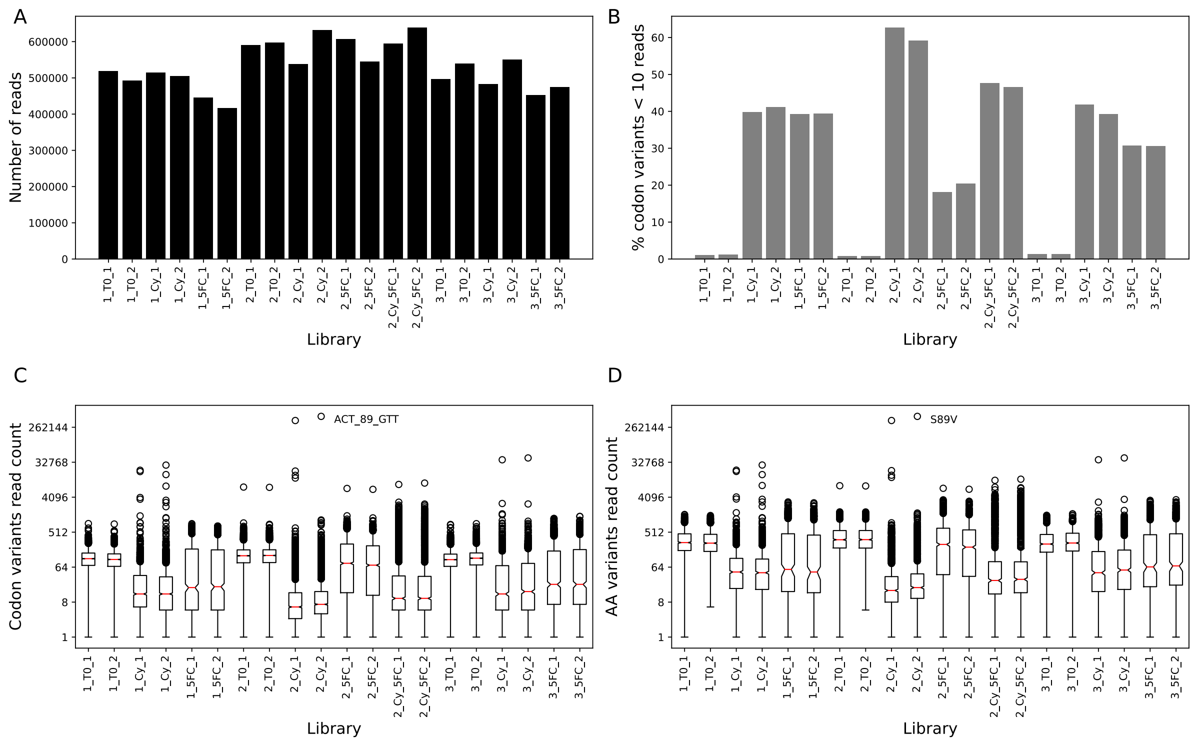


**Figure S4: Raw library and variant read counts for the DMS pooled assays. A)** Total number of reads per library after quality control filtering. **B)** Fraction of variants covered at a depth smaller than 10 reads in each library. Before selection (T0 timepoints), less than 1% of codon level variants have less than 10 read counts. The pre-selection libraries thus contain almost all possible FCY1 single codon variants. The diversity and average read counts drop significantly because of strong selection imposed during the competition. **C)** Codon level read counts in each library. **D)** Distribution of amino acid level raw read counts in each library. Boxplots represent the upper and lower quartiles of the data, with the median shown as a red bar and notches denoting the 95% confidence interval around the median. Whiskers extend to 1.5 times the interquartile range (Q3–Q1) at most. While the ACT_89_GTT (S89V) variant greatly increases in abundance after cytosine selection, results from the validation studies (see Figure 4A) did not find this mutation to provide an increase in growth rate in cytosine, suggesting a secondary mutation unrelated to *FCY1* that arose during strain construction or competition.

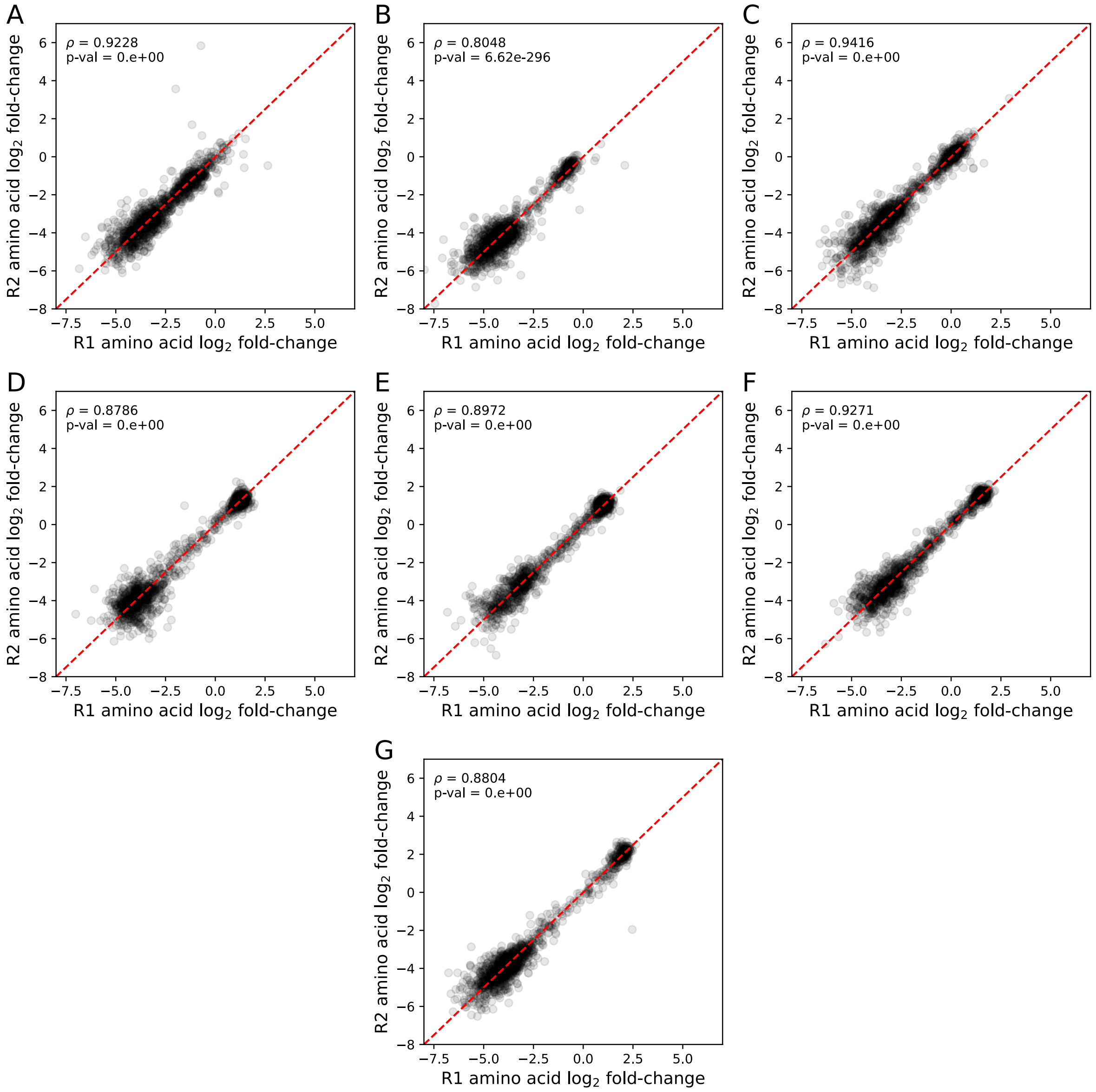


**Figure S5: Replicate amino acid $\log_2$ fold-changes in the DMS experiments.** Correlation between R1 and R2 was measured using Spearman's rank correlation. **A)** Pool 1, 5-FC (n=1372). **B)** Pool 2, 5-FC (n=1298). **C)** Pool 3, 5-FC (n=1374). **D)** Pool 1, cytosine (n=1372). **E)** Pool 2, cytosine (n=1298). **F)** Pool 3, cytosine (n=1374). **G)** Pool 2, 5-FC + Cytosine (n=1298).

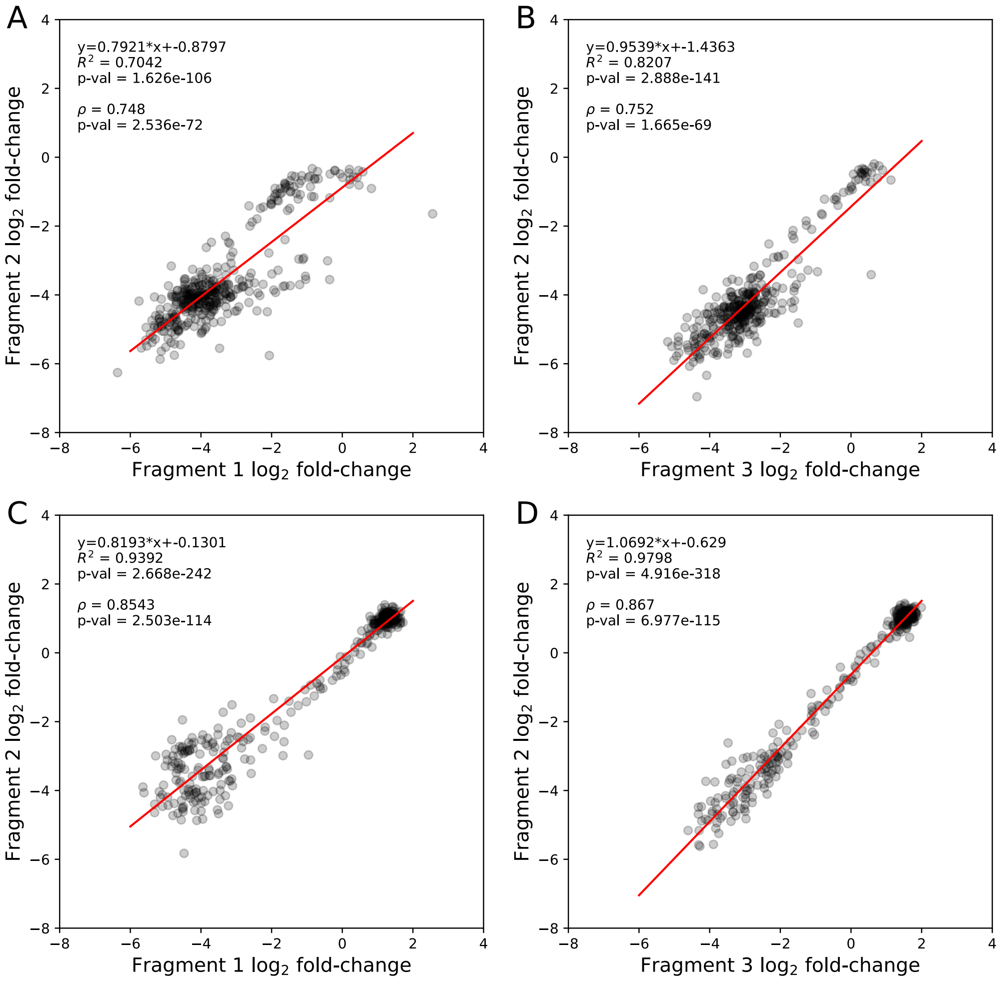


**Figure S6: Using pool overlaps to harmonise the $\log_2$fold-changes of the *FCY1* mutant pools.** For each panel, the linear regression parameters are shown, with Spearman's rank correlation shown below. **A)** Pool 1 to Pool 2, cytosine (n=397). **B)** Pool 3 to Pool 2, cytosine (n=375). **C)** Pool 1 to Pool 2, 5-FC (n=397). **D)** Pool 3 to Pool 2, 5-FC (n=375).

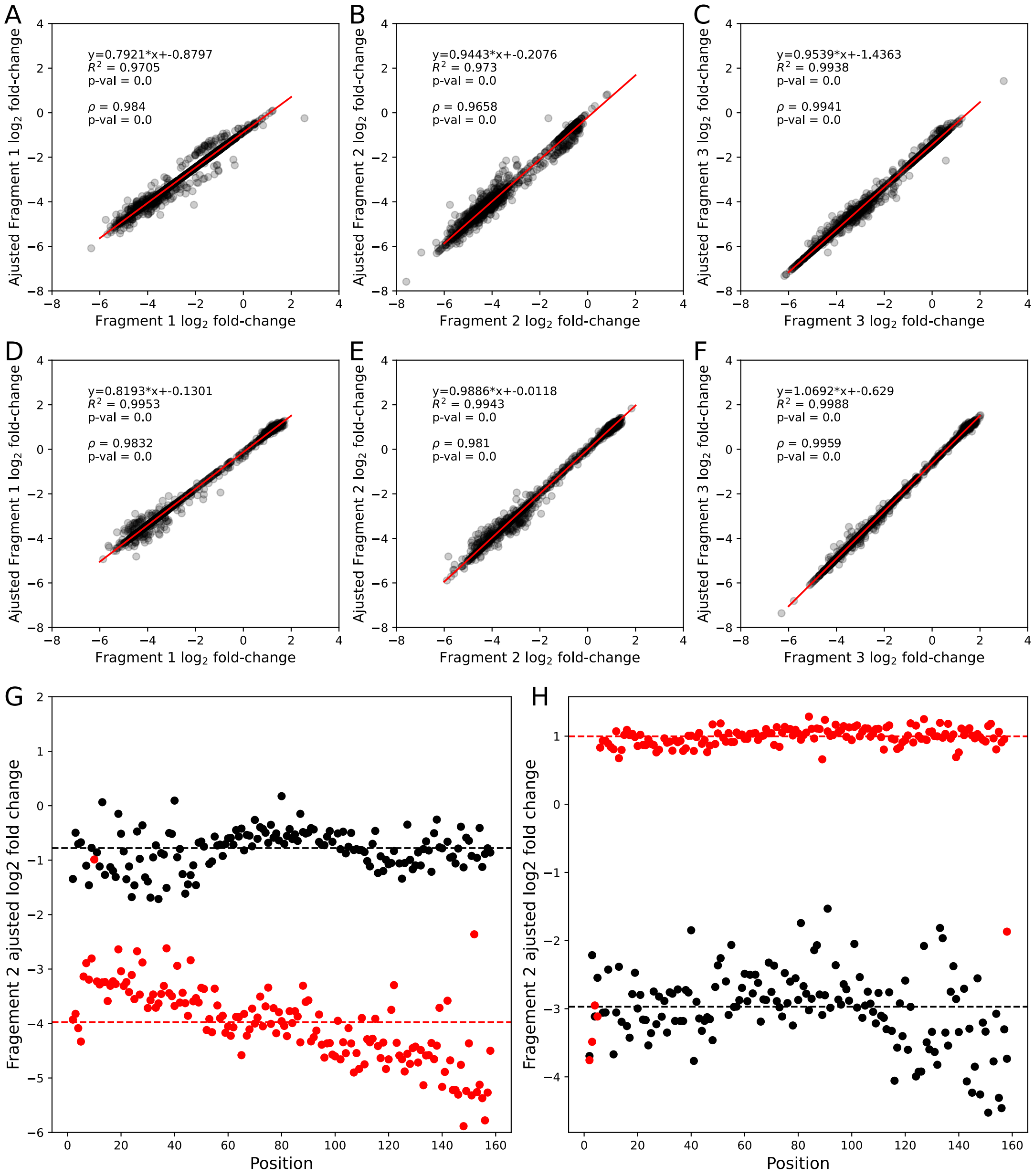


**Figure S7: Scaling log2 fold-changes to pool 2 and scaling scores to synonymous and nonsense mutations.** For panel A-F, the linear regression parameters are shown, with Spearman's rank correlation shown below for the comparison between raw pool scores and pool 2 adjusted scores. **A)** Raw pool 1 vs adjusted pool 1, cytosine (n=1372) **B)** Raw pool 2 vs adjusted pool 2, cytosine (n=1298). **C)** Raw pool 3 vs adjusted pool 3, cytosine (n=1374). **D)** Raw pool 1 vs adjusted pool 1, 5-FC (n=1372) **E)** Raw pool 2 vs adjusted pool 2, 5-FC (n=1298). **F)** Raw pool 3 vs adjusted pool 3, 5-FC (n=1374). **G)** Adjusted log2 fold-change for synonymous (black) and nonsense (red) mutants in cytosine (n=148 synonymous, n=156 nonsense mutants) along the protein positions. For positions where Met and Trp are the wild-type amino acids, there are no synonymous codons. This occurs 8 times in the *FCY1* coding sequence (not including the start Met). **H)** Adjusted log2 fold-change for synonymous (black) and nonsense (red) mutants in 5-FC (n=148 synonymous, n=156 nonsense mutants).

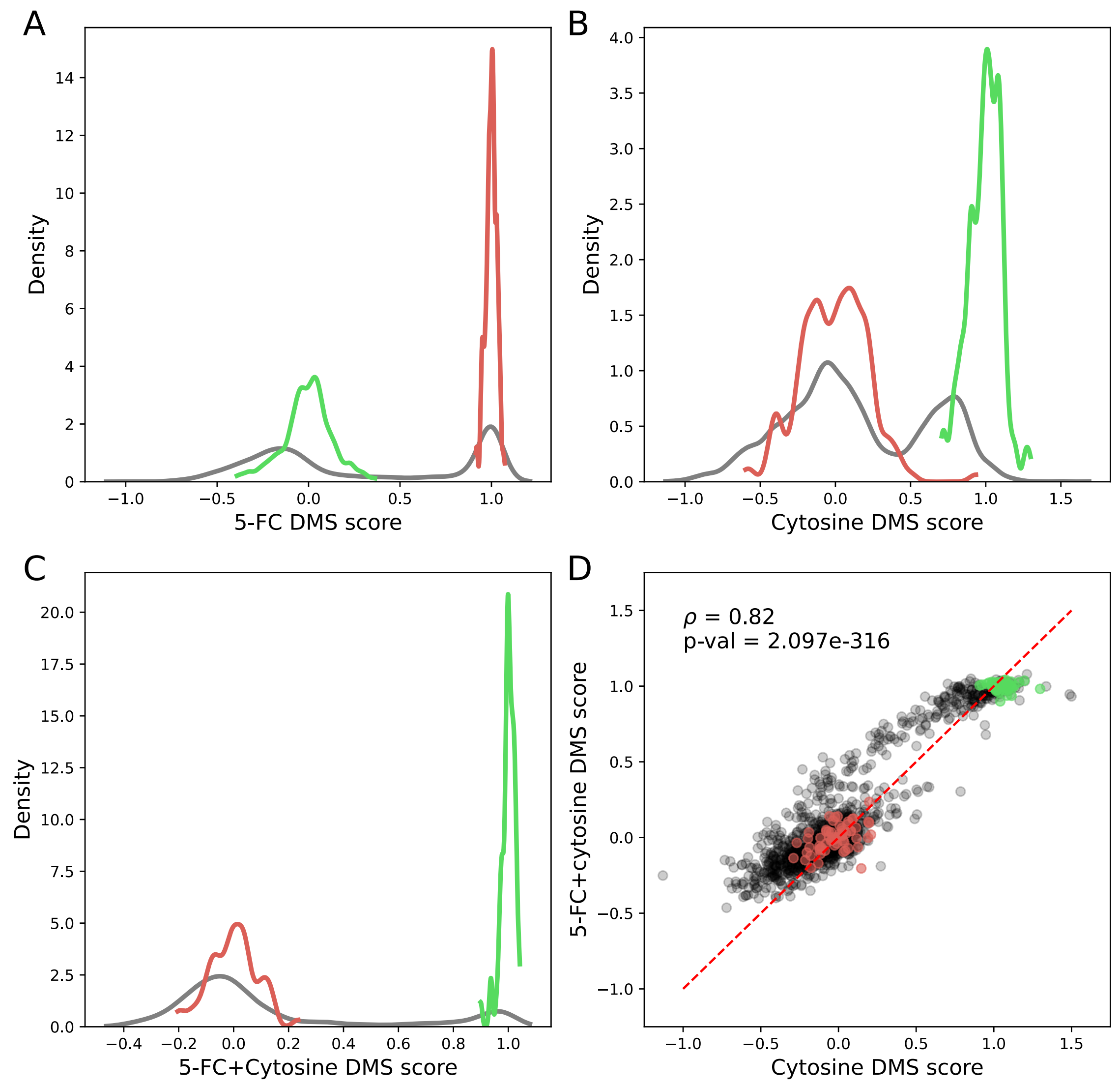


**Figure S8: DMS score distributions by mutant type.** Silent mutations are shown in green, nonsense in red, and missense mutants in grey. **A)** 5-FC (n=148 silent, 151 nonsense, 2968 missense). **B)** Cytosine (n=148 silent, 151 nonsense, 2968 missense) **C)** 5-FC + cytosine (n=59 silent, 62 nonsense, 1177 missense). **D)** DMS score in 5-FC + cytosine as a function of scores in cytosine only (n=1298 mutants).

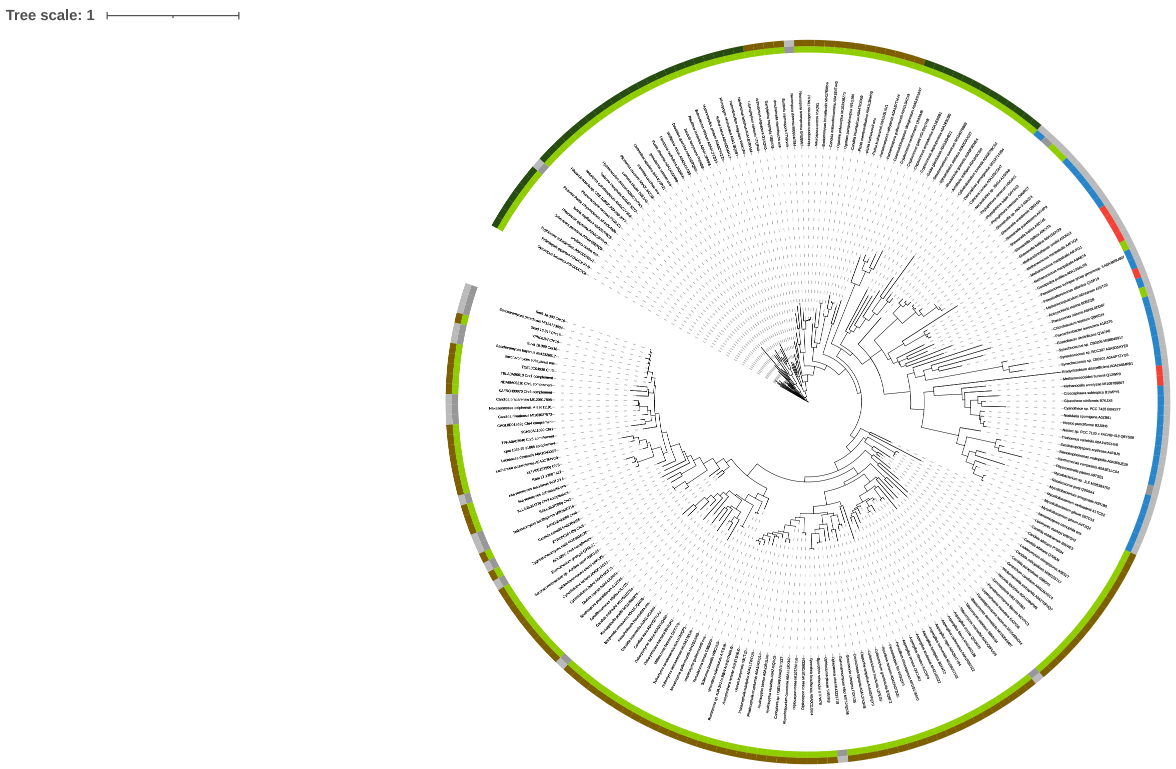


**Figure S9: Phylogenetic tree of *FCY1* orthologs used for calculating evolutionary rates.** The inner circle represents the kingdom (green: Eukarya, blue: Bacteria, red: Archea, grey: Missing) while the outer circle represents division within fungi (dark green: basidiomycota, brown: ascomycota). The tree was generated using the iTOL software[82]. The full size PDF file is available as Supplementary Data 4.

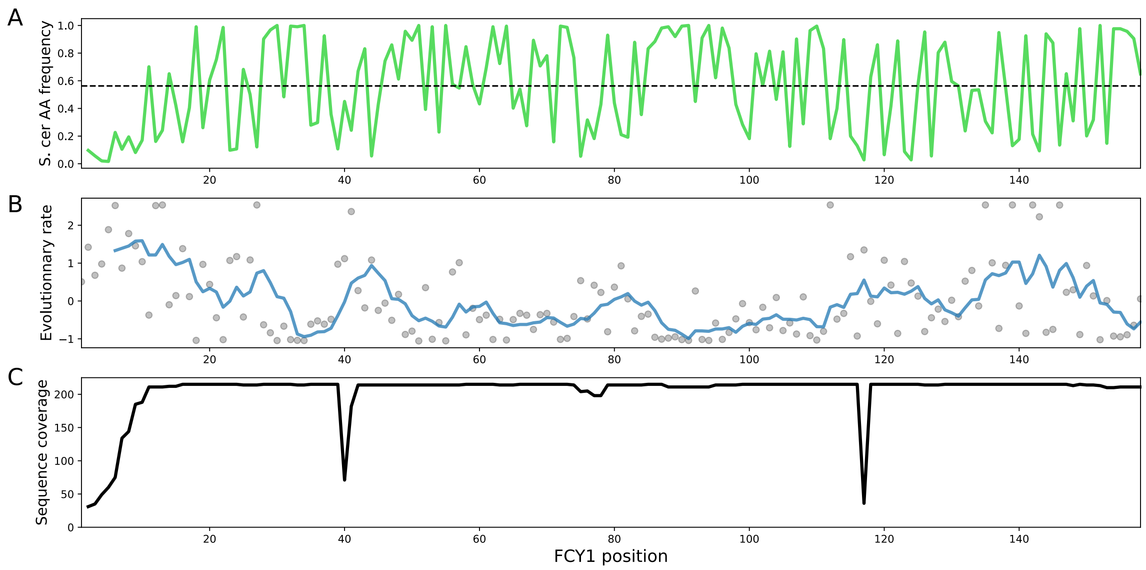


**Figure S10: *FCY1* evolutionary rate and orthologous residue diversity. A)** frequency of the *S. cerevisiae* amino acid at each position in the orthologous sequences. **B)** Normalized evolutionary rate for all *FCY1* residues (Rate4site[34]) from 215 orthologs. The blue line represents the rolling average over a 6 amino acid window. This statistic represents the rate at which amino acids change along a phylogeny: higher values represent more variable positions, while lower values represent highly conserved positions. **C)** Multiple sequence alignment coverage of *S. cerevisiae FCY1* positions in the set of orthologs. The maximum value is 215, representing perfectly conserved amino acids positions across all sequences.

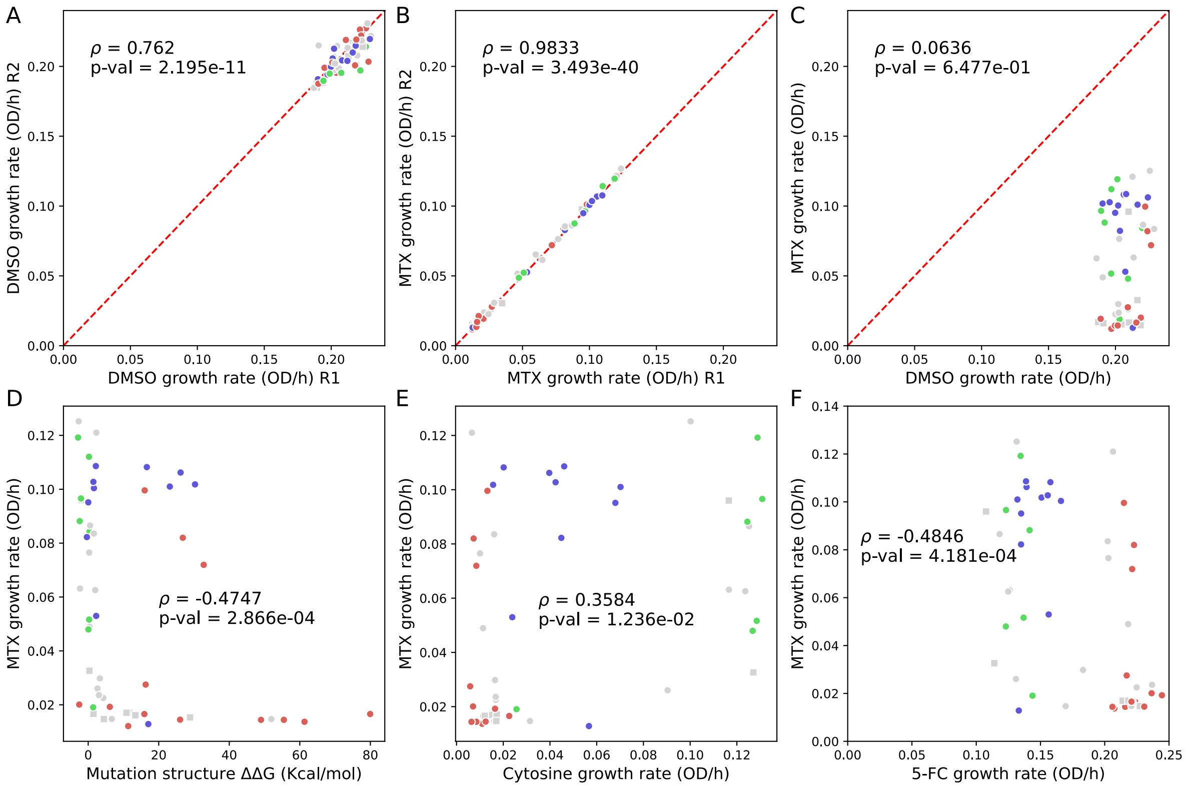


**Figure 11: DHFR-PCA data supports protein structure stability predictions for validation mutants.** *FCY1* variants were tagged with a DHFR-PCA[35] fragment to measure protein complex formation with a wild-type copy of *FCY1*.In this approach, Fcy1 is fused to DHFR fragments that complement upon dimerization allowing growth in media containing methotrexate (MTX). Growth reflects the amount of complex formed, therefore providing a quantitative measure of the stability of the Fcy1 subunits and complex. The 54 mutants are colored based on their DMS cluster (see Figure S13). **A)** Growth rates in DMSO of the validation mutants for the two replicates. DMSO is the MTX solvent and is the control condition for the DHFR-PCA assay. **B)** Growth rates in MTX of the validation mutants for the two replicates. **C)** Growth rate in MTX as a function of their growth rate in DMSO. As expected, there is no strong correlation between the two. **D)** Growth rate in MTX as a function of FoldX[32] predicted change in Fcy1 structure stability measured as ΔΔG. Positive ΔΔG represents detstabilization. **E)** Growth rate in MTX as a function of the growth rate in cytosine media of the haploid strain. **F)** Growth rate in MTX and in 5-FC media of the haploid strain.

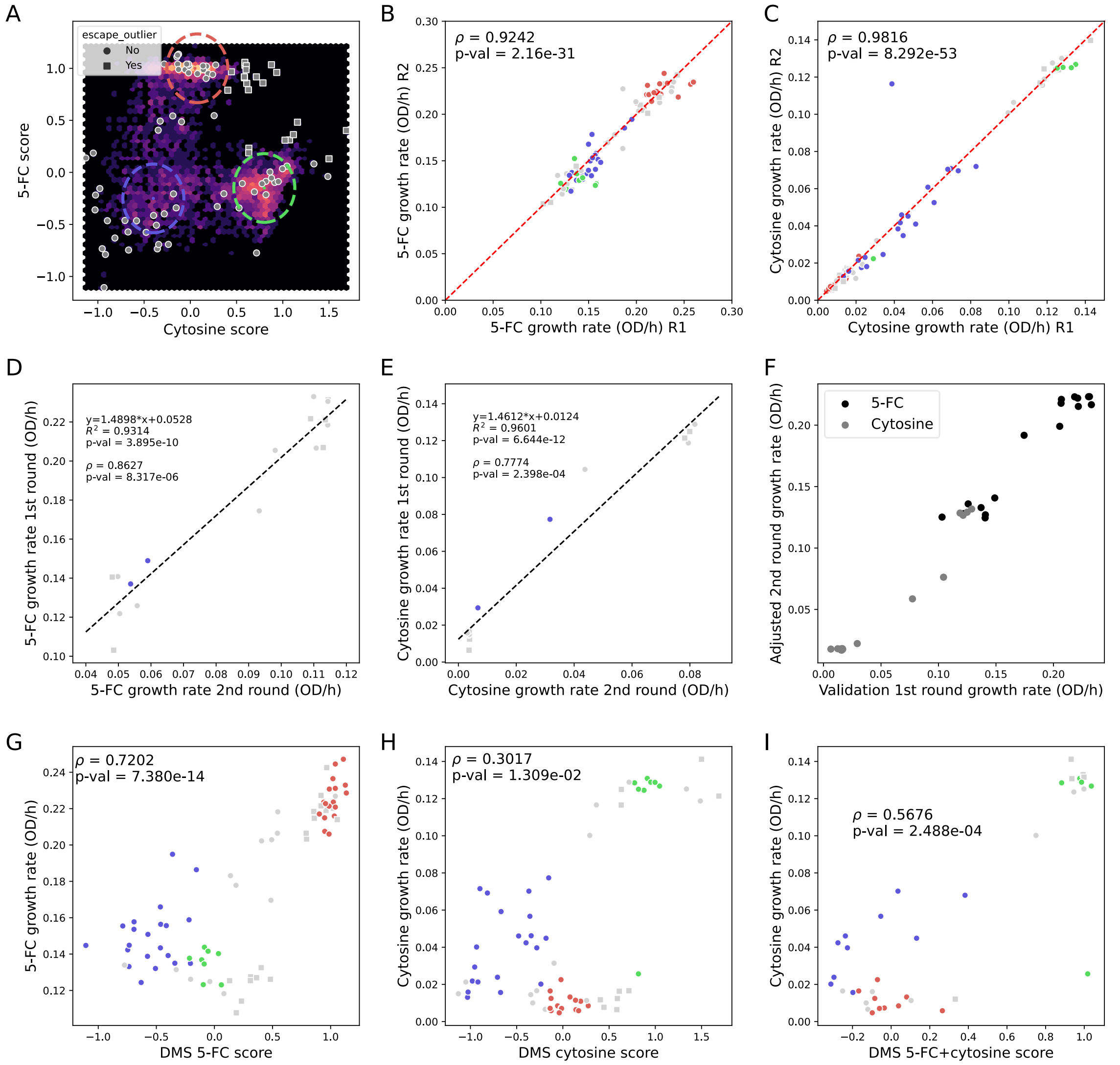


**Figure S12: DMS score validations in cytosine and 5-FC media. A)** Location on the cytosine/5-FC landscape of the Fcy1 variants (shown as grey dots) selected for validations superimposed on the density plot presented in Figure 2D. The circles used to define the three clusters are also shown: green for silent-like mutants, red for nonsense-like mutants, and blue for front minimum mutants. Mutants falling outside these clusters were classified as ‘other’ and encompass most outliers from the DMS screen. Variants with both high 5-FC and cytosine DMS scores are shown as squares: these outliers potentially escape the resistance-function trade-off. **B)** Spearman’s correlation between growth curve replicates in 5-FC media, n=73 variants. **C)** Spearman’s correlation between growth curve replicates in cytosine media, n=72 variants. Data collected from the cytosine media from the outlier (T86M) was excluded from downstream analysis. **D)** Linear regression fit and Spearman’s correlation between 5-FC growth rates measured in the 1st and 2nd rounds of validations for 17 mutants present in both growth curve assays. **E)** Linear regression fit and Spearman’s correlation between cytosine growth rates measured in the 1st and 2nd rounds of validations for 17 mutants present in both assays. **F)** Growth rate values of the 2nd round of validations scaled to the growth rate of the 1st round for the 17 mutants present in both assays. **G)** Spearman’s correlation between validation growth curves in 5-FC media and DMS 5-FC score, n=79 variants. **H)** Spearman’s correlation between validation growth curves in cytosine media and DMS cytosine score, n=79 variants. **I)** Spearman’s correlation between validation growth curves in cytosine media and DMS 5-FC + cytosine score, n=38 variants.

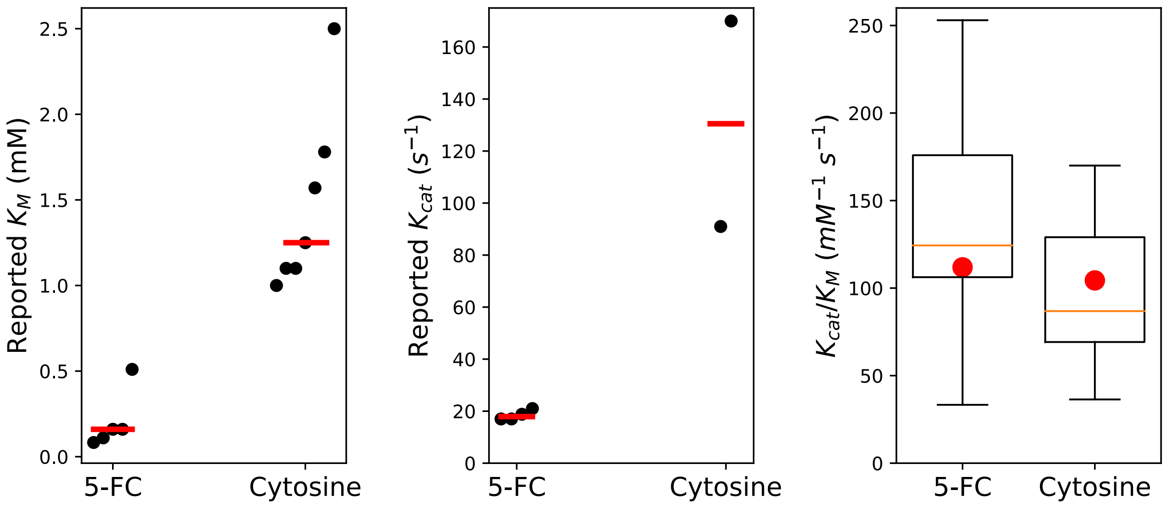


**Figure S13: Reported $K_M$ and $K_{cat}$ values for 5-FC and cytosine lead to similar distributions of catalytic efficiencies.** For $K_M$ and $K_{cat}$, the red line represents the median reported value. For the $K_M/K_{cat}$ distribution, the red dot represents the value obtained for the median $K_{cat}$/median $K_M$. Data are from BRENDA[37] (EC 3.5.4.1).

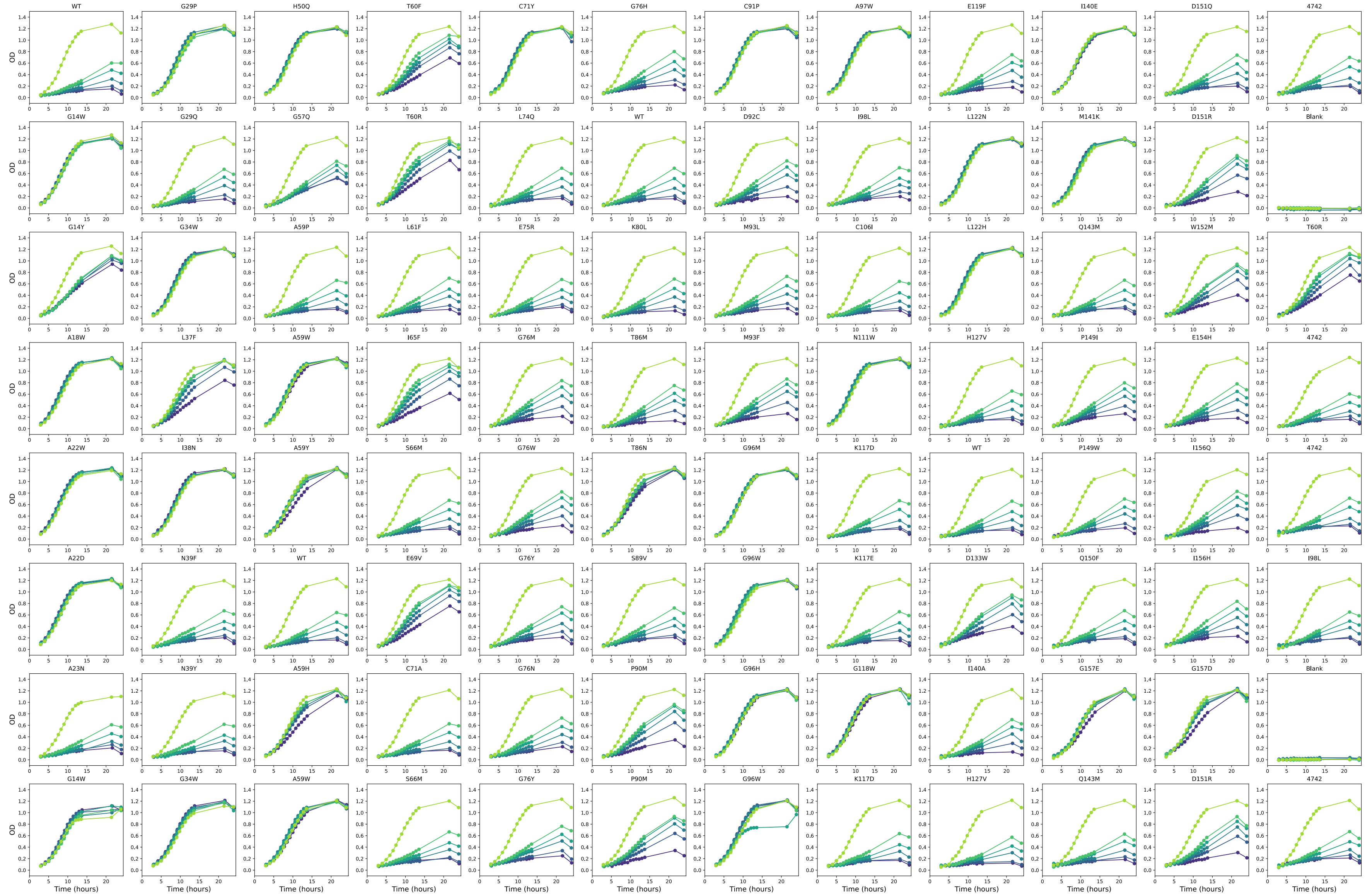


**Figure S14: Individual growth curve measurements for the validation mutants in various 5-FC concentrations.** Curves are colored as a function of the increasing 5-FC concentration, from light to dark green/blue: 0 μM, 24 μM,48 μM, 97 μM, 194 μM, 387μM.

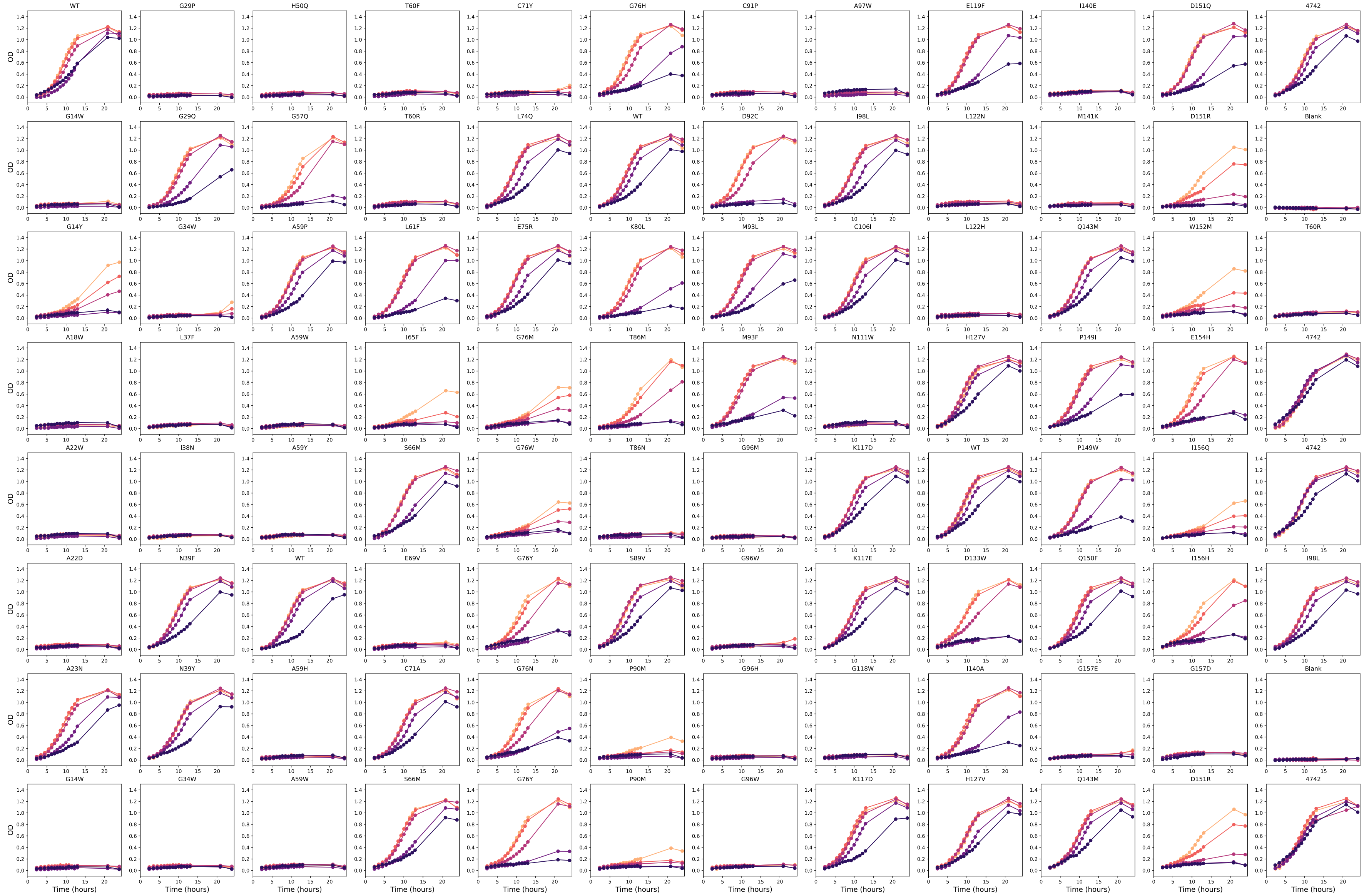


**Figure S15: Individual growth curve measurements for validation mutants in various cytosine concentrations.** Curves are colored as a function of the decreasing cytosine concentration, from light to dark orange/purple: 90 μM, 113 μM, 225 μM, 450 μM, 901 μM.

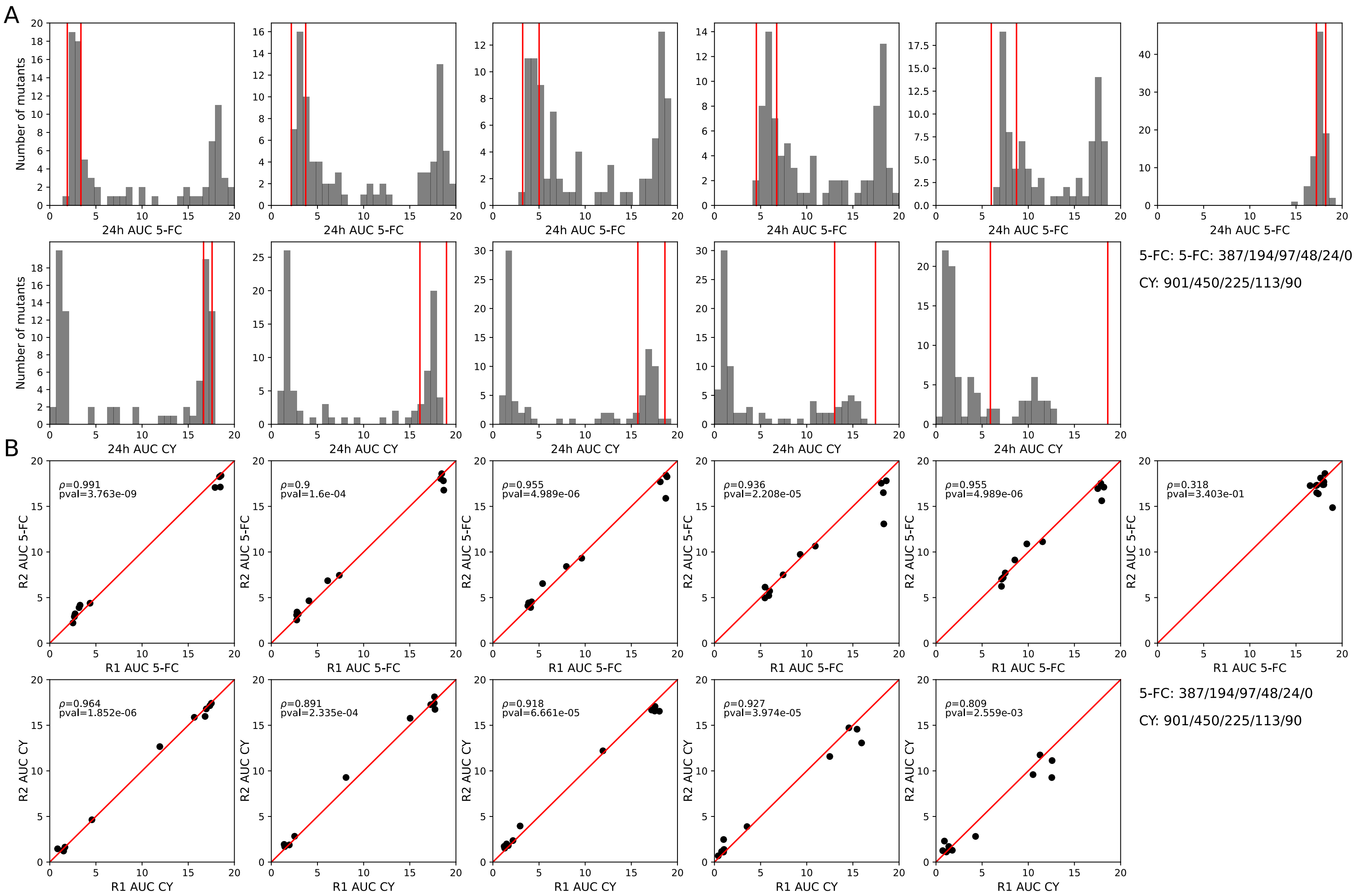


**Figure S16: Dose-response area under the curves (AUC) values and replication. A)** AUC distribution for all variants in the different 5-FC and cytosine plates. The red lines represent 3 absolute median deviations around the median of the wild-type and BY4742 replicates. **B)** AUC values of both replicates for mutants present in duplicates for all 5-FC and cytosine conditions (n=13).

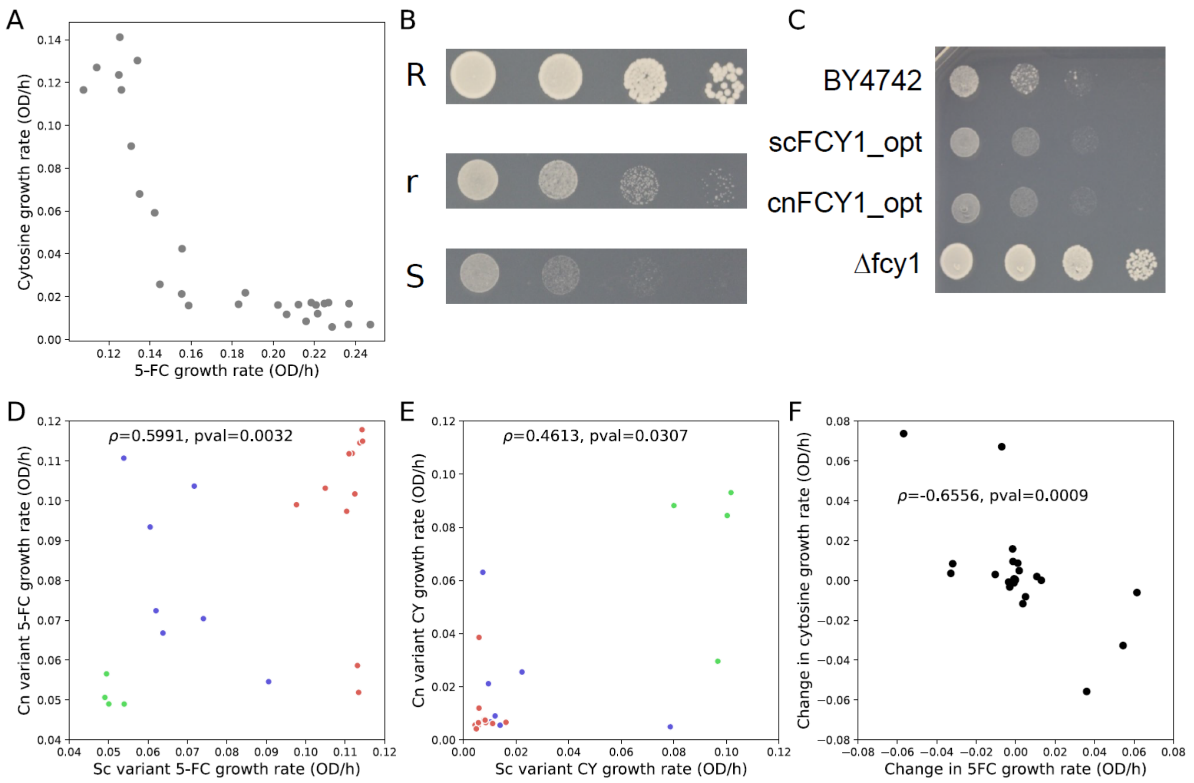


**Figure S17: Phenotypes of the *scFCY1* and *cnFCY1* mutants . A)** Growth rate for the 29 (27 of which were successfully constructed) *scFCY1* variants selected in the DMS assay validation experiments. **B)** Representative examples of the phenotypes observed in the spot assays (Synthetic media + 194 μM 5-FC, 10-fold dilutions starting at 1 OD/ml), where S: sensitive, r: low growth and R: full resistance. **C)** Phenotypes of scFCY1_opt ( *S. cerevisiae* codon optimised *FCY1* at the FCY1 locus), cnFCY1_opt (C. neoformans codon optimised FCY1 at the *S. cerevisiae* FCY1 locus) compared to the parental strain (BY4742) and the deletion mutant (*Δfcy1*). **D)** Comparison of growth rate between orthologous variants in 5-FC media. Spearman's rank correlation is shown (n=22 pairs). Variants are colored by the position along the trade-off of the *scFCY1* variant. **E)** Comparison of 5-FC media growth rate between orthologous variants. Spearman's rank correlation is shown (n=22 pairs). **F)** Changes in growth rate in SC + 12 μM 5-FC and SC-Ura + 84 μM cytosine between *scFCY1* and *cnFCY1* variants. Spearman's rank correlation is shown (n=22 pairs).

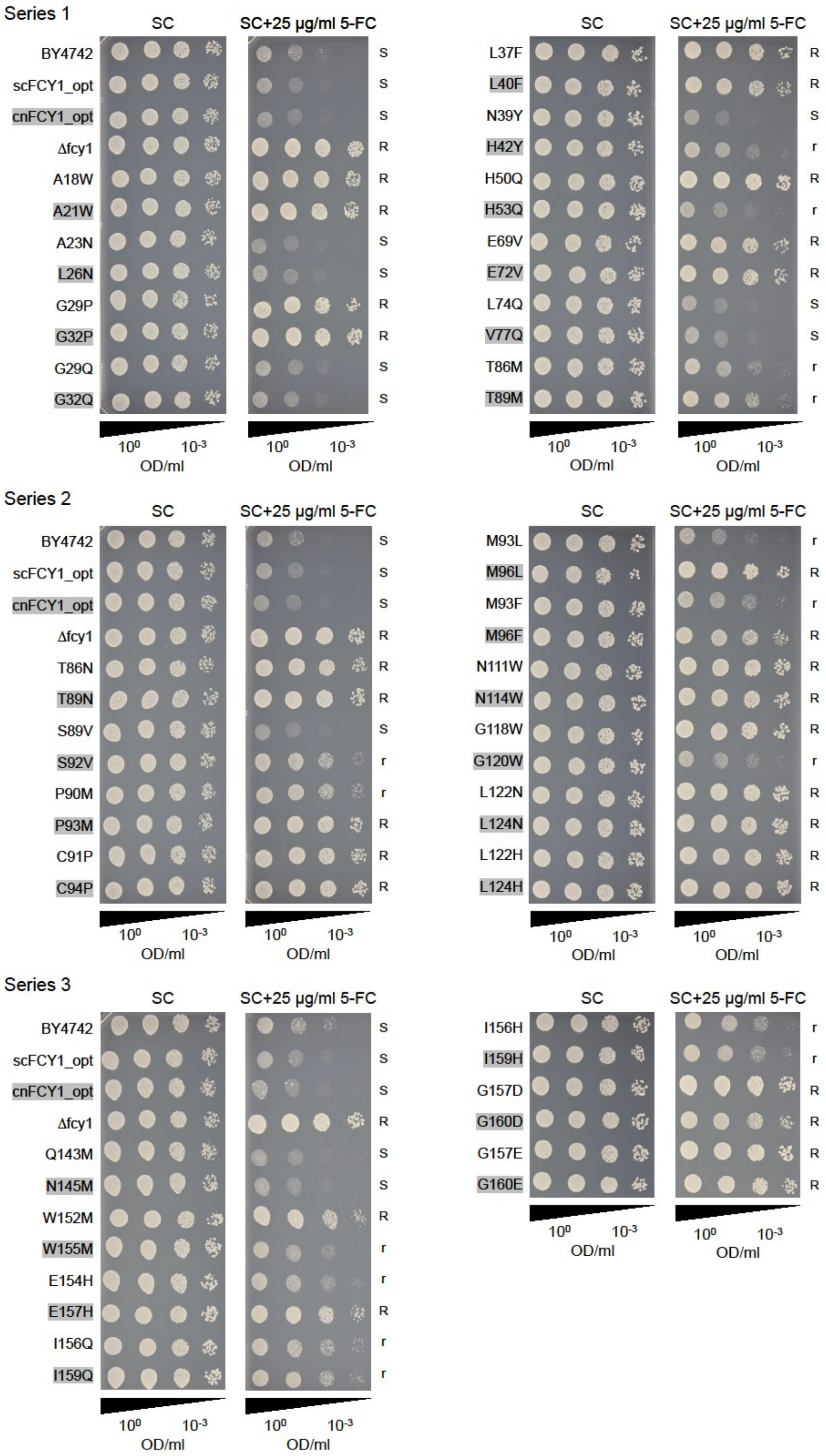


**Figure S18: Spot dilution assay phenotypes are most often conserved between orthologous mutants of *scFCY1* and *cnFCY1*.** The same dilutions of control strains BY4742 (WT *FCY1*), scFCY1_opt (*S. cerevisiae* codon optimised *FCY1* at the FCY1 locus), cnFCY1_opt (C. neoformans codon optimised FCY1 at the *S. cerevisiae* FCY1 locus) and *Δfcy1* were spotted on each plate. For each mutant pair, the scFCY1 strain is in white and the cnFCY1 is highlighted in grey. The phenotype score (as defined earlier) for each strain is shown on the right. The raw images used to generate the figure are available as Supplementary Data 5.
