## Supplementary figures and images for "Asymmetrical dose-responses shape the evolutionary trade-off between antifungal resistance and nutrient use"

### 01_WT_YPD.JPG

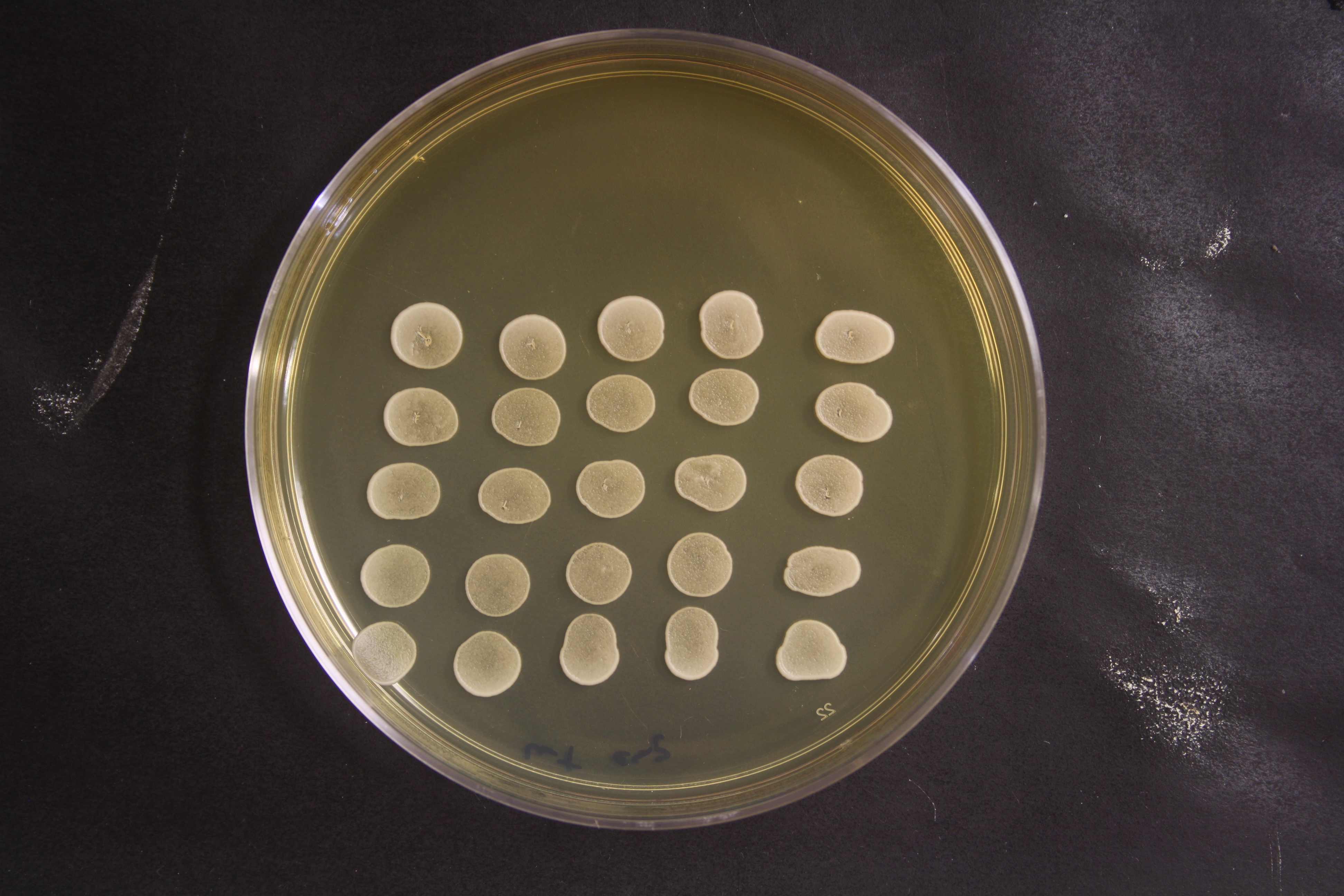

### 02_Delta_YPD.JPG

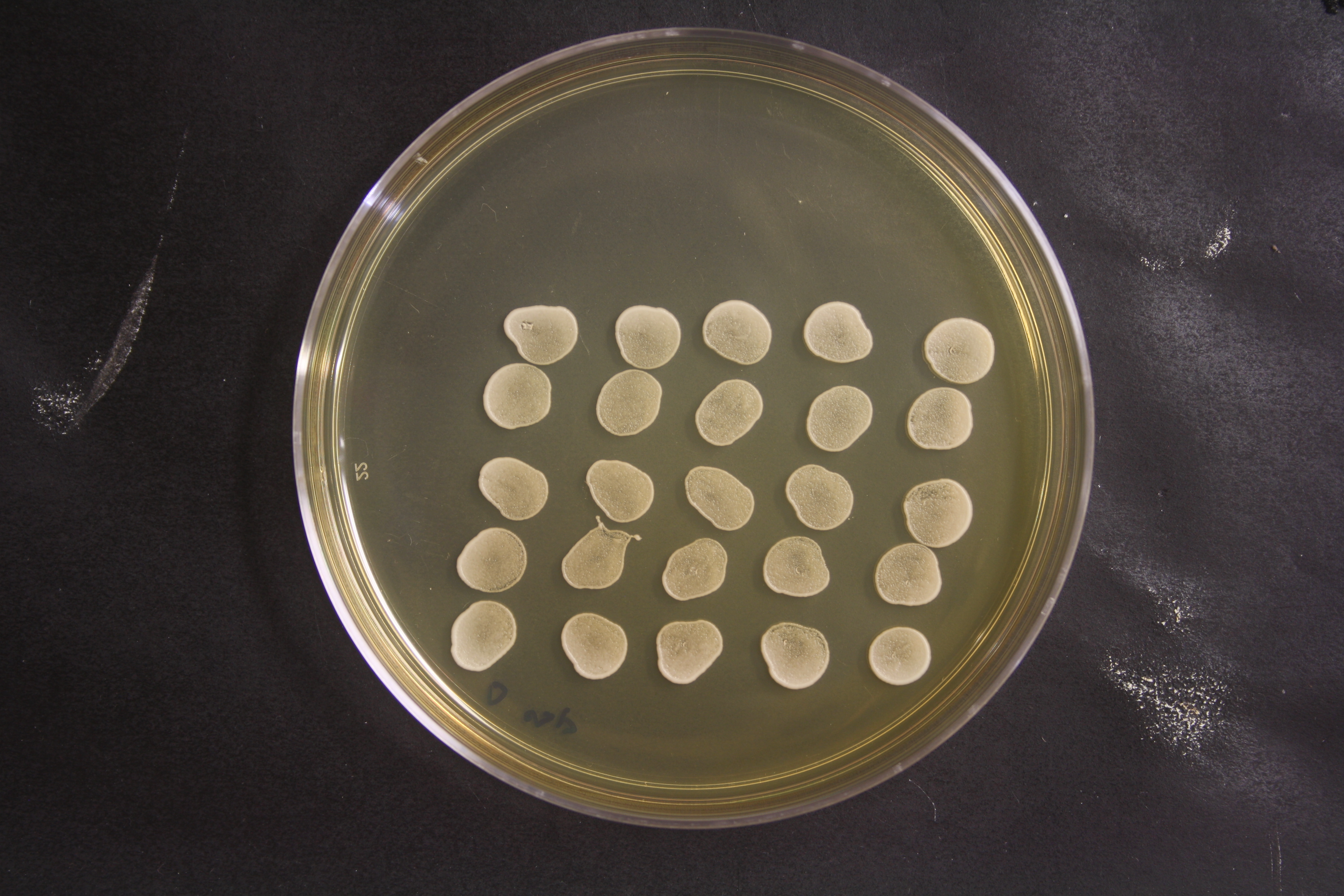

### 03_Grid_YPD.JPG

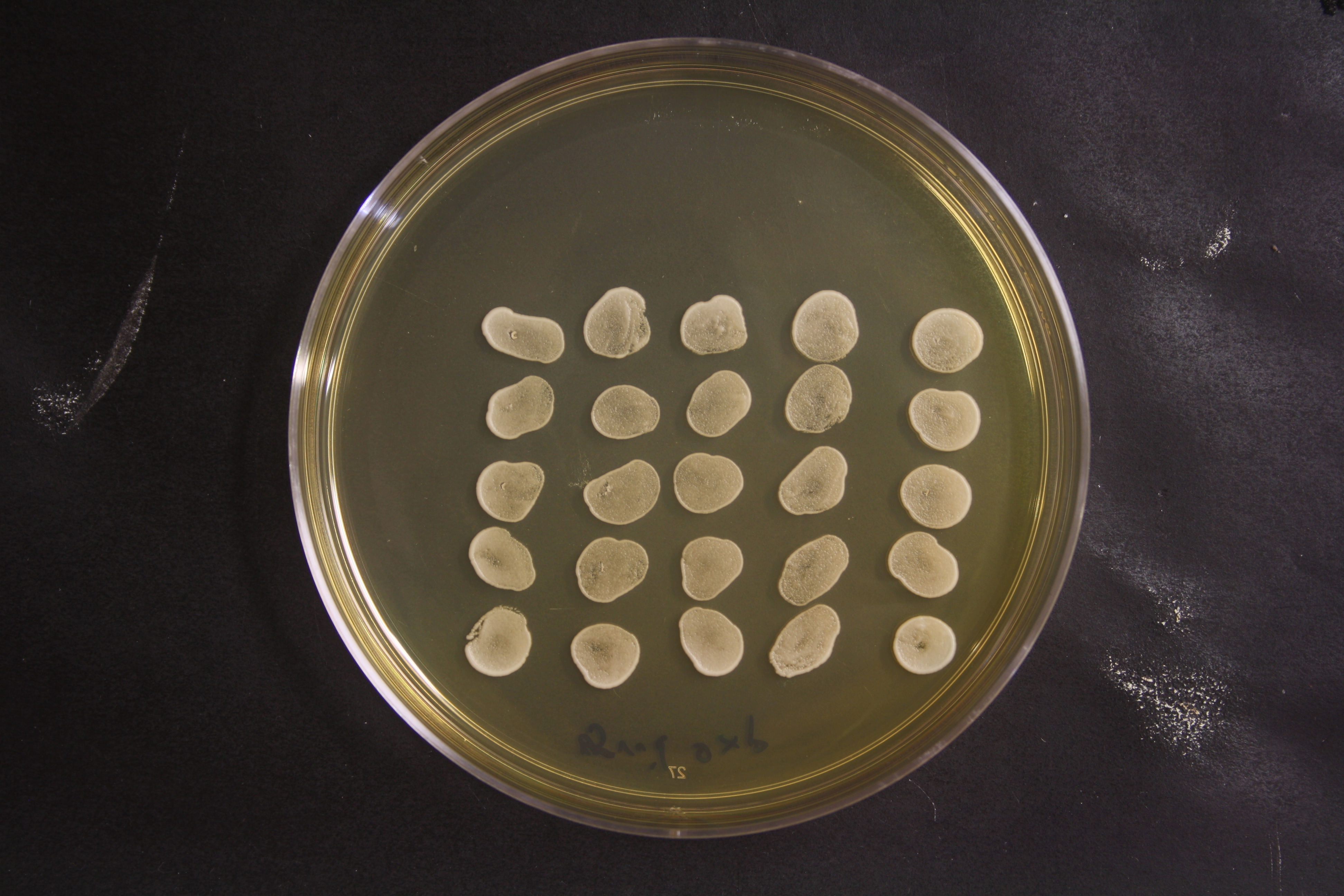

### 04_WT_5FC.JPG

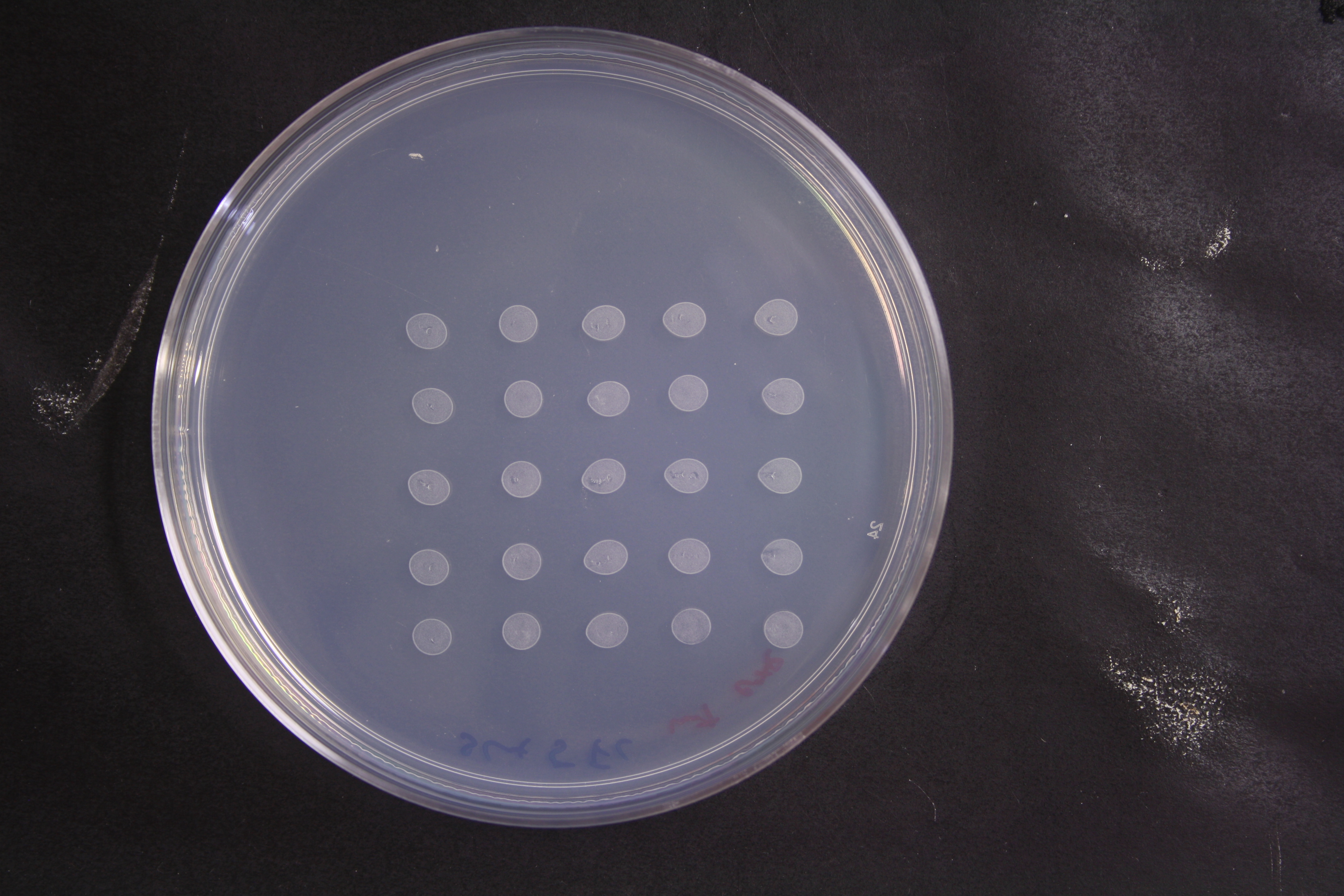

### 05_Delta_5FC.JPG

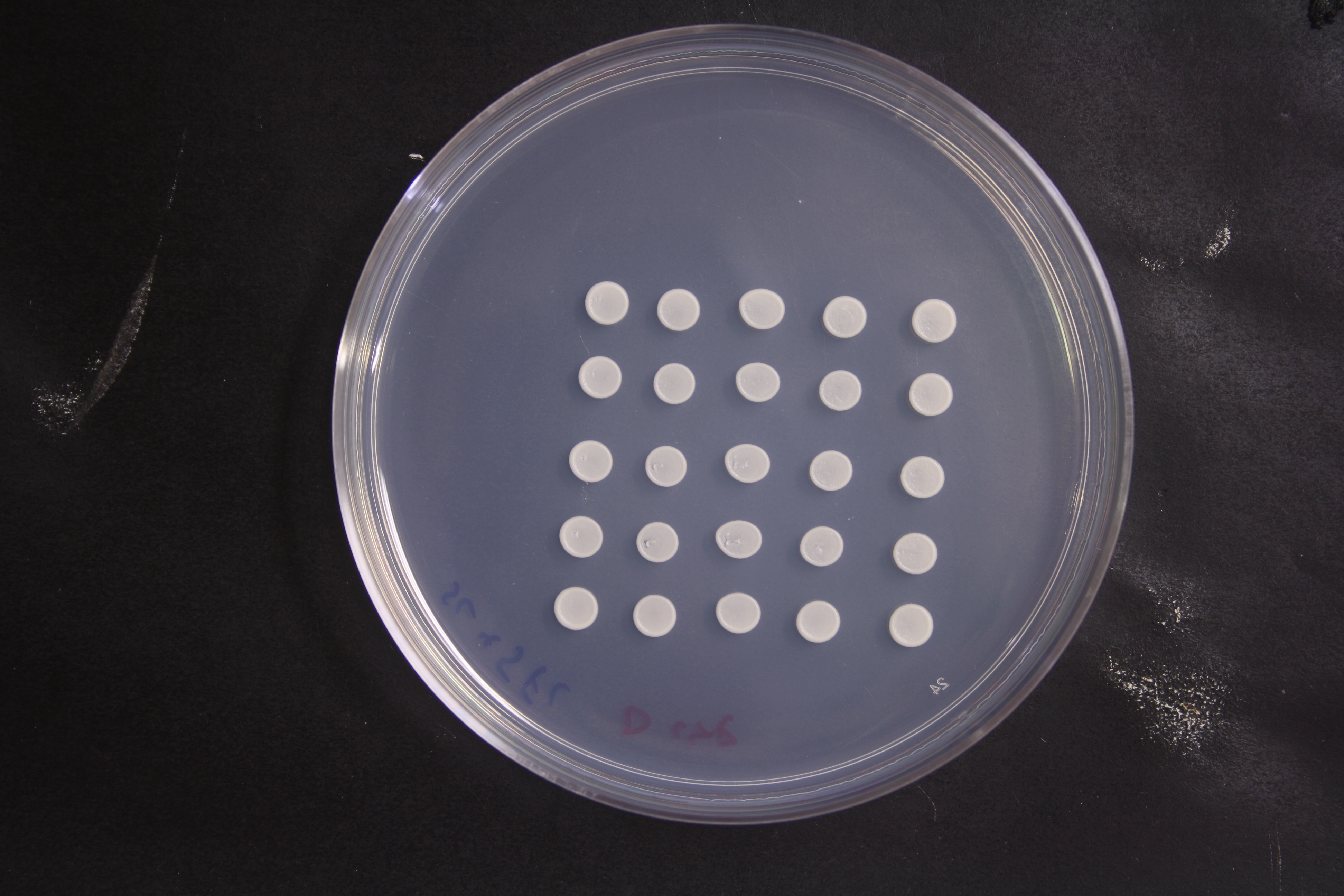

### 06_Grid_5FC.JPG

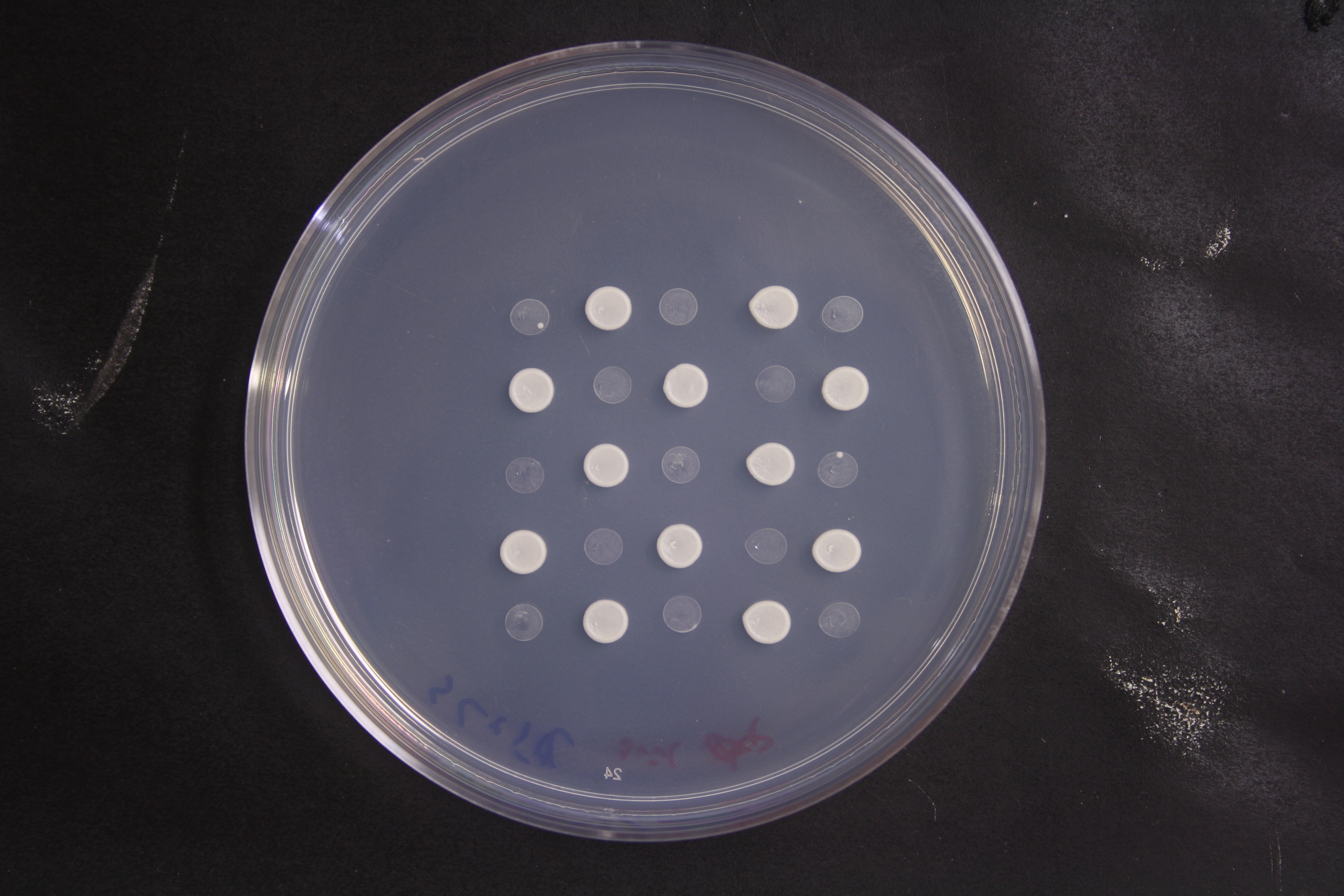

### 07_WT_Cytosine.JPG

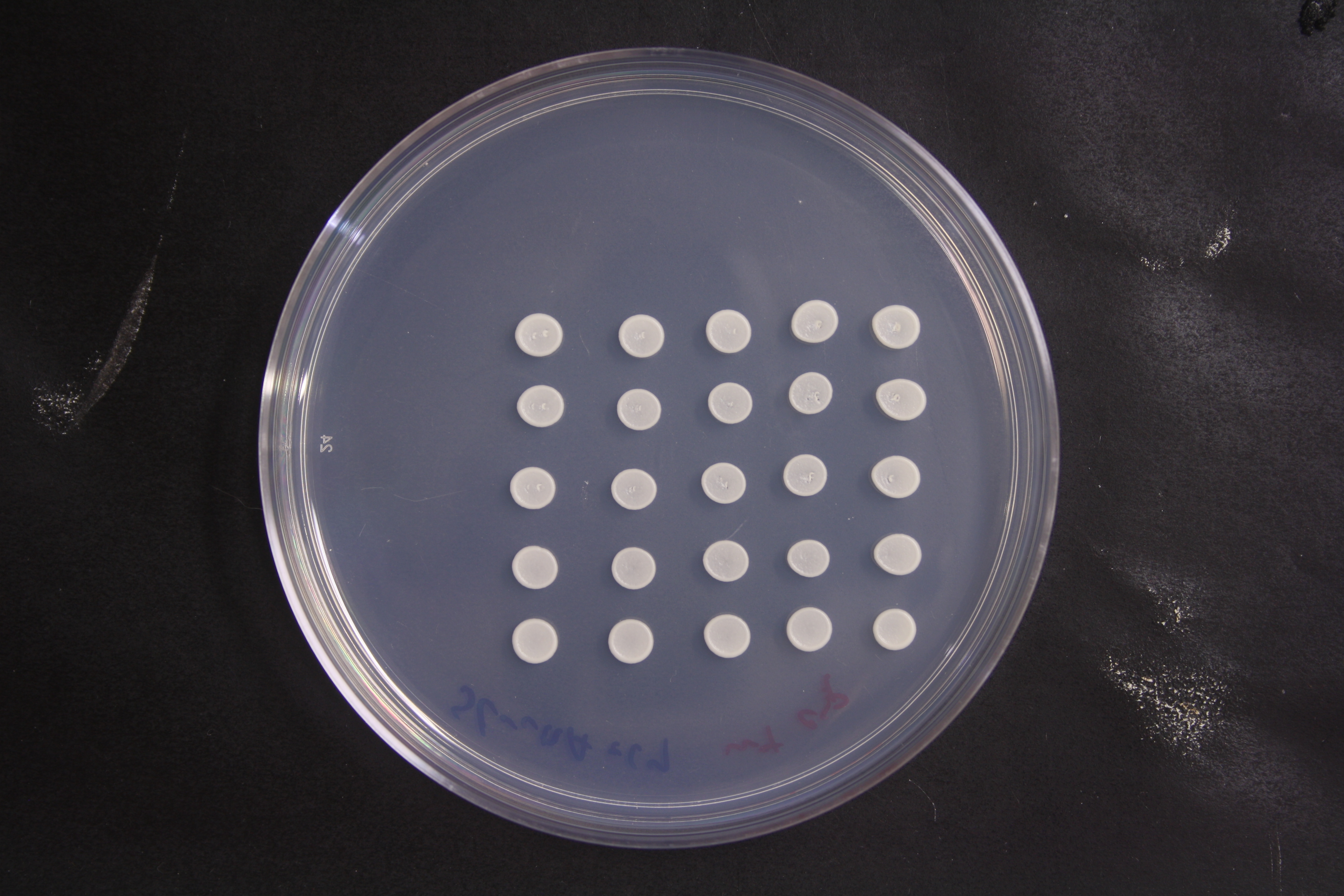

### 08_Delta_Cytosine.JPG

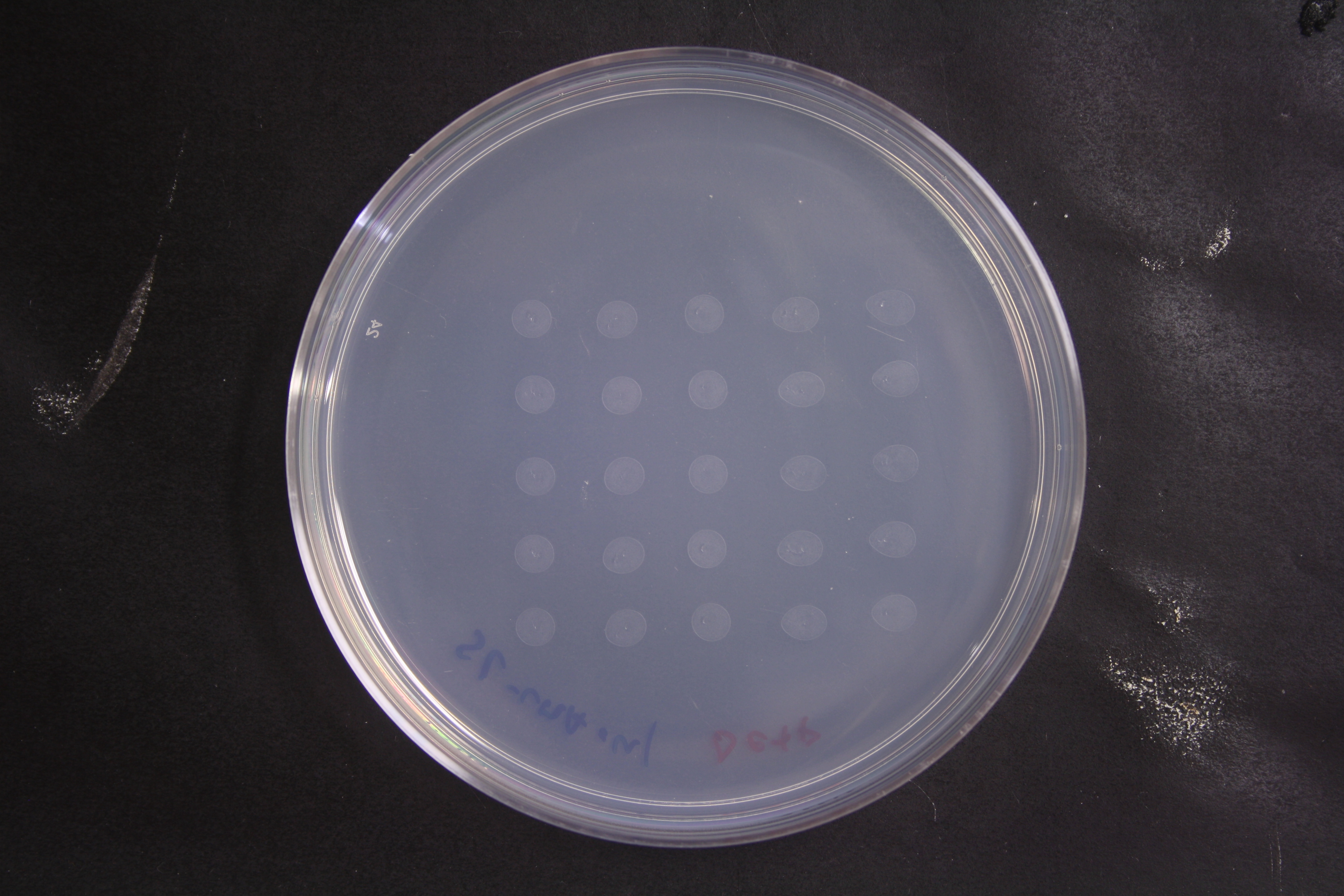

### 09_Grid_Cytosine.JPG

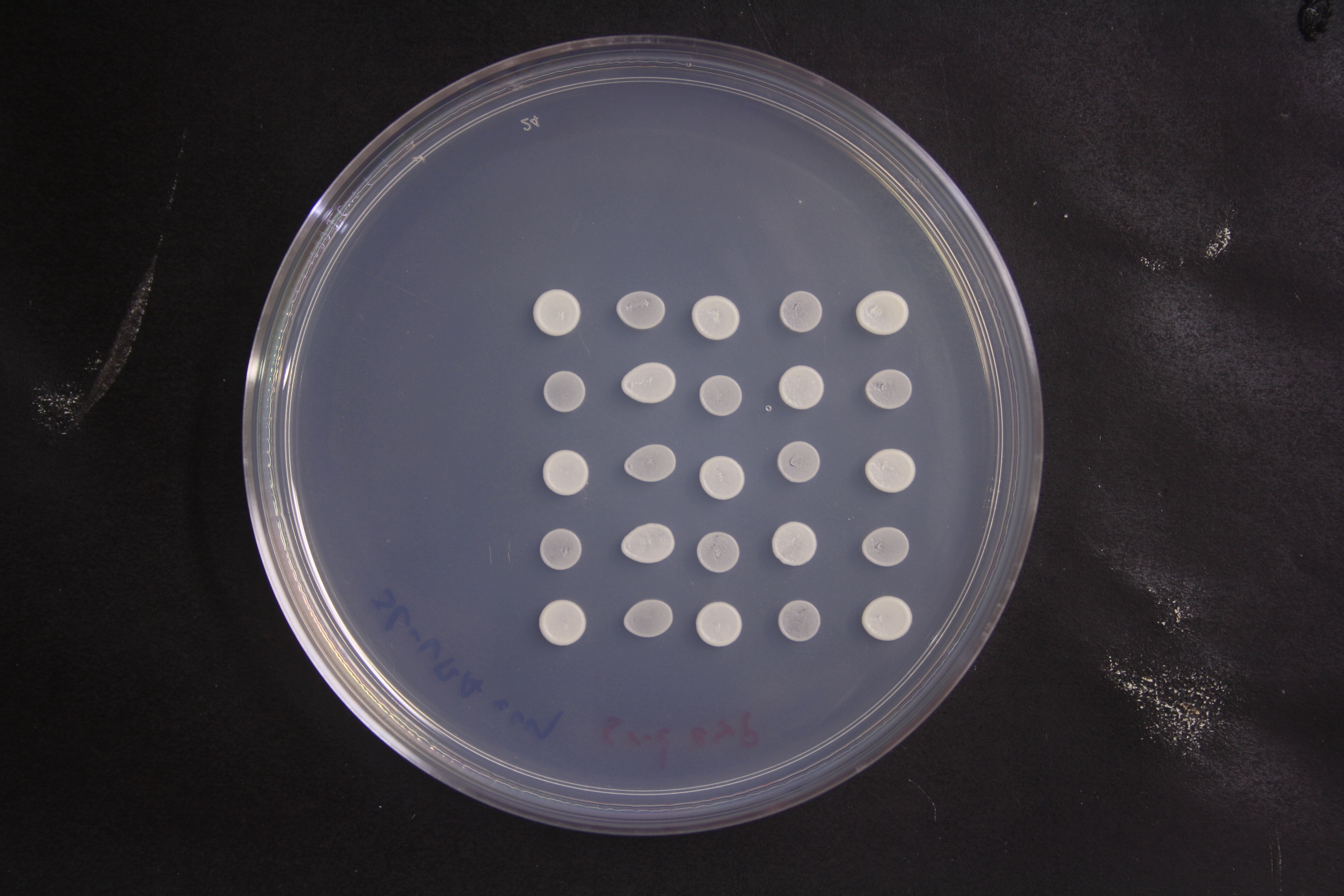

### FCY1_c_neoformans_65444_coverage.png

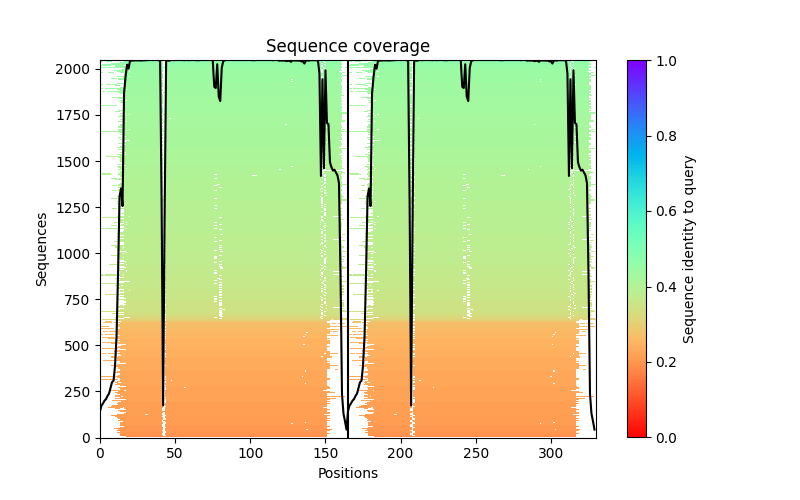

### FCY1_c_neoformans_65444_PAE.png

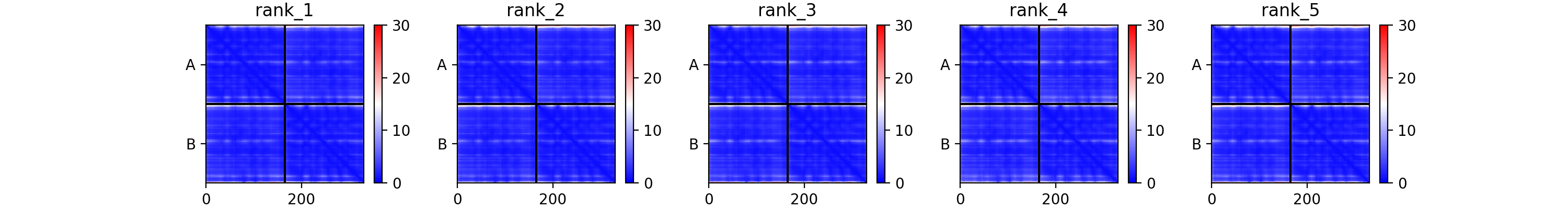

### FCY1_c_neoformans_65444_plddt.png

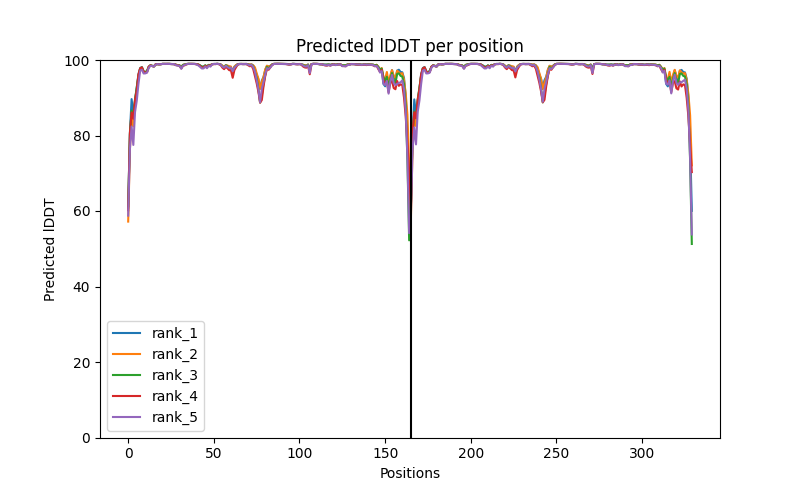

### plate_1_SC_5FC.JPG

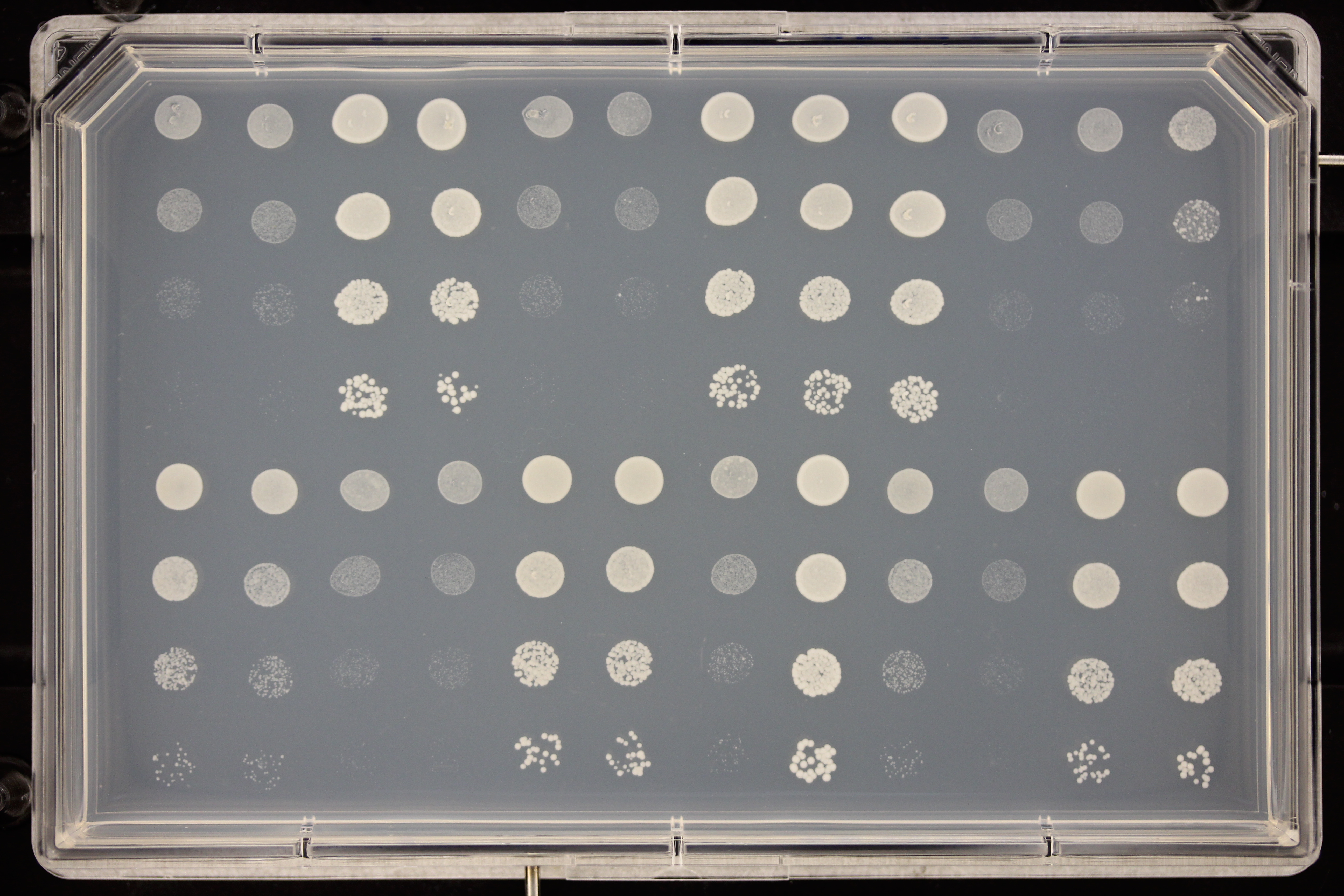

### plate_2_SC.JPG

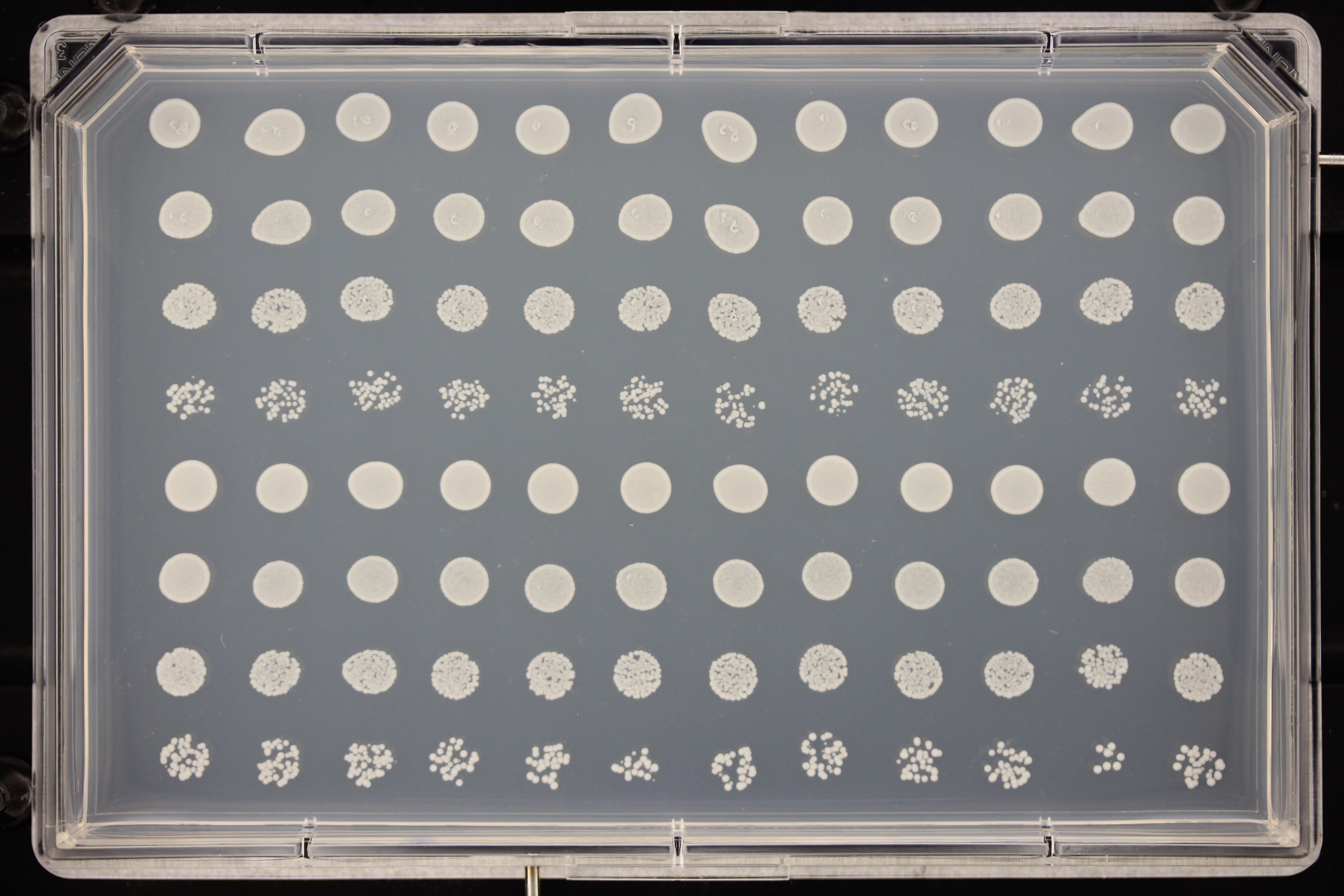

### plate_2_SC_5FC.JPG

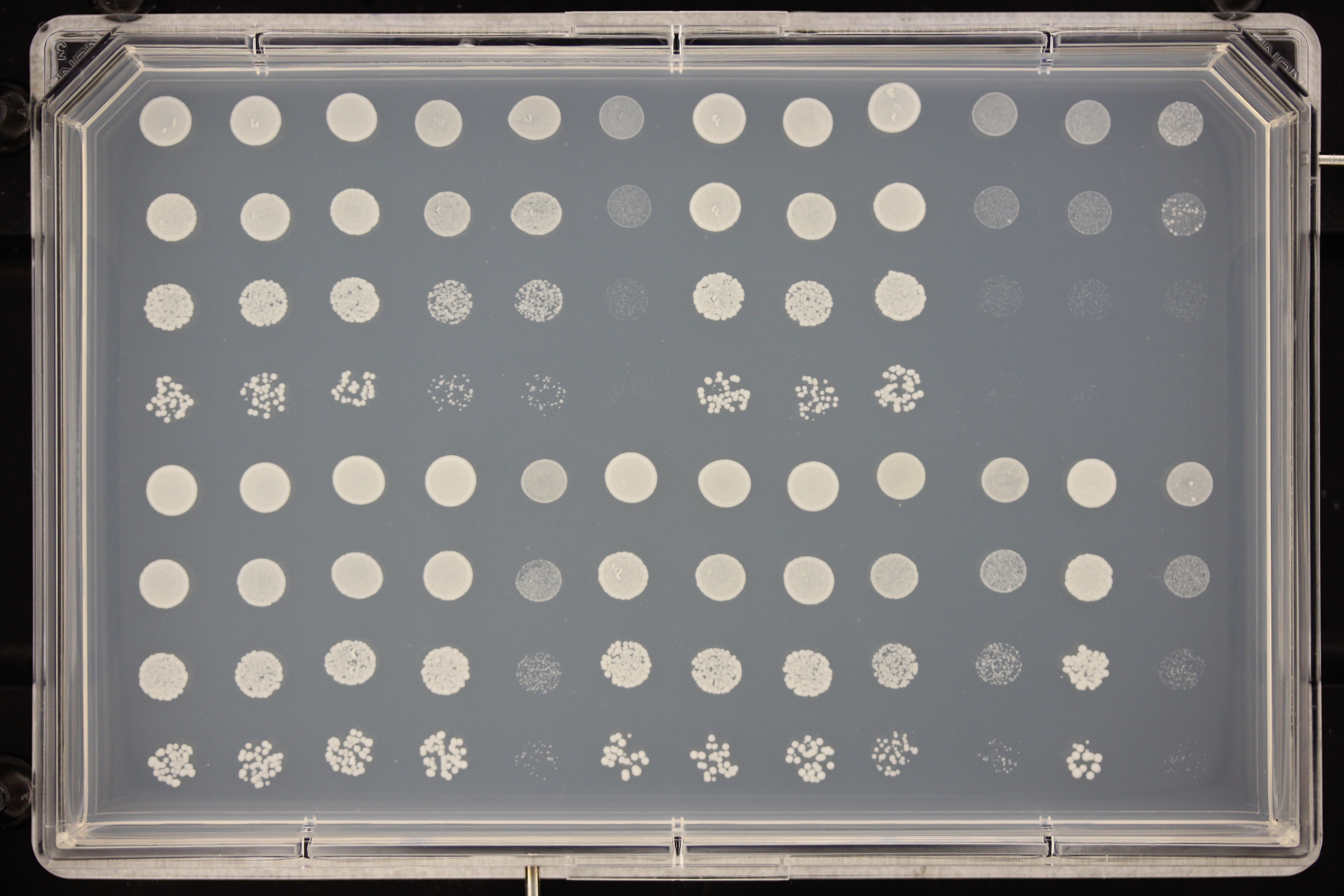

### plate_3_SC.JPG

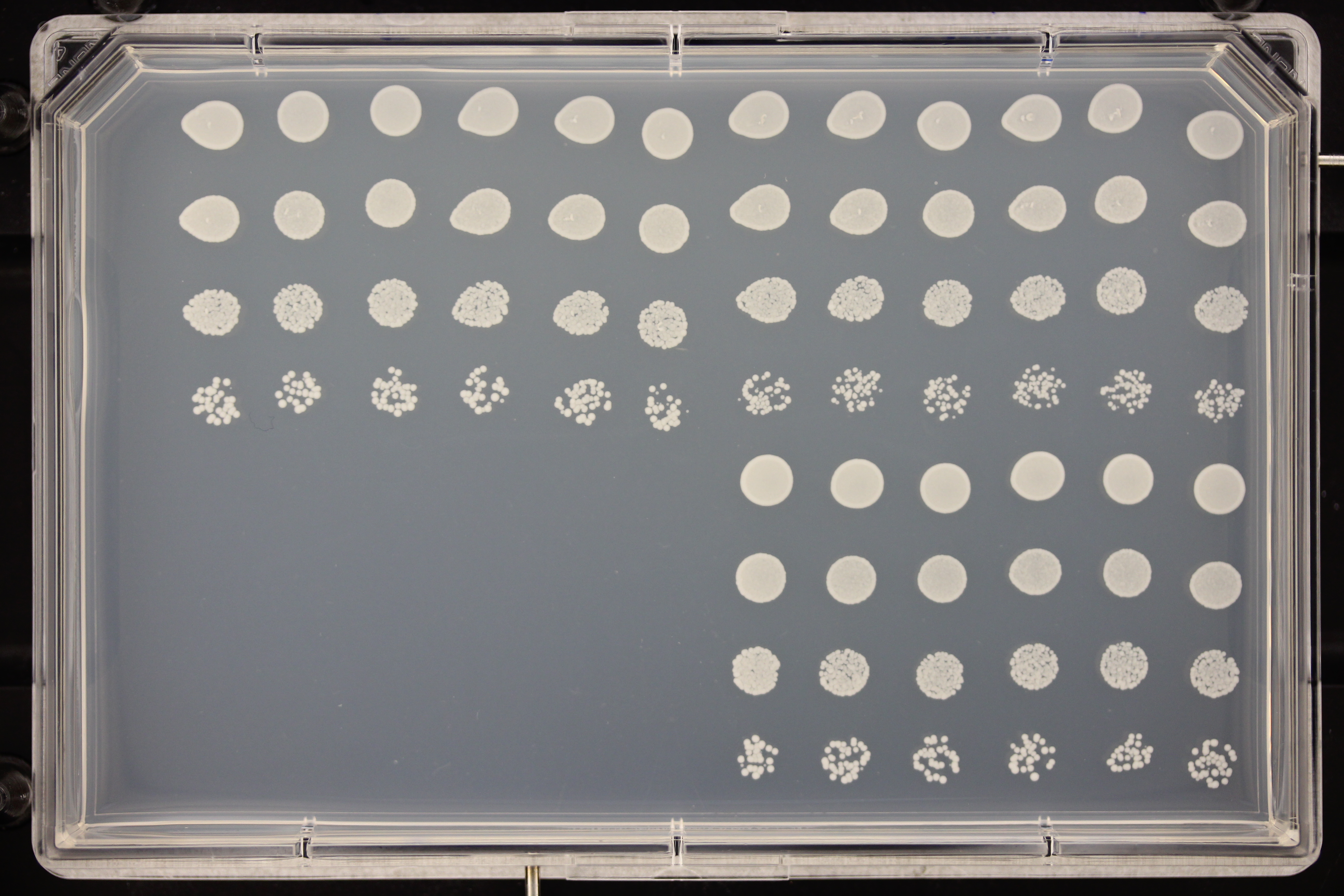

### plate_3_SC_5FC.JPG

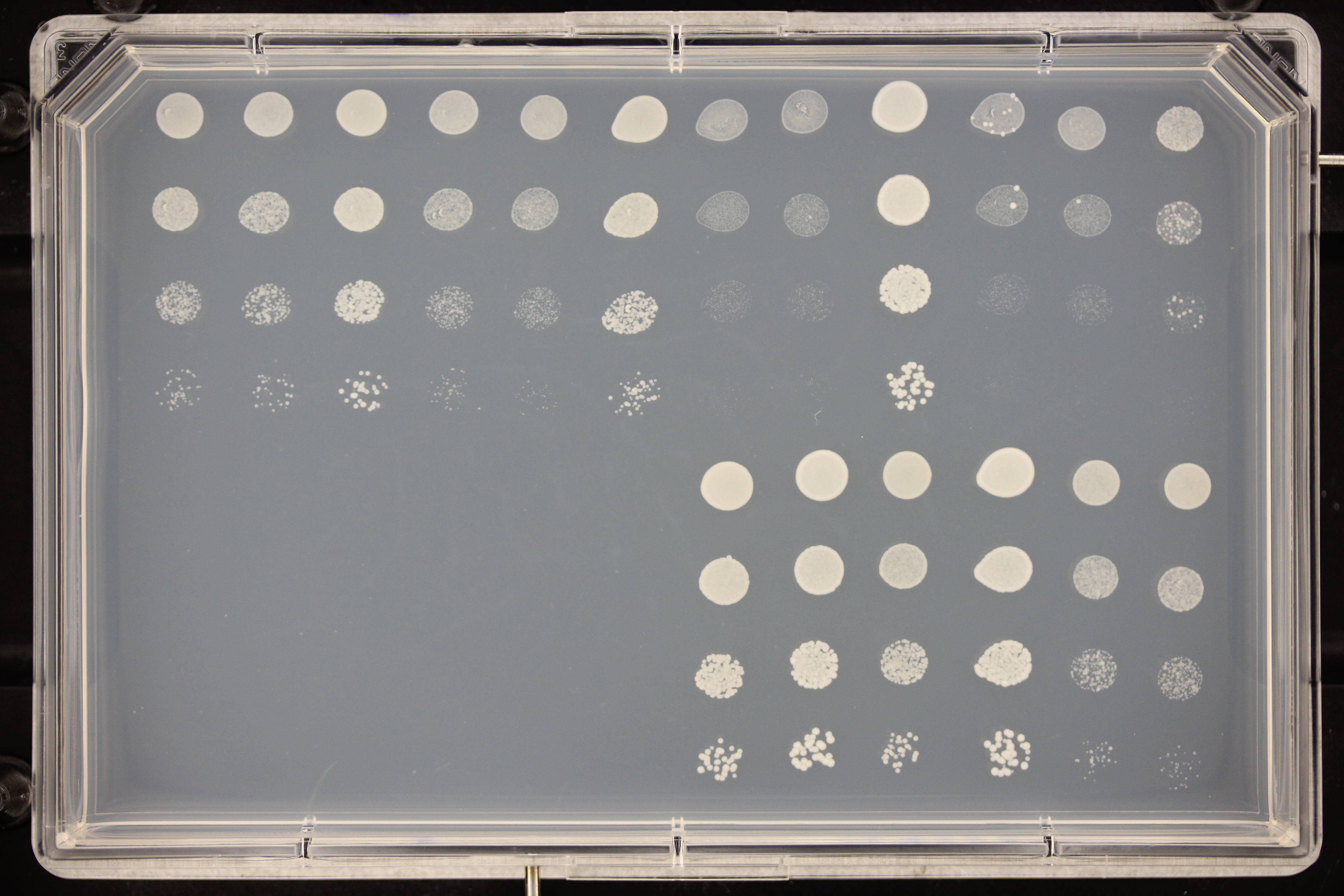

### Supplementary_data_4.pdf

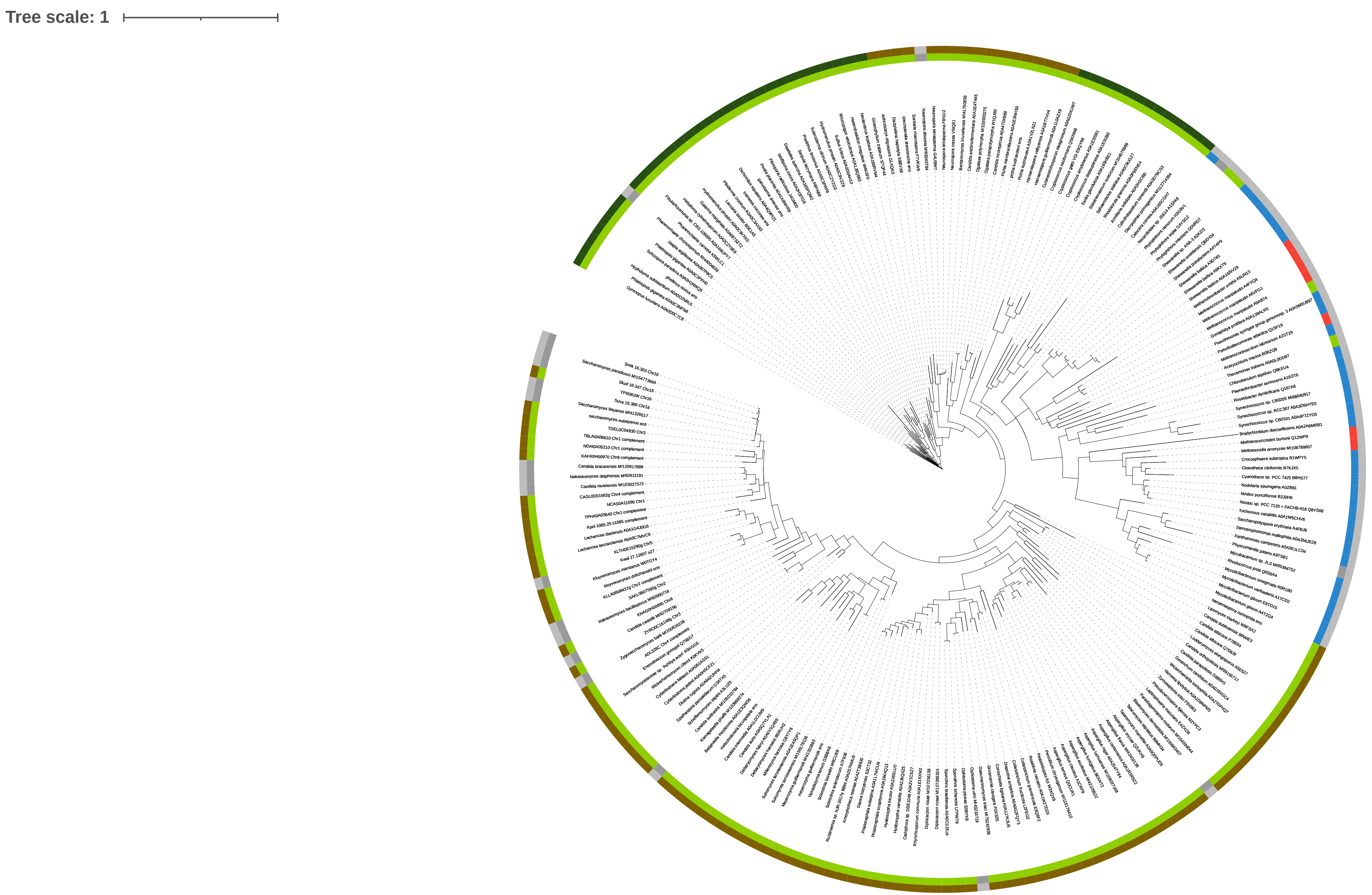
