## Supplementary tables 1-11 for "Asymmetrical dose-responses shape the evolutionary trade-off between antifungal resistance and nutrient use": S_table_7.readme.txt

#Description of dataframe columns in table S6. The is one row for each DMS (position x coding variant) possibility.

codon: the position in WT Saccharomyces cerevisiae Fcy1p
wt: the wild-type amino acid at this position in Saccharomyces cerevisiae
measured: the DMS variant
CY_1: Measured log2 fold-change of variant in fragment 1, cytosine
CY_1_R1: Cytosine pool 1, replicate 1 amino acid log2 fold-change
CY_1_MAD_1: Cytosine pool 1, replicate 1 codon log2 fold-change median absolute deviation for amino acid
CY_1_R2: Cytosine pool 1, replicate 2 amino acid log2 fold-change
CY_1_MAD_2: Cytosine pool 1, replicate 2 codon log2 fold-change median absolute deviation for amino acid
FC_1: Measured log2 fold-change of variant in fragment 1, 5-FC
FC_1_R1: 5-FC pool 1, replicate 1 amino acid log2 fold-change
FC_1_MAD_1: 5-FC pool 1, replicate 1 codon log2 fold-change median absolute deviation for amino acid
FC_1_R2: 5-FC pool 1, replicate 2 amino acid log2 fold-change
FC_1_MAD_2: 5-FC pool 1, replicate 2 codon log2 fold-change median absolute deviation for amino acid
CY_2: Measured log2 fold-change of variant in fragment 2, cytosine,
CY_2_R1: Cytosine pool 2, replicate 1 amino acid log2 fold-change
CY_2_MAD_1: Cytosine pool 2, replicate 1 codon log2 fold-change median absolute deviation for amino acid
CY_2_R2: Cytosine pool 2, replicate 2 amino acid log2 fold-change
CY_2_MAD_2: Cytosine pool 2, replicate 2 codon log2 fold-change median absolute deviation for amino acid
FC_2: Measured log2 fold-change of variant in fragment 2, 5-FC
FC_2_R1: 5-FC pool 2, replicate 1 amino acid log2 fold-change
FC_2_MAD_1: 5-FC pool 2, replicate 1 codon log2 fold-change median absolute deviation for amino acid
FC_2_R2: 5-FC pool 2, replicate 2 amino acid log2 fold-change
FC_2_MAD_2: 5-FC pool 2, replicate 2 codon log2 fold-change median absolute deviation for amino acid
FC_CY_2: Measured log2 fold-change of variant in fragment 2, 5-FC + cytosine
FC_CY_2_R1: 5-FC+cytosine pool 2, replicate 1 amino acid log2 fold-change
FC_CY_2_MAD_1: 5-FC+cytosine pool 2, replicate 1 codon log2 fold-change median absolute deviation for amino acid
FC_CY_2_R2: 5-FC+cytosine pool 2, replicate 2 amino acid log2 fold-change
FC_CY_2_MAD_2: 5-FC+cytosine pool 2, replicate 2 codon log2 fold-change median absolute deviation for amino acid
CY_3: Measured log2 fold-change of variant in fragment 3, cytosine
CY_3_R1: Cytosine pool 3, replicate 1 amino acid log2 fold-change
CY_3_MAD_1: Cytosine pool 3, replicate 1 codon log2 fold-change median absolute deviation for amino acid
CY_3_R2: Cytosine pool 3, replicate 2 amino acid log2 fold-change
CY_3_MAD_2: Cytosine pool 3, replicate 2 codon log2 fold-change median absolute deviation for amino acid
FC_3: Measured log2 fold-change of variant in fragment 3, 5-FC
FC_3_R1: 5-FC pool 3, replicate 1 amino acid log2 fold-change
FC_3_MAD_1: 5-FC pool 3, replicate 1 codon log2 fold-change median absolute deviation for amino acid
FC_3_R2: 5-FC pool 3, replicate 2 amino acid log2 fold-change
FC_3_MAD_2: 5-FC pool 3, replicate 2 codon log2 fold-change median absolute deviation for amino acid
CY_1_to_2: Fragment 1 log2 fold-change adjusted to fragment 2 fold-change, cytosine 
CY_3_to_2: Fragment 3 log2 fold-change adjusted to fragment 2 fold-change, cytosine
FC_1_to_2: Fragment 1 log2 fold-change adjusted to fragment 2 fold-change, 5-FC
FC_3_to_2: Fragment 3 log2 fold-change adjusted to fragment 2 fold-change, 5-FC
CY_123: Averaged log2 fold-change across all fragments, cytosine
FC_123: Averaged log2 fold-change across all fragments, 5-FC
CY_score: DMS score (log2 fold-change scaled to synonymous mutants = 1 and nonsense mutants = 0), cytosine
FC_score: DMS score (log2 fold-change scaled to synonymous mutants = 0 and nonsense mutants = 1), 5-FC
FC_CY_score: DMS score (log2 fold-change scaled to synonymous mutants = 1 and nonsense mutants = 0), 5-FC + cytosine
interface: 1 = residue at interface, 0.75 = residue at interface rim, 0 = residue distal from interface
ddG_struc_HET: change in protein structure folding stability of mutant as predicted by foldX, mutant-wt dimer 
ddG_struc_HM: change in protein structure folding stability of mutant as predicted by foldX, mutant-mutant dimer
ddG_inter_HET: change in dimer interface stability of mutant as predicted by foldX, mutant-wt dimer
ddG_inter_HM: change in dimer interface stability of mutant as predicted by foldX, mutant-mutant dimer
r4s_score: evolutionary rate of residues based on a set of 215 orthologs, calulated using the rate4site software
RSA: relative solvent accessibility of residue
temp: temperature (aka B-factor, and indicator of rigidity in the structure) of residue
ortholog_aa_freq: frequency of variant in the multiple sequence alignment
dist_act_site: shortest distance between residue and a residue part of the active site
mut_type: mutation cluster (wt-like, nonsense-like, elbow aka front minima or other)
